# *Bacillus subtilis* YisK possesses oxaloacetate decarboxylase activity and exhibits Mbl-dependent localization

**DOI:** 10.1101/2023.06.26.546597

**Authors:** Tingfeng Guo, Anthony M. Sperber, Inna V. Krieger, Yi Duan, Veronica Chemelewski, James C. Sacchettini, Jennifer K. Herman

## Abstract

YisK is an uncharacterized protein in *Bacillus subtilis* previously shown to interact genetically with the elongasome protein Mbl. YisK overexpression leads to cell widening and lysis, phenotypes that are dependent on *mbl* and suppressed by *mbl* mutations. In the present work we characterize YisK’s localization, structure, and enzymatic activity. We show that YisK localizes in a punctate and/or punctate-helical pattern that depends on Mbl, and that YisK interacts directly with another elongasome protein, FtsE. YisK belongs to the fumarylacetoacetate hydrolase (FAH) superfamily and crystal structures revealed close structural similarity to two oxaloacetate (OAA) decarboxylases: human mitochondrial FAHD1 and *Corynebacterium glutamicum* Cg1458. We demonstrate that YisK can also catalyze the decarboxylation of OAA (K_m_ = 134 µM, K_cat_ = 31 min^-1^). A catalytic dead variant (YisK E148A, E150A) retains wild-type localization and still widens cells following overexpression, indicating these activities are not dependent on YisK catalysis. Conversely, a non-localizing variant (YisK E30A) retains wild-type enzymatic activity in vitro, but no longer widens cells following overexpression. Together these results suggest YisK may be subject to spatial regulation that depends on the cell envelope synthesis machinery.

**IMPORTANCE:** The elongasome is a protein complex that guides lengthwise growth in some bacteria. We previously showed that in *B. subtilis*, overexpression of an uncharacterized enzyme (YisK), perturbed function of the actin-like elongasome protein Mbl. Here we show that YisK exhibits Mbl-dependent localization and interacts directly with another component of the elongasome, FtsE. Through biochemical and structural characterization, we demonstrate that like it’s mitochondrial homolog FAHD1, YisK can catalyze the decarboxylation of the oxaloacetate to pyruvate and CO_2_. YisK is the first example of an enzyme implicated in central carbon metabolism with subcellular localization that depends on Mbl.

## INTRODUCTION

The bacterial cell envelope serves essential functions in nutrient uptake, secretion, energy production, morphogenesis, and reproduction and acts as a physical barrier against immune detection, antimicrobials, toxic metabolites, and bacteriophage. In addition, the envelope’s peptidoglycan (PG) layer helps protect the integrity of the cytoplasmic membrane. A highly tensile polymer, PG is elastic enough to allow for expansion and contraction during osmotic shifts, but strong enough to withstand multiple atmospheres of pressure (1–4). The envelope layers external to PG vary in composition by genus, species, and growth condition. In the Gram-positive bacterium *Bacillus subtilis*, the outer layers are composed of the anionic polymers teichoic and teichuronic acid that are anchored to the lipid bilayer, extracellular proteins, exopolysaccharides, and poly-γ-glutamic acid (5–7).

In many rod-shaped bacteria, including *B. subtilis*, the lengthwise incorporation of new PG during growth is governed by the elongasome, a membrane-associated, multiprotein complex that guides growth along the cell’s long axis (8). The elongasome is thought to be scaffolded by the actin-like protein MreB (9). MreB dynamically polymerizes and, along with its associated factors MreC and MreD (10), helps recruit enzymes that insert, modify, and hydrolyze PG (8, 11). Some bacteria possess multiple MreB paralogs and *B. subtilis* encodes three; MreB (encoded in the conserved *mreBCD* operon), Mbl, and MreBH. Each *B. subtilis* paralog can support maintenance of rod shape and viability if expressed at sufficient levels, consistent with functional redundancy (12). At the same time, the paralogs also possess some specialized activities (13–16). MreBH facilitates the function of LytE, a PG endopeptidase associated with envelope stress (14, 16, 17), whereas Mbl influences the activity of CwlO, a major PG endopeptidase utilized during cell growth (16). CwlO activity is also dependent on the conserved ABC transporter FtsEX (16, 17) and perturbed in the absence of two recently discovered cofactors, SweC and SweD (18).

In a prior study, we found evidence of a genetic interaction between Mbl and a putative enzyme, YisK (BSU10750). YisK overexpression causes cell-widening and lysis which can be suppressed by specific substitutions in Mbl or by deleting *mbl* in a background that relieves Mbl essentiality (Δ*ponA*, encoding PBP1A)(19). These results indicate YisK and Mbl interact at least genetically and are consistent with YisK overexpression disrupting Mbl function.

YisK is in the SigH regulon (20), induced during the early stages of sporulation (19), and classified as a general starvation protein (21). Based on transcriptomic and proteomic analysis, YisK is also expressed in a variety of other conditions including exponential growth in M9 minimal medium (22–24). In M9 containing glucose and malate as carbon sources, YisK is estimated to be present at ∼1,000 copies per cell across exponential, transition, and stationary phases of growth (23). Thus, despite being in the SigH regulon, YisK is clearly expressed in growth contexts outside of stationary phase, sporulation, and general starvation.

Based on sequence conservation, YisK belongs to the fumarylacetoacetate hydrolase (FAH) superfamily, a class of catabolic enzymes that catalyze diverse reactions, including dehydrations, decarboxylations, and isomerizations (25). FAH proteins are found in all domains of life and share a common mixed β-sandwich-like fold (26). The catalytic pocket coordinates a divalent metal ion, usually Mg^2+^, Mn^2+^ or Ca^2+^, that is essential for enzymatic activity (27, 28). No FAH proteins are known to be involved in cell envelope synthesis, so possible reasons for the epistatic interaction between YisK and Mbl could not be inferred in the prior study. In the present study, we show that YisK exhibits Mbl-dependent localization and interacts directly with FtsE, characterize YisK structurally, and demonstrate that YisK can catalyze the decarboxylation of OAA to pyruvate and CO_2_.

## RESULTS

### YisK protein is expressed in sporulation by resuspension medium, CH, and LB

*yisK* is transcriptionally upregulated in “sporulation by resuspension” medium or during growth in a rich medium permissive for sporulation, CH (19). To assess accumulation of YisK protein under these growth conditions, we performed western blot analysis using a polyclonal antibody specific for YisK (Fig S1). When cells were grown in sporulation by resuspension medium, YisK reached maximal levels by 60 min (Fig 1A)(19); these data correlate well with the appearance of GFP signal from a P*_yisK_*-GFP reporter in the same growth conditions (19). In the amino acid-based medium CH, YisK was detectable during exponential growth (OD_600_ 0.3 and 0.6) and reached maximal levels between OD_600_ readings of 3.6 and 4.6 (Fig 1B). The higher OD_600_ range corresponds to when some cells in the population start to enter the sporulation program in CH (19). We also examined YisK levels in LB supplemented with 10.0 mM MgCl_2_ (LB-Mg), which permits growth to higher cell densities (29). In LB-Mg, YisK was detectable during early exponential growth, but increased to higher levels at transition state and beyond (Fig 1C).

**Figure 1.**
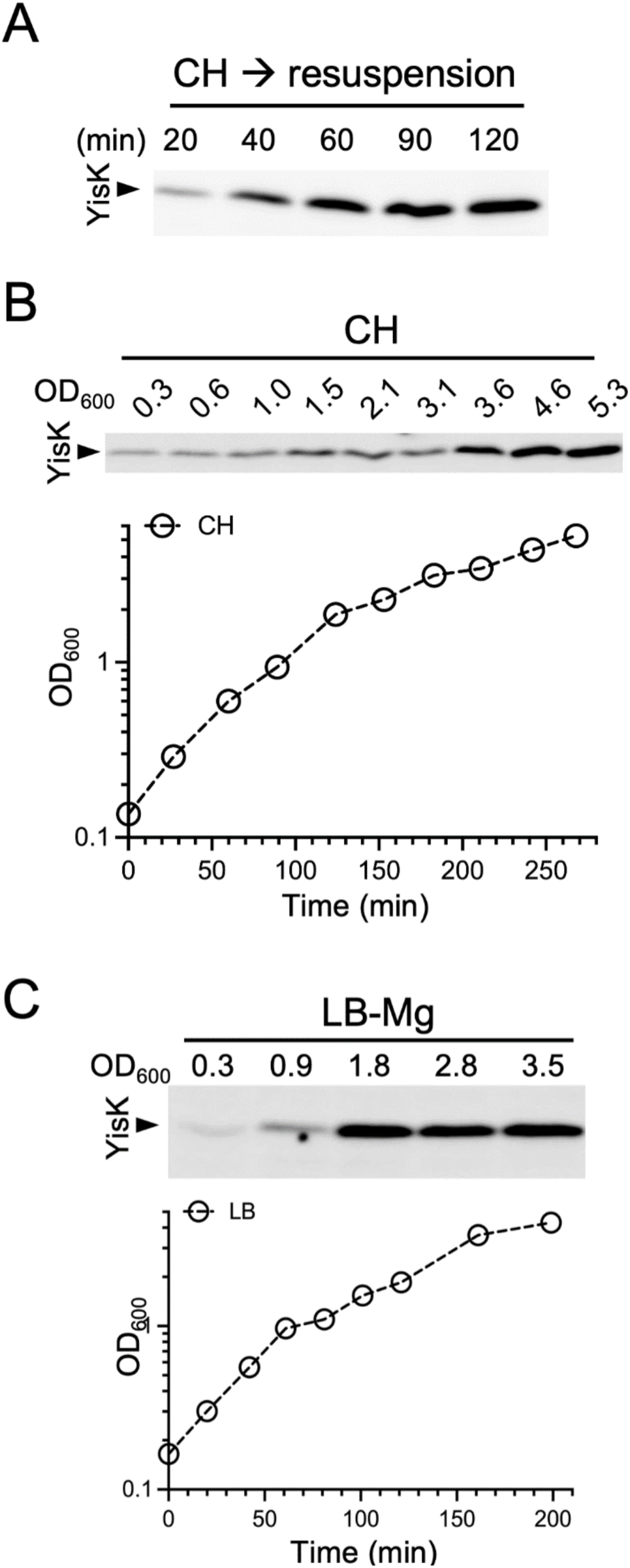
Analysis of YisK protein levels during growth. (A) Representative western blot of samples collected during growth in sporulation by resuspension medium. Representative western blots and growth curves from cells grown at 37°C in (B) CH and (C) LB-Mg.

### YisK interacts directly with FtsE

In a prior study, YisK was shown to interact with Mbl genetically (19). We used bacterial two-hybrid (B2H) analysis to assess if a direct YisK-Mbl interaction could be detected. Although one of the pairwise combinations was positive (indicated by development of blue color), the signal was indistinguishable from one of the negative controls (Fig 2A). We also tested for interaction between YisK and the proteins FtsE and FtsX, as they are proposed to act in the same genetic pathway as Mbl (16). In this case, a pairwise interaction between YisK and the ATPase FtsE was readily detectable before the negative control turned blue (Fig 2B). FtsE and the transmembrane protein FtsX form a complex (30). When FtsX was used as bait, no interaction was detected; however, if untagged FtsE was co-expressed with FtsX as bait, a pairwise interaction was again observed (Fig 2B). Together these results suggest that YisK can interact directly with FtsE and the FtsEX complex.

**Figure 2.**
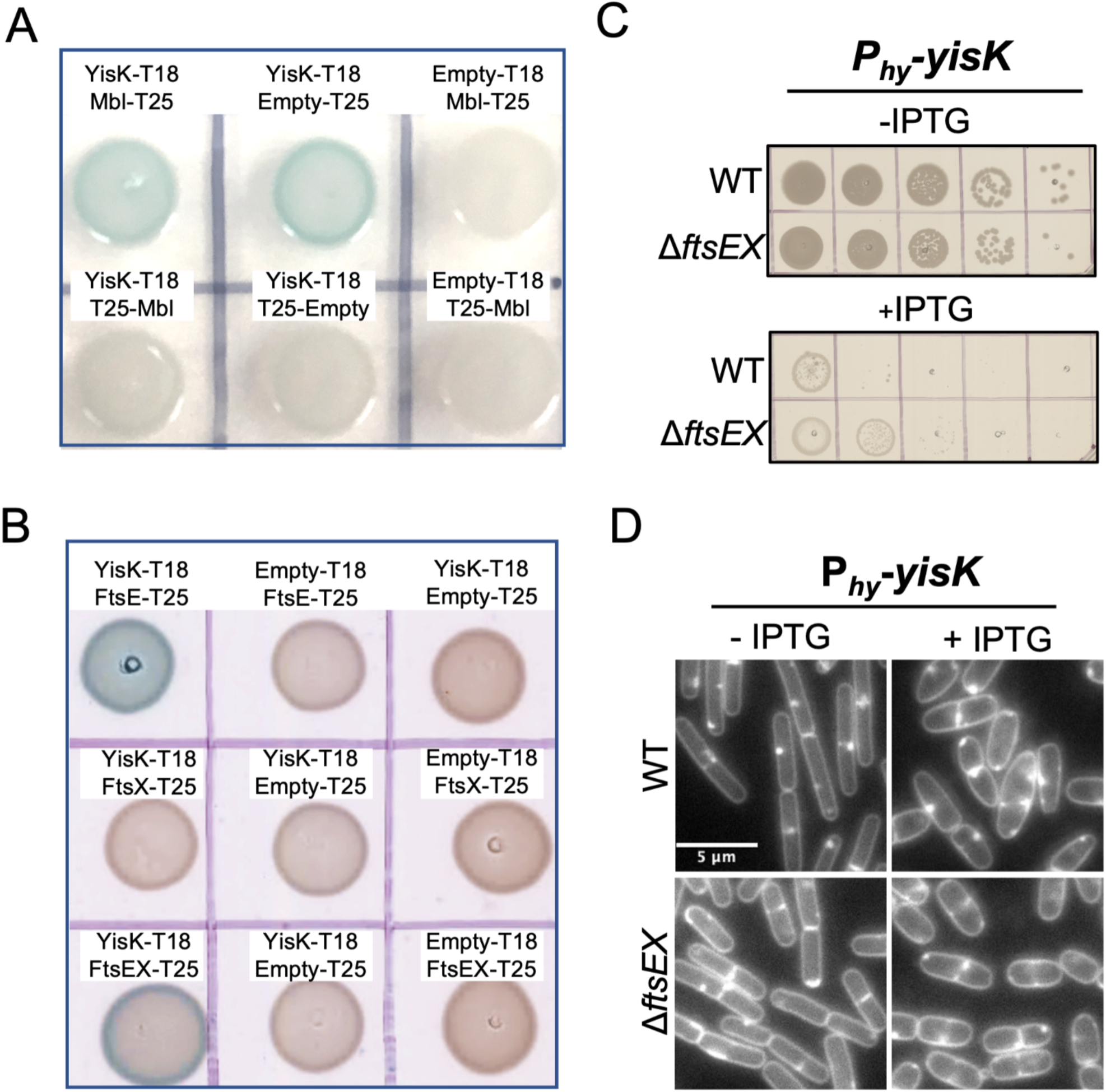
Assessing YisK interactions. (A and B) Bacterial two-hybrid analysis. (C) YisK overexpression assay. Cells harboring *P_hy_-yisK* were spotted on LB without or with 1.0 mM IPTG and incubated overnight at 37°C before imaging. (D) Representative micrographs of cells without and with YisK induction. Cells were grown at 37°C in LB-Mg and induced with 1.0 mM IPTG for 60 min. Membranes were stained with TMA. Images are scaled identically.

To test if FtsEX was required for YisK’s previously characterized overexpression phenotypes, we introduced Δ*ftsEX* into the P*_hy_*-*yisK* overexpression strain. The 11*ftsEX* strain was still inhibited for growth following YisK overexpression on LB plates (Fig 2C) and still exhibited cell widening in liquid LB culture (Fig 2E), indicating that FtsEX is not required for YisK to perturb Mbl function.

### YisK forms Mbl-dependent puncta in the cell

Our data suggest YisK interacts with two components of the *B. subtilis* elongasome: Mbl, based on genetic data (19) and FtsE based on direct interaction (Fig 2B). FtsE and Mbl both lack transmembrane domains but exhibit subcellular localization (31–34). To determine if YisK also localizes within the cell, we generated a fluorescent GFP “sandwich” fusion (YisK_SW_-GFP) by introducing GFP into a non-conserved loop in YisK between residues E111 and A112. Cells overexpressing YisK_SW_-GFP from an IPTG-inducible promoter (P*_hy_*) were inhibited for growth on LB plates (Fig 3A) and became wide in LB (Fig 3B), consistent with the previously reported YisK overexpression phenotypes (19). These results indicate that YisK_SW_-GFP retains the YisK-associated phenotypes.

**Figure 3:**
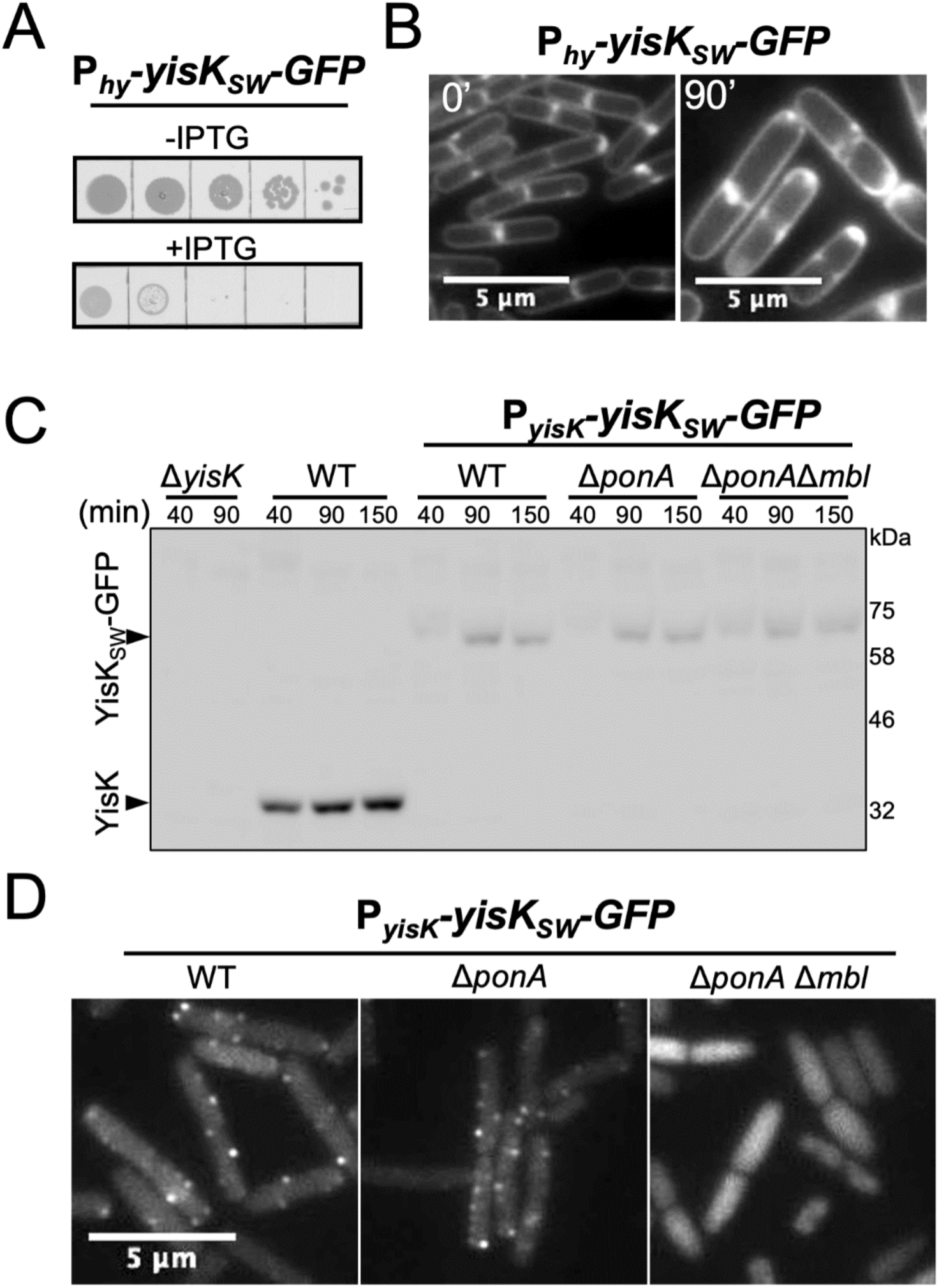
Assays assessing the functionality, expression, and localization of YisK_SW_-GFP. (A) Cells harboring *P_hy_-yisK_SW_-GFP* were spotted on LB without or with 1.0 mM IPTG and incubated at 37°C overnight before imaging. (B) Representative micrograph of cells harboring *P_hy_-yisK_SW_-GFP* following 90 min induction with 1.0 mM IPTG. Membranes were stained with TMA. Images are the same magnification. (C) Western blot analysis of YisK and YisK_SW_-GFP expression during growth in sporulation by resuspension medium using ⍺-YisK serum. (D) Representative micrographs of cells harboring P*_yisK_-yisK_SW_-GFP* following 40 min growth in sporulation by resuspension medium. Images are the same magnification and scaled identically in the GFP channel.

Next, we used allelic exchange to replace wild-type *yisK* with *yisK_SW_-GFP* at the native locus. In this strain, *yisK_SW_-GFP* is under the control of the native *yisK* promoter (P*_yisK_*) and no other copy of *yisK* is present in the cell. Having already established that untagged YisK is expressed when cells are resuspended in a sporulation medium (Fig 1A)(19), we first monitored YisK_SW_-GFP expression and stability at several timepoints after resuspension using western blot analysis (Fig 3C). Similar to untagged YisK, YisK_SW_-GFP protein was observable at 40 min and increased in abundance between 40 and 90 min (Fig 3C). The reduced signal from YisK_SW_-GFP in comparison to the untagged YisK may be due to occlusion of anti-YisK epitopes by the internal GFP tag and/or differences in efficiency of transfer of the different proteins to the membrane. YisK_SW_-GFP was detected as a single band using both anti-YisK and anti-GFP antibodies (Fig 3C and Fig S2B), indicating the fusion is not proteolyzed.

Having established that the YisK_SW_-GFP retains phenotypes and is expressed as a stable fusion from the native locus in resuspension medium, we next monitored YisK_SW_-GFP localization using epifluorescent microscopy. Following growth in resuspension medium, YisK_SW_-GFP localized both diffusely and in punctate or punctate-helical arrangements (Fig 3D and Fig S2A). Similar localization was also observed in CH and LB (Fig S2C and S2E).

Based on the fact that YisK overexpression perturbs Mbl function and that Mbl is required for YisK to widen cells (19), we hypothesized that YisK might depend on Mbl for localization. To create a strain background permissive for deletion of *mbl* (11), deleted *ponA* in the P*_yisK_-yisK-GFP* harboring strain before introducting the *mbl* deletion. Next we compared YisK_SW_*-*GFP localization in the Δ*ponA* and Δ*ponA* Δ*mbl* backgrounds to wildtype. As can be seen in Fig 3D, YisK_SW_*-*GFP, still displayed punctate localization in the Δ*ponA* cells. In contrast, YisK_SW_*-*GFP was completely diffuse in the Δ*ponA* Δ*mbl* background (Fig 3D). YisK_SW_*-*GFP was expressed equivalently and migrated at the same apparent molecular weight in all three backgrounds, indicating the diffuse signal is not due to clipping/release of GFP or a change in expression levels at the population level (Fig 3C). From these data we conclude that YisK exhibits diffuse and punctate localization, and that the punctate localization depends on Mbl.

### Identification of molecules that stabilize YisK

To gain further insight into YisK function, we sought to characterize YisK biochemically. Following overexpression and purification of a His-tagged fusion of YisK (Fig 4A), we noted that concentrated YisK solution (∼100.0 µM) was tinted slightly pink (Fig 4B). Since characterized FAH proteins typically coordinate a divalent cation in a conserved active site (25), Mn^2+^ solutions are pink, and the pink color was absent when two of YisK’s three metal-coordinating residues were substituted with alanine (YisK E148A, E150A)(Fig 4B), we hypothesized that YisK most likely coordinated Mn^2+^. To test, we monitored the thermostability of YisK without and with buffer containing either MnCl_2_ or MgCl_2_. Notably, YisK’s melting temperature increased by 8°C in the presence of 5.0 mM MnCl_2_ but differed insignificantly with 5.0 mM MgCl_2_ (Fig 4C). Even though purified YisK was pink compared to the E148A E150A variant, the fact that the additional metal was stabilizing suggests the majority of purified protein lacked a metal cofactor. To estimate the apparent binding affinity (K_a_) of YisK for Mn^2+^ and Mg^2+^, we monitored YisK’s thermostability across a gradient of metal concentrations (Fig 4D and 4E). From the curves, the apparent K_a_ of YisK was estimated to be approximately 30 µM for Mn^2+^ and 60 mM for Mg^2+^. Collectively, these data suggest YisK likely utilizes Mn^2+^ over Mg^2+^ as a cofactor.

**Figure 4.**
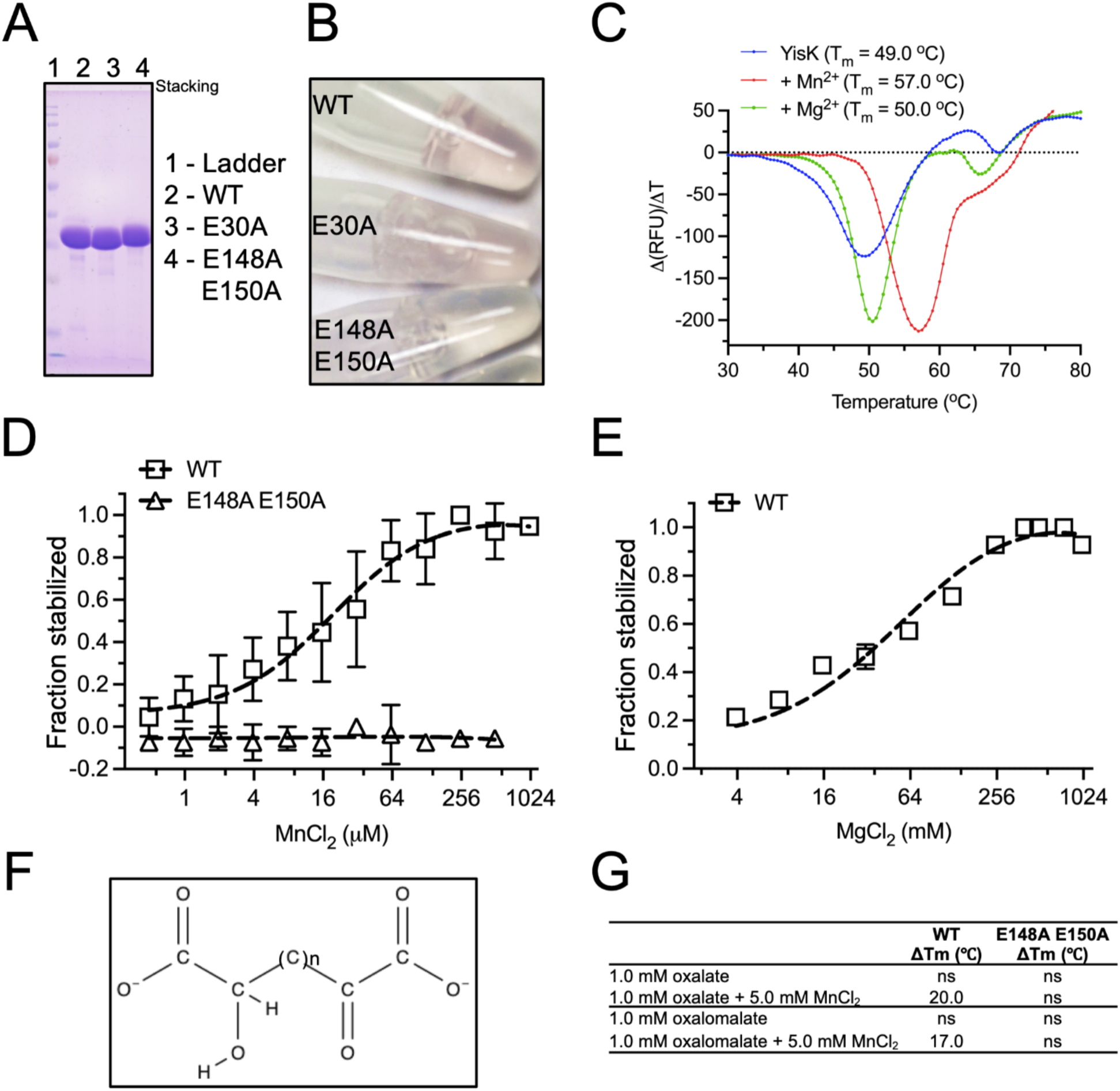
Affinity of YisK for Mn^2+^ as a cofactor. (A) Colloidal Coomassie blue staining of gel of purified proteins used in assays. (B) Purified YisK and YisK variants concentrated to ∼100 µM. (C) DSF with 10.0 µM YisK (WT) and buffer containing either 5.0 mM MnCl_2_ or 5.0 mM MgCl_2_. (C and D) DSF assay with gradient of (C) MnCl_2_ or (D) MgCl_2_. The fraction of YisK stabilized was calculated according to formula: ΔT_m_/(T_mMax_-T_m0_). (E) Structural elements conserved amongst compounds that increase YisK’s melting temperature. n = 0 to 2 carbons. (G) DSF assay containing 10.0 µM of the indicated YisK protein. A thermal shift of ≤ 2.0°C was not considered significant (ns).

The first YisK crystals (more below) were obtained in the presence of Tacsimate, a mixture of organic acids including L-tartrate. As a positive electron density consistent with tartrate was observed coordinating metal in the active site, we hypothesized that YisK’s substrate might possess properties similar to the molecules constituting Tacsimate. Using differential scanning fluorimetry (DSF), we found tartrate markedly increased YisK’s thermostability (Table S1). By screening a panel of structurally related compounds, we identified additional molecules that increased YisK’s melting temperature as well as those that did not (Fig S4 and Table S1). Oxalomalate, D,L-4-hydroxy-2-ketoglutarate (HOGA), and oxalate all dramatically stabilized YisK, increasing the melting temperature (T_m_) by more than 15°C (Fig S4). Glyoxylate, OAA, and pyruvate were not stabilizing in isolation, but the combinations of glyoxylate and either OAA or pyruvate increased YisK’s melting temperature significantly (Fig S4). Overall, molecules that stabilized YisK had properties similar to the molecule shown in Fig 4F.

Based on characterized FAH family proteins, YisK’s ligand/substrate would bind in the active site by coordinating the Mn^2+^ cofactor. To test if Mn^2+^ was required for compounds to stabilize YisK, we repeated the DSF experiments for oxalate and oxalomalate in buffer without and with addition of metal; for both compounds, Mn^2+^ was required for stabilization (Fig 4G). Moreover, the YisK E148A E150A variant (which cannot coordinate Mn^2+^) was not stabilized by oxalate or oxalomalate even when Mn^2+^ was included in the buffer (Fig 4G). These results are consistent with the ligands coordinating the metal cofactor in the active site as observed in published FAH crystal structures (25, 28).

### Structural characterization of YisK

To investigate YisK’s potential catalytic activity and address the possible functional implications for the interaction between YisK and Mbl, we characterized YisK structurally. C-terminally His-tagged YisK was overexpressed and purified, and following screening, initial crystals hits were obtained in the presence of tacsimate. To optimize the conditions and achieve uniformly liganded protein, we complexed YisK with Mn^2+^ and a subset of the ligands identified in DSF experiments, and repeated crystallization condition screening. The newly identified PEG-based crystallization condition was used for all structures reported in this paper. The structure of YisK in complex with oxalate (a potent inhibitor of the FAH proteins FADH1 and Cg 1458)(27, 35) and Mn^2+^ was solved at a resolution of 2.4 Å by molecular replacement using a homology model created by Phyre2 (36). YisK crystalized as homodimer with one dimer in the asymmetric unit of a *P2_1_2_1_2_1_* crystal lattice. Each subunit is composed of a large and small domain (Fig 5A and 5B). The fold of the larger domain is highly conserved among FAH proteins and harbors a triad of negatively charged amino acids (E148, E150, and D179 in YisK) that coordinate a divalent cation typically required for catalysis (25). The smaller domain is variable in size and sequence among the FAH superfamily and often corresponds to a domain of unknown function (DUF2437/PF10370).

**Figure 5.**
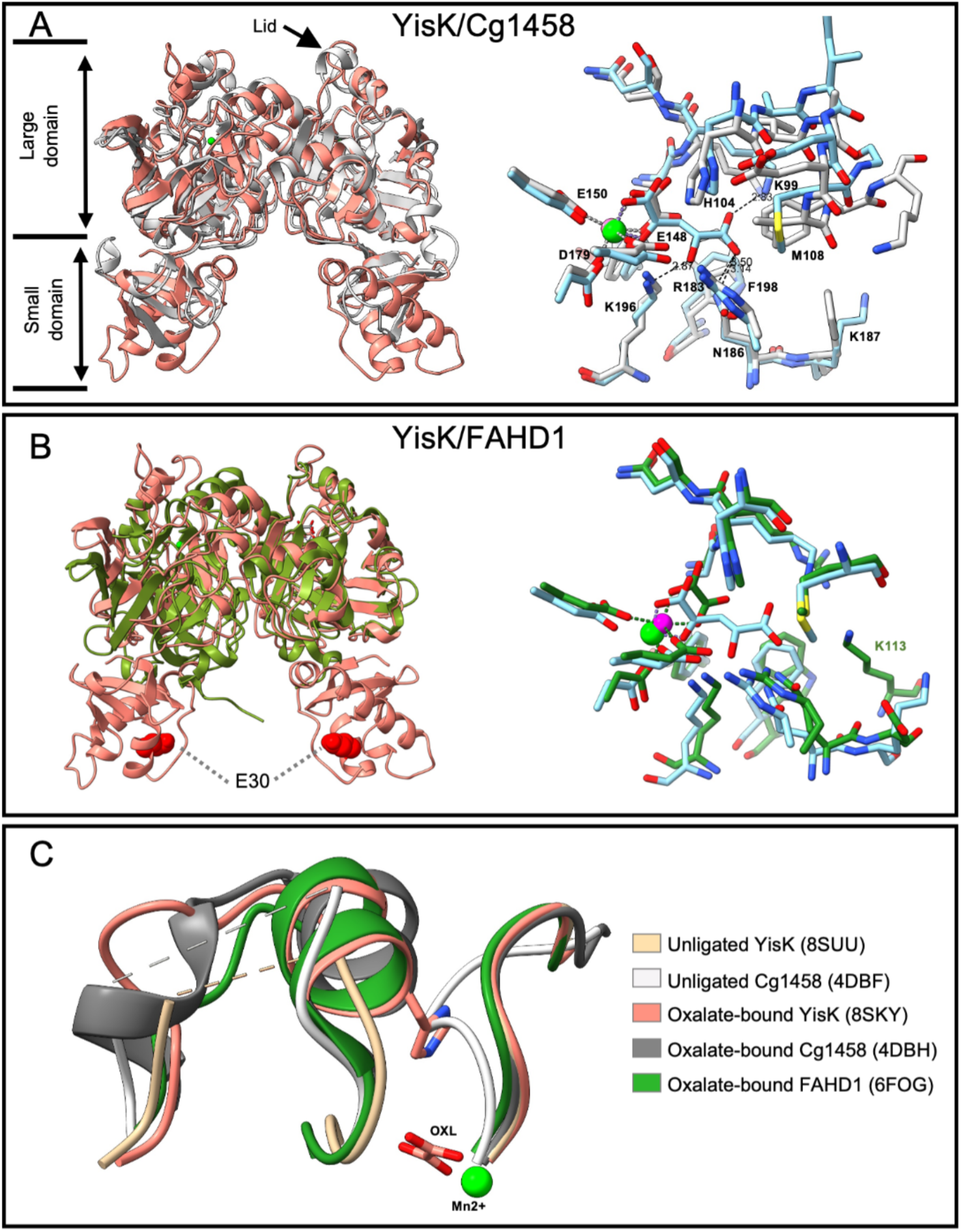
Comparison of YisK structure with the oxaloacetate decarboxylases Cg1458 and FAHD1. (A and B, left) Cartoon representation of YisK dimer (salmon) superimposed with (A) Cg1458 (light grey) and (B) FAHD1 (green). Residue E30 (red spheres) is important for YisK localization and part of the small domain absent in FAHD1. (A and B, right) Active site comparison between HOGA--bound YisK (8SUT, light blue carbons) with (A) oxalate-bound Cg1458 (4DBH, light grey carbons) and (B) FAHD1 (6FOG, green carbons). The models are colored by heteroatom. Only the HOGA ligand is shown. Mn^2+^ (green) and Mg^2+^ (magenta) are shown as spheres. (C) Conformational changes between the liganded and unliganded FAH enzyme structures. Unliganded YisK (beige) and Cg1458 (white) with the lids disordered and open, respectively. The oxalate-bound forms of YisK (salmon), Cg1458 (dark grey), and FAHD1 (green) with the lid closed. Disordered segments are depicted as dashed lines. Oxalate (OXL) and YisK’s His104 side chain are shown as stick models. The Mn^2+^ (green sphere) is from the YisK structure (8SKY). There is a positional clash between loop 238-242 of the unliganded Cg1458 (white) and His104 of oxalate-bound YisK.

A structural similarity search using VAST (37) indicated YisK shares the highest homology with FAH from mouse (1HYO), 5-carboxymethyl-2-hydroxymuconate delta isomerase from *Mycobacterium smegmatis* (4PFZ), and HpcE from *Thermus thermophilus* (1WZO), with vast scores of 39.4, 39.3, and 39 respectively. The root mean-square deviation (RMSD) calculated for Cα of the YisK large domain superposed with these hits varies between 1.5 to 2.8 Å. The next eighteen VAST hits had scores from 23.4 to 38.1, with the similarity driven by the high sequence conservation of the FAH catalytic fold.

As FAH domain proteins are known to sustain an array of catalytic functions (*eg*. hydrolase, decarboxylase, isomerase), comparing the overall fold of YisK to known FAH family proteins is not very informative for revealing potential catalytic and substrate specificity. Therefore, we instead examined the composition of the active site residues and compared those to homologs, considering not only sequence, but also 3D positional alignments. Highest sequence similarity was discovered with two OAA decarboxylases; human mitochonrial FAHD1 with 43% identity over 59% of covered sequence and *Corynebacterium glutamicum* Cg1458 with 36% identity over 74% of covered sequence (Fig S3A); the RMSDs over the aligned main chain pairs were 0.96 and 0.85 Å, respectively. The overall structures of the large domains are similar, differing only in the small domain (Fig 5A and 5B), particularly for FAHD1 in which the small domain is essentially absent (Fig 5B). In the structures, YisK binds Mn^2+^ while both Cg1458 and FAHD1 bind Mg^2+^. However, Cg1458 possessed highest activity in vitro with Mn^2+^-containing buffer (27). The overlay of the ligand-bound FAHD1, Cg1458, and YisK structures shows that all of the residues proposed to coordinate the metal and substrate and participate in the catalysis are conserved; this includes the histidine glutamate-water triad (H104, E148) common among FAH family proteins with carbon-carbon bond cleaving activities (Fig S3A)(27). Only a few differences occurred in close proximity to the ligand; the K99/F192 equivalent positions of YisK are replaced with R25/W119 in FAHD1 and R68/W160 in Cg1458. In addition, in place of YisK M108 (of the lid)(Fig S3A), Cg1458 has V77 followed by F78, the latter of which blocks Cg1458’s active site due to its bulky sidechain. Additionally, FAHD1 has a short sequence insertioncompared to YisK (Fig S3A), that contributes K113 to the closing of the active site (Fig 5D). These structural observations led us to hypothesize that YisK can decarboxylate OAA, which was confirmed by a coupled enzyme assay (see below). Given that the differences in YisK make the active site less constricted compared to the other OAA decarboxylases, it is possible YisK can also accommodate larger substrates.

Prompted by the thermal shift analysis which identified other potential ligands, we successfully co-crystallized YisK with oxalomalate. We noted that the difference electron density for the ligand coordinated to Mn^2+^ was consistent with the product of oxalomalate decarboxylation, HOGA (Fig S3B), suggesting that YisK can decarboxylate other substrates; although we think it is unlikely based on the crystallization conditions, we do not rule out the possibility that the oxalomalate’s decarboxylation could have occurred spontaneously. Regardless, this structure illustrates how a ligand larger than OAA can be accommodated in the YisK active site: distal to the Mn^2+^, the keto-acid moiety is stabilized by contacts with N186, R183 and K99 (Fig 5A). The overall protein structure does not differ significantly between oxalate and 4-hydroxy-2-oxoglutarate bound structures (Fig S3C).

Interestingly, analysis of the electron density maps of YisK crystals formed in the presence of OAA revealed an unliganded active site, with only a Mn^2+^ ion coordinated by the protein side chains and water molecules. From this we think it is likely that OAA bound to the active site was decarboxylated during crystallization, and that the resultant pyruvate lacked sufficient affinity to stay ordered. Consistent with this interpretation, a density for pyruvate was also not observed following crystallization in the presence of high concentrations pyruvate. The crystals set with OAA produced an effectively “apo” structure of YisK. The open lid conformation (Fig 5C) signified by the loss of ordered structure for the residues 105-114 on chain A, and 103-113 on chain B, is similar to what was shown for apo Cg1458 and FAHD1 (27, 35). Notably, this transition starts at the catalytic H104, which is still visible on chain A, but is a part of disordered segment on chain B. From this we infer that ligand binding is the most likely driver of the conformation change resulting in the lid closure and ordering of the 103-114 segment. In contrast to the true apo structure of Cg1458, we do not observe in our apparent apo YisK structure the transition equivalent to Cg1458 loop 238-242, which takes the place of the lid residues’ side chains, including the catalytic histidine of the liganded structure conformation (Fig 5C). Although this segment is relatively conserved by sequence (SPAGT in Cg1458 versus TPSGV in YisK), in YisK it is followed by a seven amino acid long insertion (Fig S3A) that may affect the mobility of the neighboring loop. Like YisK, the unliganded FAHD1 structure (6FOH), does not show the movement of analogous loop. Hence, loop 238-242 blocking the active site conformation state may be unique to Cg1458; it is also possible that the way we obtained unliganded structure of YisK – through dissociation of the reaction product – did not allow for this conformation.

### YisK protein can catalyze the decarboxylation of OAA to pyruvate and CO_2_

Although we identified several chemical features YisK’s substrate(s) is/are likely to possess (Fig 4F), the number of potential substrates remained large. Comparison of the YisK structure with the crystal structures of other FAH family proteins revealed strong active site similarity with two OAA decarboxylases, FAHD1 and Cg1458 (Fig 5A and 5B). To test if YisK also has OAA decarboxylase activity, we performed a coupled lactate dehydrogenase (LDH) assay, monitoring the conversion of pyruvate to lactate spectrophotometrically by following the oxidation of NADH to NAD+ at 340 nm. OAA spontaneously decarboxylates in solution (Fig 6A)(38, 39), so the apparent kinetic constants for YisK-catalyzed OAA decarboxylation were determined after subtracting the rate of spontaneous decarboxylation (Fig 6A) that occurred in the absence of YisK. The YisK-catalyzed reaction rate was plotted (Fig 6B) and used to derive the following apparent kinetic constants: *k*_cat_: 0.52 s^−1^ ± 0.02 s^−1^, *K*_m_: 134 µM ± 22 µM, and *k*_cat_/*K*_m_: 3.8 × 10^3^ M^−1^ s^−1^.

**Figure 6.**
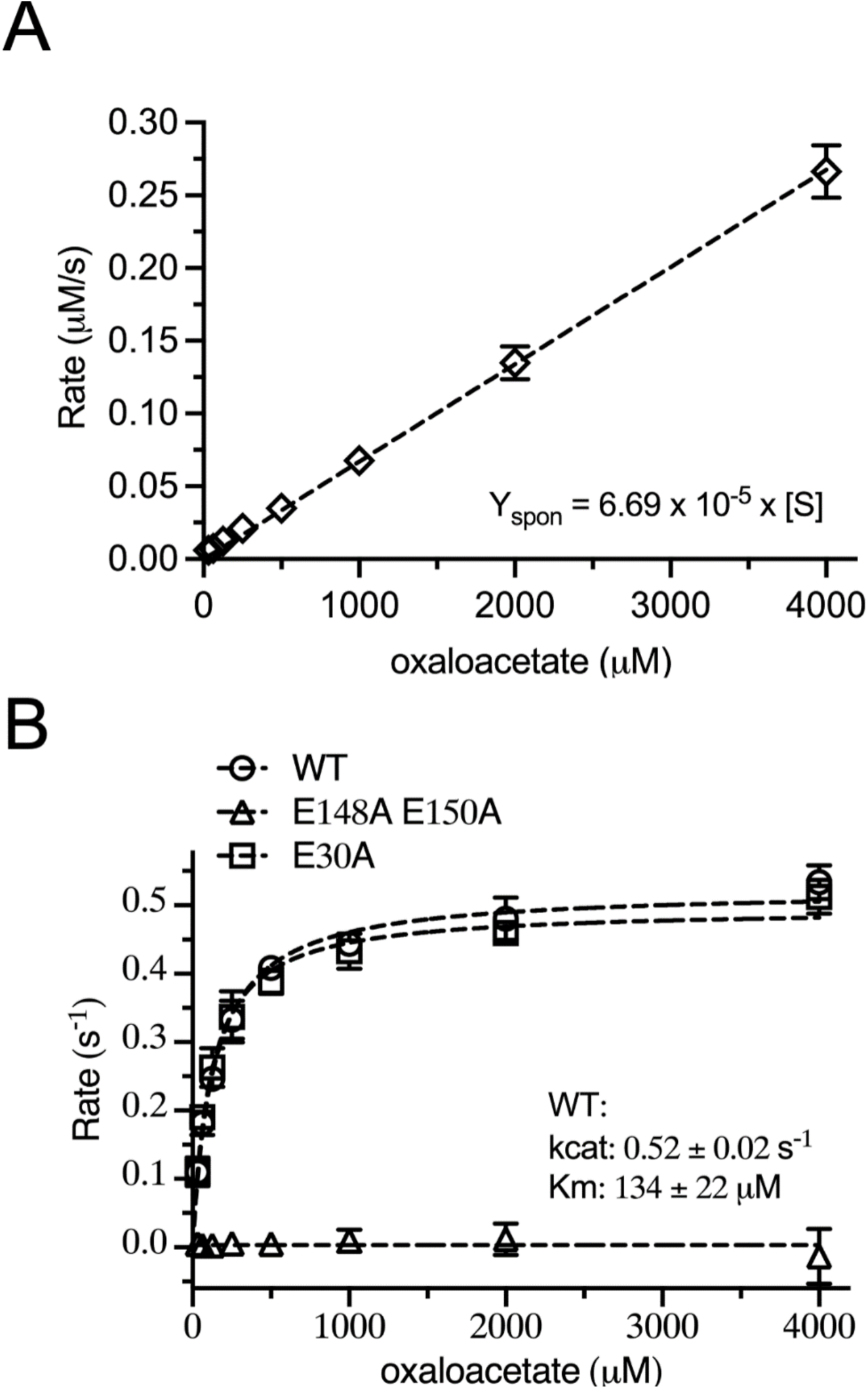
Coupled assay to detect oxaloacetate decarboxylase activity. (A) spontaneous decarboxylation of oxaloacetate at 25°C in buffer containing 0.3 mM NADH, 1.0 unit lactate dehydrogenase, and the indicated concentration of oxaloacetate. The dashed line represents the linear fits for the data. (B) Michaelis-Menten plot for decarboxylation of oxaloacetate catalyzed by YisK and variants after subtracting spontaneous decarboxylation. Reactions were carried out at 25°C in buffer containing 0.3 mM NADH, 1.0 unit lactate dehydrogenase, and the indicated concentration of oxaloacetate, and initiated by the addition of 0.5 µM YisK. The dashed line represents the fit of the data to *v*/E_t_=*k*_cat_(S)/(*K*_m_+S).

As a control, the assay was also performed with the YisK E148A E150A variant that is predicted to be catalytically dead due to loss of the Mn^2+^ cofactor. As expected, YisK E148A E150A had no detectable activity (Fig 6B). In contrast, introducing a substitution in the small domain distal to the active site (YisK E30A)(Fig 5B) had no effect on catalytic activity (Fig 6B). These data support the idea that YisK-dependent OAA decarboxylation requires the metal co-factor and exclude the possibility that the observed activity is due to either spontaneous decarboxylation or another enzyme contaminating the protein preparation. We conclude that YisK possesses OAA decarboxylase activity.

### Catalytically inactivated YisK can still widen cells and exhibits wild-type localization

To test whether YisK’s enzymatic activity is required for the overexpression phenotypes, we placed YisK E148A E150A under the control an IPTG inducible promoter. Following induction, the catalytic dead variant still prevented growth on LB plates and led to loss of control over cell dimensions in liquid LB (Fig 7A). Moreover, a GFP fused version of YisK E148A E150A still exhibited punctate localization (Fig 7B). These results indicate that YisK’s localization and interaction with the elongasome does not depend on its catalytic activity. In addition, since YisK E148 E150A is unlikely to efficiently bind ligands (Fig 4G), the results also suggest that YisK’s punctate localization is not dependent on ligand binding.

**Figure 7.**
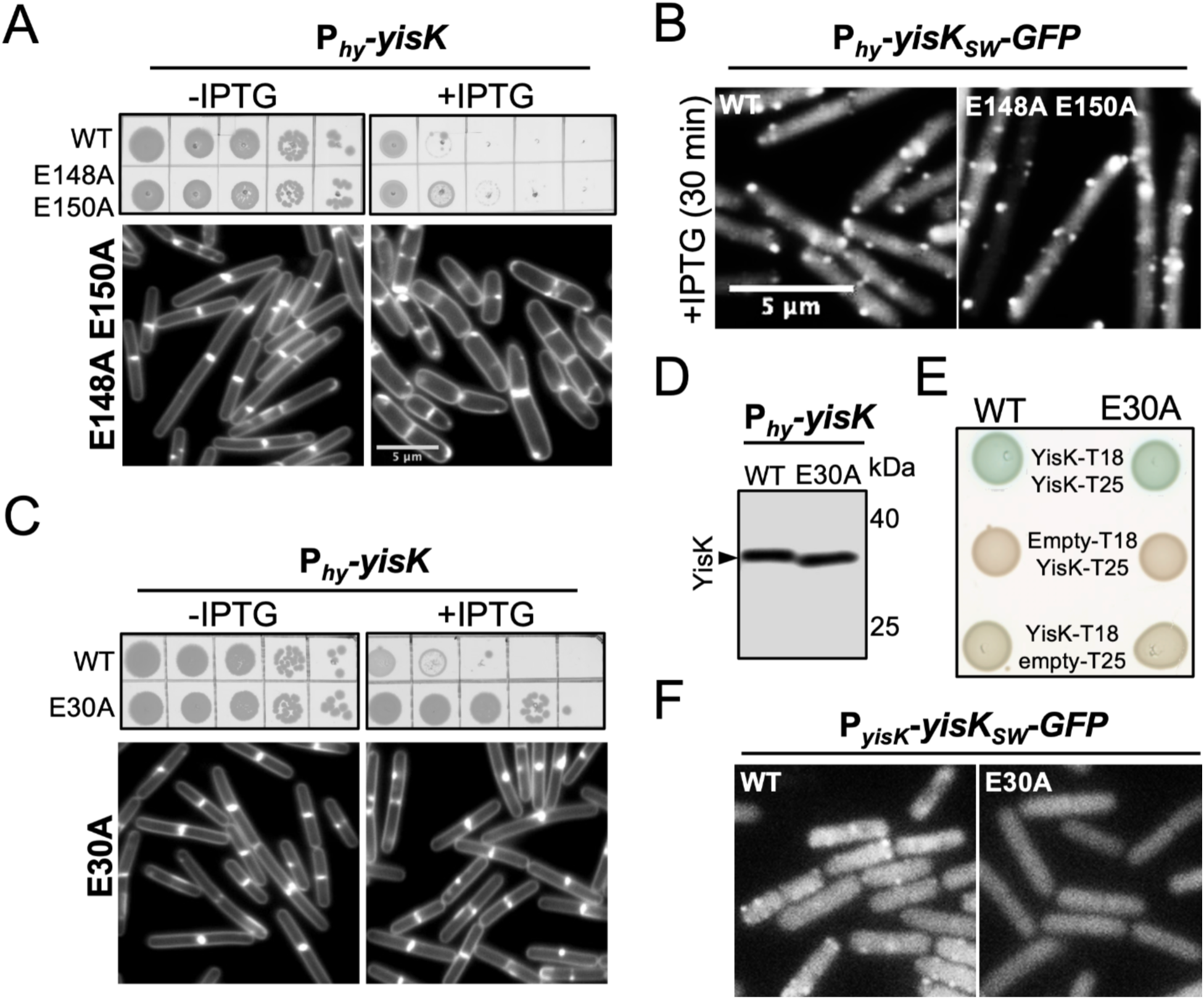
YisK variant overexpression phenotypes and localization. (A and C) Growth of cells harboring indicated construct after overnight incubation (top) and following 90 min overexpression in LB-Mg (bottom). Membranes are stained with TMA and shown at the same magnification. (B) Localization of YisK_sw_-GFP in the indicated background following 30 min induction in LB-Mg. (D) Western blot analysis with anti YisK antibody of samples collected from cells harboring wild-type *P_hy_-yisK* and *P_hy_-yisK* E30A following 60 min induction. (E) Bacterial 2-hybrid analysis for YisK self-interaction. (F) Representative micrographs of YisK_SW_-GFP localization following 40 mins growth in sporulation by resuspension medium (native promoter). Images are the same magnification and scaled identically in the GFP channel.

### YisK E30A no longer kills or widens cells and localizes diffusely

Prior to solving the crystal structure, we identified a small cluster of charged residues (EKK) in the small domain of YisK that we guessed might be surface exposed. We hypothesized that this region of YisK might be important for protein-protein interaction and/or localization and generated a YisK E30A substitution to assay for functionality. In contrast to wild-type YisK, overexpression of YisK E30A did not prevent growth or cause cell widening (Fig 7C). YisK E30A expression was indistinguishable from wild-type YisK by western blot analysis (Fig 7D), suggesting that the loss-of-function phenotypes were not attributable to a problem with protein expression. Moreover, YisK E30A maintained wild-type self-interaction in a bacterial two-hybrid assay (Fig 7E) and exhibited wild-type enzymatic activity in vitro (Fig 6B), both consistent with proper folding. Strikingly, YisK_SW_-GFP E30A expressed from YisK’s native promoter exhibited only diffuse localization (Fig 7F). These results suggest that the small domain is important for YisK’s Mbal-dependent punctate localization and ability to perturb Mbl function.

## DISCUSSION

### Is OAA YisK’s preferred substrate?

The YisK-catalyzed turnover of OAA was 31 min^−1^, rather slow for a typical enzyme. We can envision several possible explanations for the modest turnover. First, it is entirely possible that OAA is not a preferred substrate. If so, additional testing may eventually lead to better candidates. Second, assuming an intracellular OAA concentration of 0.5 µM (the estimate for *E.coli*)(40), YisK-catalyzed decarboxylation would be around 30 times faster than the spontaneous rate observed in our reaction buffer (Fig. 6A). This degree of catalytic enhancement could conceivably be sufficient for the cell’s needs, especially considering that YisK is a relatively abundant protein (21, 23). Third, YisK’s punctate localization suggests the enzyme may reach high concentrations locally, which could be important for in vivo activity; relatedly, the in vitro assay may lack one or more factors that enhance YisK activity, such as an interaction partner. Finally, YisK could function more as a metabolite sensor than an enzyme. For example, upon ligand binding (or release), YisK could adjust some activity related to elongasome function. YisK likely does not require a ligand to localize, as the variant with low affinity for ligands (YisK E148A E150A)(Fig 5G) and no detectable enzymatic activity (Fig 6B), can still form puncta and widen cells (Fig 7A and 7B); at the same time, our data do not rule out the possibility that ligand binding could cause YisK to delocalize and/or change its function.

OAA is an important metabolic intermediate that along with phosphoenolpyruvate and pyruvate forms the central node connecting glycolysis, gluconeogenesis, and the TCA cycle (41, 42). It is also the precursor to aspartate, the aspartate-derived amino acids asparagine, methionine, threonine, isoleucine, and lysine, and in bacteria, the PG precursor mesodiaminopimelic acid. OAA is also a potent inhibitor of succinate dehydrogenase (43–47) and control of OAA levels is proposed to be integral to the TCA cycle regulation and electron transport chain function (48, 49). Inhibition of succinate dehydrogenase by OAA leads to a drop in proton motive force and ATP synthesis, but also reduces oxidative stress (50). Depletion of mitochondrial FAHD1 in human epithelial cell lines disrupts electron transport chain function and triggers senescence, and it has been proposed that FAHD1 may play a role in TCA cycle regulation (51). If YisK functions as an OAA decarboxylase in vivo, then it could also buffer the cell against succinate dehydrogenase inhibition. Notably, YisK protein accumulates to highest levels when carbon or nitrogen sources become limited (23, 52, 53) and after cells exit exponential growth (Fig 1 and Fig S1). As this is also when the punctate localization becomes most evident, it is of considerable interest to learn if the foci have more, less, or the same level of enzymatic activity as the diffuse protein.

### YisK localization

The catalytic dead variant, YisK E148A E150A, retains punctate localization (Fig 7B) and still widens cells when overexpressed (Fig 7A). Conversely, cells are insensitive to overexpression of the YisK E30A variant which possesses wild-type enzymatic activity (Fig 6B) but localizes only diffusely (Fig 7C). Notably, the 11*mbl* 11*ponA* background which is also resistant to YisK overexpression (19) similarly supports only diffuse localization (Fig 3C and S2A). Although a direct interaction between YisK and Mbl would explain both YisK’s dependence on Mbl for localization and its overexpression phenotypes (19), we currently lack evidence supporting this model. The fact that YisK can interact directly with FtsE (Fig 2B) further links YisK to the elongasome; however, as FtsE is not required for YisK’s punctate localization (Fig S2F) or overexpression phenotypes (Fig 2C and 2D), we do not yet understand the functional significance of the interaction.

The small domain of YisK is structurally unique, as a VAST search identified only limited structural similarity for regions 1-29 and 74-82 that together constitute part of a domain of unknown function frequently fused to the N-terminus of FAH-like proteins; addressing the small domain’s role in YisK function could also inform our understanding of this uncharacterized protein fold. In contrast, region 30-73 had no structurally similar hits in the Protein Data Bank. Remarkably, a single substitution in the small domain (E30A) completely abolished formation of YisK’s Mbl-dependent foci, suggesting this region of YisK likely constitutes part of a localization domain. Localization of the catalytically inactive mutant (E148A, E150A) was indistinguishable from that of wild-type YisK (Fig 7B). Consistent with this observation, no significant conformational changes were detected in the small domain between the ligand-bound and apo states (Fig S3C). At the same time, we do not exclude the possibility that the ligand-dependent conformational changed in the lid region of YisK could be coupled to regulation of another partner and/or binding event.

We are left with the question as to why an enzyme implicated in central carbon metabolism would not only exhibit subcellular localization, but also evolve associations with components of the envelope synthesis machinery. There is evidence of higher order associations among the TCA cycle enzymes (54) and precedence for functional interactions between metabolic enzymes and the cell division protein FtsZ (55–58). To our knowledge YisK is the first example of an enzyme not obviously linked to envelope synthesis that depends on Mbl for localization. YisK’s interaction with the elongasome could be a means for the cell to couple central carbon metabolism with envelope synthesis and/or degradation. Alternatively, the direct interaction between YisK and the FtsEX complex required for the D,L-endopeptidase activity of CwlO may suggest YisK possesses an enzymatic activity or regulatory role more directly related to cell wall hydrolysis. Future experiments will aim to address the functional significance of YisK’s interaction with the elongasome.

## MATERIAL AND METHODS

### General methods

Details of strain construction can be found in the supplemental material. Cells were stored at -80°C in 15.0% glycerol (v/v). All LB is Lysogeny Broth-Lennox. Strains were streaked for isolation on LB plates containing 1.5% (w/v) bacto agar and incubated overnight (∼16 hr) at 37°C. Cultures for experiments were started from same-day plates by inoculating single colonies into a 20 mm glass tubes containing 5.0 mL of medium indicated in the associated figure. The tubes were incubated at 37°C in a roller drum until exponential stage. For experiments, the exponential pre-cultures were used to inoculated 25.0 mL of medium in a 250 mL baffled flask. Cells were incubated in a shaking water bath set to 280 rpm and 37°C.

Liquid LB was made by dissolving 20.0 g of LB-Lennox powder (Difco, product #240230) in 1.0 L ddH_2_O, followed by sterilization in an autoclave. LB-Mg indicates LB supplemented with 10.0 mM MgCl_2_. Each 1.0 L of CH (59) contained 10.0 g Casein acid hydrolysate (Acumedia, Lot No.104,442B), 3.7 g L-glutamic acid (25.0 mM), 1.6 g L-asparagine monohydrate (10.0 mM), 1.25 g L-alanine (14.0 mM), 1.36 g (10.0 mM) KH_2_PO_4_ anhydrous, 1.34 g (25.0 mM) NH_4_Cl, 0.11 g Na_2_SO_4_ (0.77 mM), 0.1 g NH_4_NO_3_(1.25 mM), and 0.001 g FeCl_3_•6H_2_O (3.7 mM) and ddH_2_O. The pH was adjusted to 7.0 with 10.0 N NaOH before the volume was brought up to 1.0 L with ddH_2_O, and the medium was sterilized in an autoclave. After autoclaving, sterilized solutions were added to give the following final concentrations: 0.18 mM CaCl_2_, 0.4 mM MgSO_4_, 0.1 mM MnSO_4_ and 0.1 mM L-Tryptophan. Sporulation by resuspension medium was made as described (59).

### Plate growth assays

*B. subtilis* strains were cultured in 5.0 mL LB tube on the roller drum at 37°C to OD_600_ ∼0.5. Then, cells are back diluted in LB to OD_600_ 0.1, 0.01, 0.001, 0.0001 and 0.00001. Five μL of each sample are spotted on LB-Lennox plates containing 100.0 μg/mL spectinomycin without and with 1.0 mM IPTG, respectively. Plates were incubated at 37°C overnight, and images were captured on a ScanJet G4050 flatbed scanner (Hewlett Packard).

### Bacterial two-hybrid analysis

Bacterial two-hybrid was performed as described (60) with the following modifications: cloning was carried out in the presence of 0.2% glucose. Cells harboring the relevant pairwise interactions were grown to early exponential phase in LB with 0.2% glucose, 50.0 μg/mL ampicillin and 25.0 μg/mL kanamycin. Samples were to an OD_600_ of 0.1 and 5.0 μL of each culture spotted on M9-glucose minimal media plates containing 40.0 μg/mL 5-bromo-4-chloro-3-indolyl-β-D-galactopyranoside (X-Gal), 250.0 μM isopropyl-β-D-thiogalactopyranoside (IPTG), 50.0 μg/mL ampicillin and 25.0 μg/mL kanamycin. Plates were incubated at 37°C in the dark for 20 hr for color development prior to image capture.

### Western blot analysis

Cultures were grown to exponential stage and one mL of culture was collected and spun at 21,130 X g for 1 min at room. OD_600_ value at time of sampling was recorded. The pellet was re-suspended in the lysis buffer (20.0 mM Tris pH [7.5], 10.0 mM EDTA, 1.0 mg/mL lysozyme, 10.0 µg/mL DNase I, 100.0 µg/mL RNase A, 1.0 mM PMSF, 1 µL protease inhibitor cocktail (Sigma P8465-5ML) to give a final OD_600_ equivalent of 15. The samples were incubated at 37°C for 10 min followed by addition of an equal volume of sodium dodecyl sulfate (SDS) sample buffer (0.25 M Tris pH 6.8, 4% (w/v) 20% (w/v) SDS, 20% (v/v) glycerol, 10.0 mM EDTA, and 10% (v/v) β-mercaptoethanol). Samples were boiled for 5 min prior to loading. Proteins were separated on 12% SDS-PAGE polyacrylamide gels and transferred to a nitrocellulose membrane at 100V for 60 min. membrane was blocked in 1X PBS containing 0.05% (v/v) Tween-20 and 5% (w/v) dry milk powder. The blocked membranes were then probed overnight at 4°C with anti-YisK antibody (1:10,000, rabbit serum) diluted in 1X PBS with 0.05% (v/v) Tween-20 and 5% (w/v) milk powder. The membranes were washed three times with 1X PBS containing 0.05% (v/v) Tween-20 before transferring to 1X PBS with 0.05% (v/v) Tween-20 and 5% (w/v) milk powder containing 1:5,000 horseradish peroxidase-conjugated goat anti rabbit IgG secondary antibody (Rockland 611-1302) and incubated on a shaking platform for 1 hr at room temperature. The membranes were washed 3X with PBS containing 0.05% (v/v) Tween-20 and signal was detected using SuperSignal West Femto maximum sensitivity substrate (Thermofisher) and a Biorad Gel Doc Imaging System.

### Fluorescence microscopy

For microscopy experiments, all strains were grown in the indicated medium in volumes of 25.0 mL in 250-mL baffled flasks and placed in a shaking water bath set at 37°C and 280 rpm. Unless stated otherwise, overexpression was performed in LB-Mg by inducing samples with 1.0 mM IPTG. Cells were always imaged below an OD_600_ of 0.7. Sporulation was performed by growing cells in CH medium (59) followed by resuspension in sporulation medium (59, 61). To capture images of membranes, one mL of cultured cells were harvested and concentrated by centrifugation at 6,010 X g for 1 min and re-suspended in 5.0 µL 1 X PBS with TMA-DPH (50.0 µM). Cells were mounted on glass sides with polylysine-treated coverslips prior to imaging. For the YisK-GFP experiments, cells were mounted on 1% (w/v) agarose pads made with ddH_2_O and overlaid with a coverslip. Cells were imaged on a Nikon Ti-E microscope with a CFI Plan Apo lambda DM 100X objective and Prior Scientific Lumen 200 illumination system. Filter cubes utilized were C-FL UV-2E/C (TMA), HC HISN Zero Shift (GFP) filter cubes. Micrographs were acquired with a CoolSNAP HQ2 monochrome camera using NIS Elements Advanced Research and processed in NIS Elements or ImageJ (62).

### Protein purification

Wild-type YisK, YisK E30A and YisK E148A E150A were expressed as C-terminal histidine tag fusions (strain construction in supplementary) by transforming competent BL21(λDE3)pLysS cells with the pET24b-based plasmids and selecting on LB agar plates supplemented with 25.0 μg/mL kanamycin. Following overnight growth at 37°C, colonies were aseptically scraped from the plate and resuspended in 5.0 mL of Cinnibar medium (Teknova). This suspension was used to inoculate a 250 mL baffled flask containing 25.0 mL Cinnibar supplemented with 0.1% (w/v) glucose and 25.0 μg/mL kanamycin to a starting OD_600_ of 0.1. Cultures were grown in shaking waterbath (250-280 rpm) at 37°C until culture reached an OD_600_ of approximately 5 at which point IPTG was added to 1.0 mM. The culture was then placed in a 16°C shaker overnight (∼16 hr). The cells were collected by centrifuging at 8,000 X g for 10 min at 4°C. The supernatant was aspirated, and the pellets were frozen at −80°C. To purify the protein, the pellet was resuspended in 25.0 mL lysis buffer (50.0 mM Tris-HCl [pH 8.0], 300.0 mM NaCl, 10.0 mM imidazole, 10% (v/v) glycerol, 200 μg/mL lysozyme, 10.0 μg/mL DNase, 1.0 μL per 35 OD*mL protease inhibitor cocktail (Sigma P8465), 0.5 mL 100.0 mM phenylmethylsulphonyl fluoride and lysed using a LM20 Microfluidizer (Microfluidics) at 20,000 psi. Cell debris was removed by centrifugation at 41,656 X g for 30 min at 4°C.

The supernatant was collected and at room temperature was passed over a 2.0 mL bed volume of Ni-NTA Sepharose resin (Qiagen). The column was then washed three times with 10.0 mL of wash buffer (50.0 mM Tris-HCl [pH 8.0], 300.0 mM NaCl, 25.0 mM imidazole, and 10% (v/v) Glycerol). Elutions were carried out with 12 x 0.5 mL of elution buffer (50.0 mM Tris-HCl [pH 8.0], 300.0 mM NaCl, 10% (v/v) glycerol), and 300.0 mM imidazole. Fractions containing the protein were pooled and dialyzed against 20.0 mM Hepes [pH 7.5], 150.0 mM NaCl, 15% (v/v) glycerol, and 2.0 mM DTT and frozen as aliquots at -80°C. These protein stocks were used in the differential scanning fluorimetry and enzyme activity assays. Protein purified for crystallization was dialyzed against 50.0 mM Tris-HCl [pH 7.5] and 1.0 mM DTT and used immediately.

### YisK crystallization

586 separate crystallization conditions were screened in a 96 well plate using a TPP LabTech Mosquito LCP machine. Initial crystals were obtained in 60% Tascimate (Hampton Research) [pH 7.5], and 10% Tascimate (Hampton Research) [pH 5.0], 20% (w/v) polyethylene glycol (PEG) 2000 with various divalent cations. Optimized conditions used YisK-6His at 12.0 mg/mL (by Bradford Assay) in buffer containing 50.0 mM Tris [pH 7.5], and1.0 mM DTT set in a 1:1 volume ratio and 2:1 ratio with mother liquor (60% Tascimate [pH 7.0], 100.0 mM MnCl_2_). Initial solution based on these crystals indicated dicarboxylic components from tacsimate binding in the active site. To find tacsimate-free crystallization condition, we pre-made YisK as a complex with Mn^2+^ and oxalate (at 3.0 mM final concentration), and repeated crystal screening. PEG-based crystal hit conditions were chosen for subsequent optimization. Oxaloacetate complexed YisK, which resulted in an apparent apo structure, and oxalomalate complexed YisK, resulted in a HOGA-bound structure (Fig, were crystallized with mother liquor of 0.2 M Mg-acetate and 20% (w/v) PEG 3350. Oxalate-bound YisK was crystallized with 0.1 M AmSO4, 25% (w/v) PEG-4000 and 15% (v/v) glycerol. All crystals were cryoprotected by adding glycerol to the corresponding mother liquor to the final concentration of 30% (v/v) and flash frozen in a liquid nitrogen for the data collection.

### Data collection and analysis

Oxalate-bound structure was solved from the crystal collected at BL502 beamline of Lawrence Berkley National Lab synchrotron; two other structures resulted from the data collected at 23ID beamline of Argonne National Lab synchrotron. Data were indexed, integrated and scaled by the beamline auto-processing pipeline: XDS (63), POINTLESS (64) and AIMLESS (65) software packages. The structure was solved by molecular replacement with Molrep software (66) using manually edited Phyre2 generated homology model as a search model for the first solution. This was followed by iterative cycles of refinement with PHENIX.REFINE and manual building in COOT (67, 68). A ligand model and restraints were created using the eLBOW tool (69). Initial solution showed poor refinement R factors, and upon examination of the electron density maps it became evident that while the larger catalytic domain refined well to the homology model, the smaller domain was substantially different both in fold and position relative to the main domain. The smaller domain was fully rebuilt through stepwise iterative building and refinement cycles. The data collection and refinement statistics are shown in Table 1.

**Table 1.**

|  | APO FORM (8SUU) | OXALATE-BOUND<br>(8SKY) | 4-HYDROXY-2-OXOGLUTARATE-BOUND (8SUT) |
| --- | --- | --- | --- |
| **Data collection** |  |  |  |
| Space group | *P2<sub>1</sub>2<sub>1</sub>2<sub>1</sub>* | *P2<sub>1</sub>2<sub>1</sub>2<sub>1</sub>* | *P2<sub>1</sub>2<sub>1</sub>2<sub>1</sub>* |
| Cell dimensions |  |  |  |
| *a*, *b*, *c* (Å) | 53.94, 93.77, 125.48 | 52.34, 94.84, 123.94 | 53.69, 93.98, 124.98 |
| $\alpha$, $\beta$, $\gamma$ (°) | 90.00, 90.00, 90.00 | 90.00, 90.00, 90.00 | 90.00, 90.00, 90.00 |
| Resolution (Å) | 46.93 – 2.26 (2.32-2.26) | 48.21-2.4 (2.49-2.4) | 46.99-1.93 (1.98-1.93) |
| *R*<sub>sym</sub> or *R*<sub>merge</sub> | 0.117 (2.209) | 0.135 (1.46) | 0.068 (1.421) |
| *I*/σ*I* | 11.9 (0.7) | 14 (1.2) | 12.4 (0.7) |
| Completeness (%) | 100 (100) | 99 (99.7) | 98.2 (83.2) |
| Redundancy | 6.3 (5.5) | 11.5 (6.2) | 4.6 (3.1) |
| CC1/2 | 0.998 (0.298) | 0.998 (0.386) | 0.998 (0.259) |
| **Refinement** |  |  |  |
| Resolution (Å) | 46.93 – 2.26 | 48.21-2.4 | 46.99-1.93 |
| No. reflections | 28549 | 24570 | 45270 |
| *R*<sub>work</sub> / *R*<sub>free</sub> | 0.211 / 0.255 | 0.189 / 0.236 | 0.183 / 0.23 |
| No. atoms |  |  |  |
| Protein | 4500 | 4674 | 4668 |
| Ligand(s) | 2 | 14 | 24 |
| Water | 50 | 117 | 148 |
| B-factors (median) |  |  |  |
| Protein | 61 | 45 | 43 |
| Ligand(s) | 101 | 60.5 | 64 |
| R.m.s deviations |  |  |  |
| Bond lengths (Å) | 0.009 | 0.008 | 0.014 |
| Bond angles (°) | 1.141 | 1.171 | 1.36 |

\*Highest resolution shell is shown in parenthesis.

### Differential scanning fluorimetry (thermal shift assays)

Differential Scanning Fluorimetry was performed as previously (70), with minor modifications. In the screen for stabilizing molecules, a master mix of either 1.0 or 10.0 μM YisK-His (concentration indicated in table legend) in 20.0 mM Hepes [pH 7.5], 150.0 mM NaCl, and 5.0 mM MnCl_2_ or when indicated in the table legend, 1.0 mM MnCl_2_, 1.0 mM MgCl_2_, 1.0 mM CaCl_2_ was made, including 5X Sypro Orange. The 5000X Sypro Orange was added to master mix before YisK-His to avoid possible effects of the concentrated Sypro Orange solvent. Thirty-nine μL of the master mix was aliquoted into each well on a BioRad Hard-Shell PCR Plates, 96-well format, thin wall (HSP9601). To each well, 1.0 μL of compound was added to give a final concentration of 1.0 mM. Reactions were mixed by repeated pipetting after adding compound. The PCR plate was then sealed with BioRad Microseal ‘B’ seal (MSB1001) and centrifuged at 1000 x g for 3 min in a centrifuge equipped with a plate adaptor at RT to remove bubbles. The plate was placed in a CFX96 Touch Real Time PCR machine running a custom program as follows: initial temperature, 25°C ramped to 95°C at a rate of 0.5 °C/min, acquiring the signal from the FRET channel. The final data was analyzed using BioRad CFX Manager, with baseline subtracted curve fit analysis mode. The derivative of the melting curve was used to determine the melting temperature of YisK in all assayed conditions.

To assay for apparent metal binding affinity, a master mix of 10.0 μM YisK-His in 20.0 mM Hepes [pH 7.5], 150 mM NaCl, 5X Sypro Orange was made with gradient MnCl_2_ or MgCl_2_. Thirty-nine μL of this master mix was aliquoted into each well on a BioRad Hard-Shell PCR Plates, 96-well format, thin wall (HSP9601). To each well either 1.0 μL of reaction buffer alone (20.0 mM Hepes pH 7.5, 150 mM NaCl) or 1.0 μL of a MnCl_2_ or MgCl_2_ were added to give the final gradient concentration indicated in the figures. The samples were then processed as described for the compound screen.

### Determination of Kinetic Constants

The kinetic constants for YisK were determined by following the conversion of NADH to NAD^+^ at 340 nm at 25°C with a Spectramax340 UV-visible spectrophotometer. Assays were performed in 50.0 mM Hepes [pH 7.5], 150.0 mM NaCl, 1.0 mM MnCl_2_, 0.3 mM NADH, and 1 unit lactate dehydrogenase in 96-well NucC plates. The spontaneous rate of decarboxylation was monitored in buffer containing all but the addition of YisK. The reactions with YisK were initiated with the addition of 0.5 µM protein to the reaction well. Kinetic constants were determined with variable levels of OAA (0.031 mM to 4.0 mM). The kinetic parameters were determined by fitting the initial rates to the equation *v*/E_t_=*k*_cat_(S)/(*K*_m_+S) using Prism 9 graph, where *v* is the initial velocity of the reaction, E_t_ is the enzyme concentration, *k*_cat_ is the turnover number, [S] is the substrate concentration, and *K*_m_ is the Michaelis constant.

### Statistical analysis and data plotting

Graphs were generated and statistical analysis was performed using GraphPad Prism version 9.4.0 for Mac (GraphPad Software, San Diego, California USA, www.graphpad.com).

### Whole-genome sequencing and analysis

Single colonies of Δ*ponA\** (BYD172) and Δ*ponA* (BYD048) were inoculated into 5.0 mL LB-Mg and grown at 37 °C for 4 hr in a roller drum. One mL of cells were collected by centrifuging at 21,130 x g (room temperature) for 2 min, then aspirating the supernatant. Pellets were resuspended in lysis buffer (20.0 mM Tris-HCl pH 7.5, 50.0 mM EDTA [pH 8.0], 100.0 mM NaCl, and 2.0 mg/mL lysozyme) and incubated at 37 °C for 30 min before sarkosyl was added to a final concentration of 1.0% (w/v). Protein was extracted with 600 μL phenol by vortexing the lysate and phenol before centrifuging at 21,130 x g for 5 min at room temperature. The aqueous layer (top) was transferred to a new microcentrifuge tube. This was followed by an extraction with 600 μL phenol-saturated chloroform as described above. After transferring the aqueous layer to a new microcentrifuge tube, a final extraction was performed with 100% chloroform, being careful to avoid the interphase material. To precipitate the genomic DNA, a 1/10th volume of 3.0 M Na-acetate and 1.0 mL of 100% ethanol was added, and the tube was inverted multiple times. The sample was centrifuged at 21,130 x g for 1 min at room temperature and the pellet was washed with 150 μL 70% (v/v) ethanol before being resuspended in 500 μL TE (10.0 mM Tris [pH 7.5], 1.0 mM EDTA [pH 8.0]). To eliminate potential RNA contamination, RNase was added to a final concentration of 200 μg/mL and the sample was incubated at 55 °C for 1 hr. To remove the RNase, the genomic DNA was re-purified by phenol-chloroform and chloroform extraction with ethanol precipitation as described above. The final pellet was resuspended in 100 μL TE buffer. Bar-coded libraries were prepared from each genomic DNA sample using a TruSeq DNA kit according to manufacture specifications (Illumina), and the samples were subjected to Illumina-based whole-genome sequencing using a MiSeq 250 paired-end run (Illumina). CLC Genomics Workbench (Qiagen) was used to map the sequence reads against the *Bacillus subtilis* 168 reference genome, and single nucleotide polymorphisms, insertions, and deletions were identified.

## ACKNOWLEDGEMENTS

We thank Craig D. Kaplan for feedback on the manuscript and Frank M. Raushel for helpful discussions and use of his spectrophotometer. This work was supported by funds from The College of AgriLife and Department of Biochemistry and Biophysics at Texas A&M University and a grant from the National Science Foundation (MCB-1514629) to J.K.H.

## Supplementary materials

**Figure S1.**
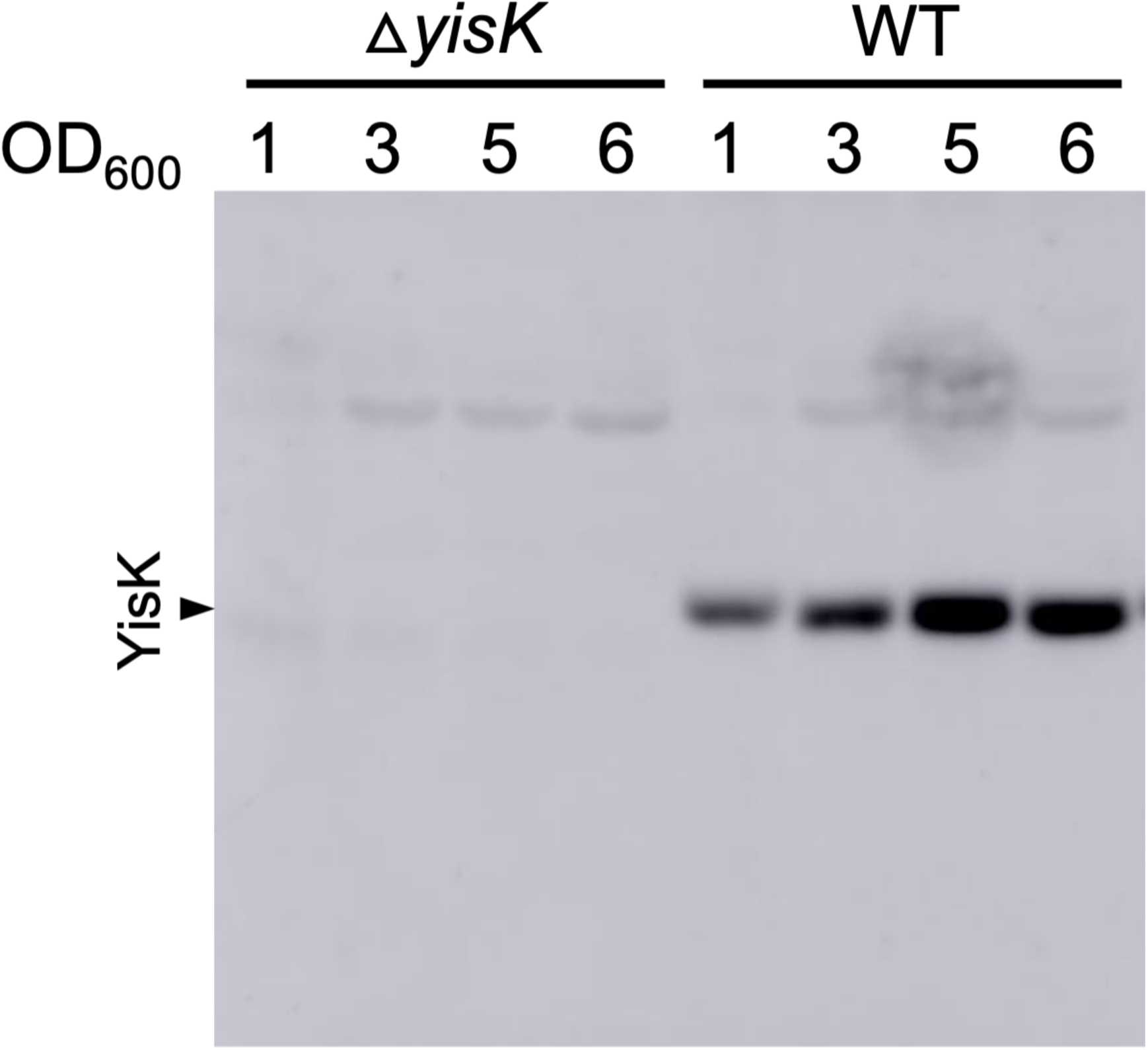
Specificity of α-YisK rabbit polyclonal antibody (serum). Wild-type (BJH004) and Δ*yisK* (BYD278) were grown at 37°C in LB-Mg and collected at the indicated OD_600_.

**Figure S2.**
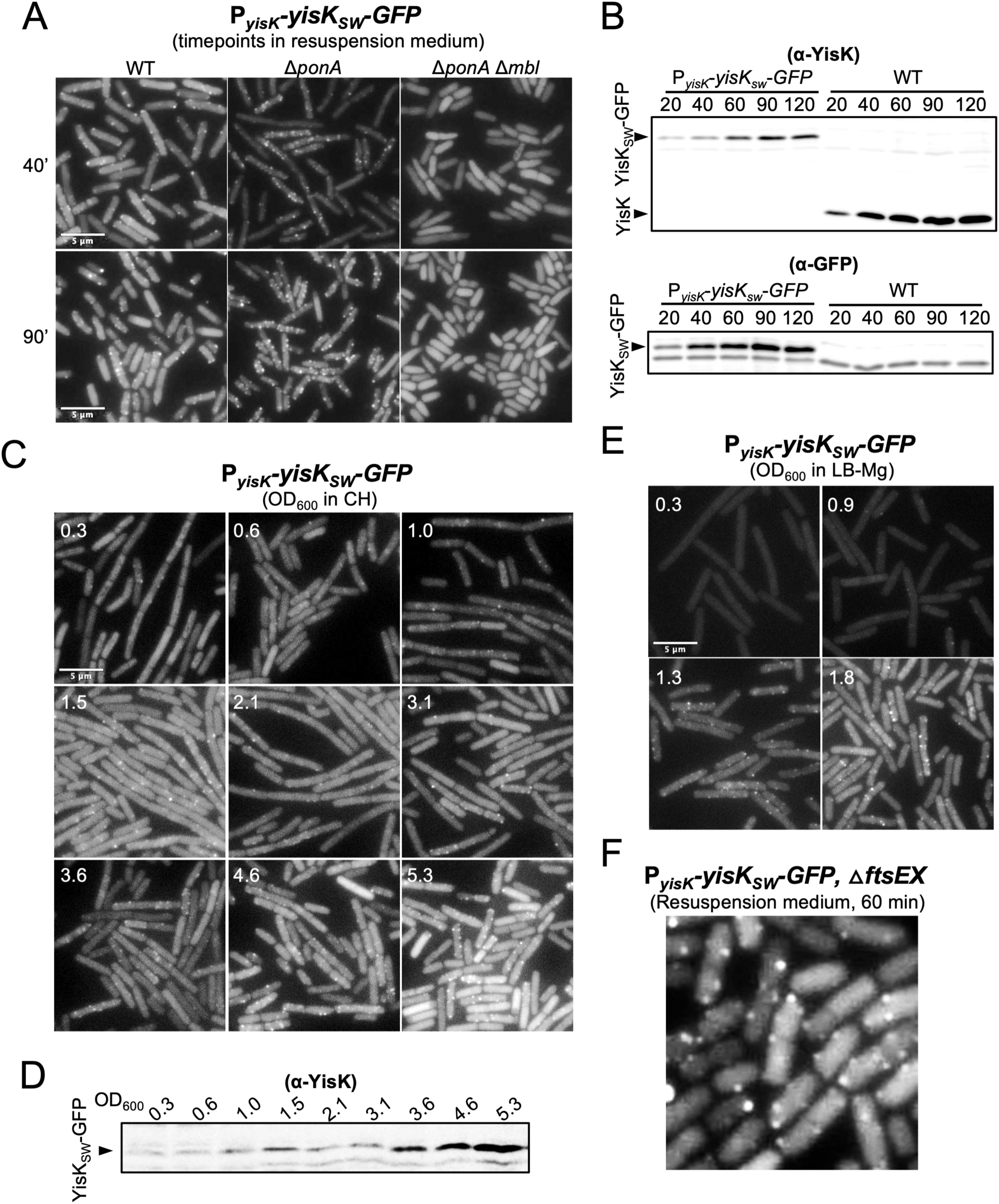
(A, C, E, and F) Representative micrographs of cells harboring P_yisK_*-yisK*_SW_*-GFP* during growth in the indicated media. Images within the same medium are shown at the same magnification and scaled identically in the GFP channel. Western blot analysis in (B) sporulation by resuspension medium and (D) CH.

**Figure S3.**
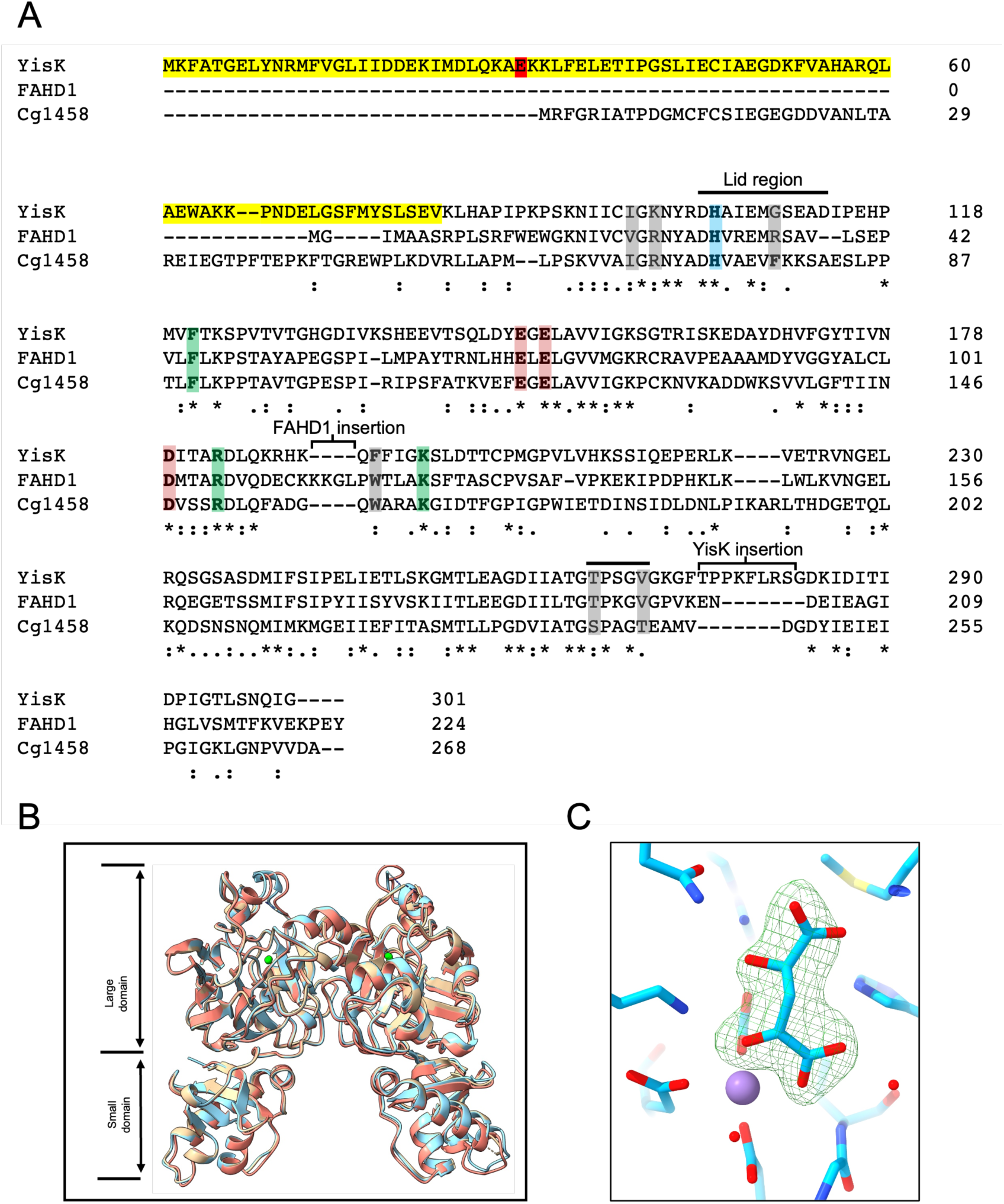
(A) Clustal O sequence alignment of YisK, FAHD1, and Cg1458. Residues corresponding to the small domain of YisK are highlighted in yellow with the E30 residue in red. Catalytic histidine (blue). Residues coordinating Mn^2+^ (pink). Invariant residues of active site (green). Residues that vary in/around the active site between YisK and FAHD1 and/or Cg1458 (grey). (B) Product of oxalomalate decarboxylation, HOGA, bound in the active site of YisK. Polder omit map contoured at 5 sigma level displayed as green mesh. Model is shown as sticks with protein and ligand carbons (light blue), oxygens (red), nitrogens (blue) and Mn2+ (purple). (C) Overlay of YisK structures in unliganded (8SUU, beige), oxalate-bound (8SKY, salmon), and HOGA-bound (8SUT, light blue). The Mn^2+^ ion is shown as a green sphere.

**Figure S4.**
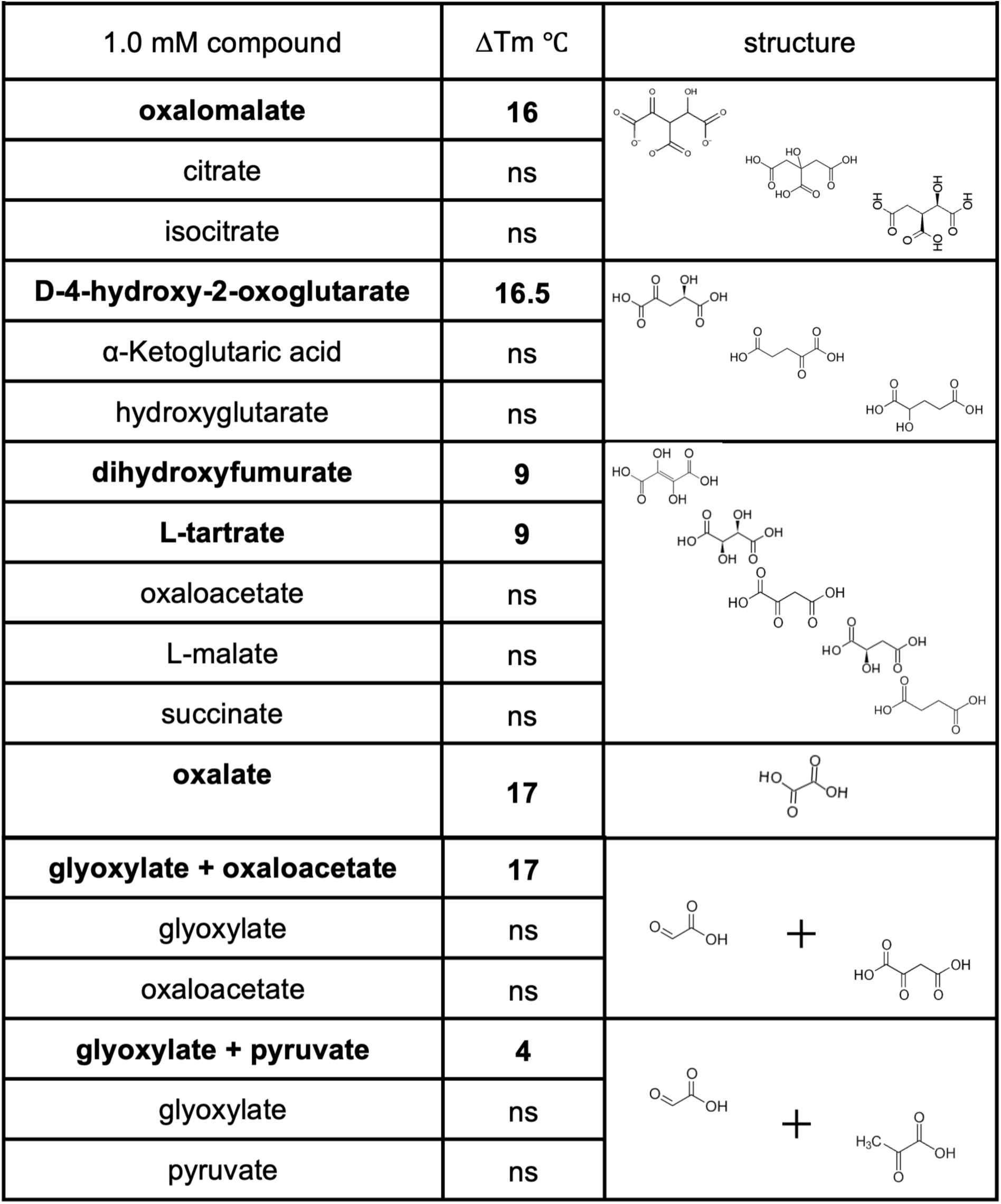
YisK differential scanning fluorimetry screen. Assay contained 1.0 µM YisK, 1.0 mM MnCl_2_, 1.0 mM MgCl_2,_ 1.0 mM CaCl_2_ and 1.0 mM of each the indicated compounds. Compounds in bold stabilized YisK significantly (>2.5°C). Compounds that stabilized YisK ≤ 2.5°C were not considered significant (ns). Oxaloacetate spontaneously decarboxylates to pyruvate and CO_2_ and may be unstable in the assay.

**Figure S5.**
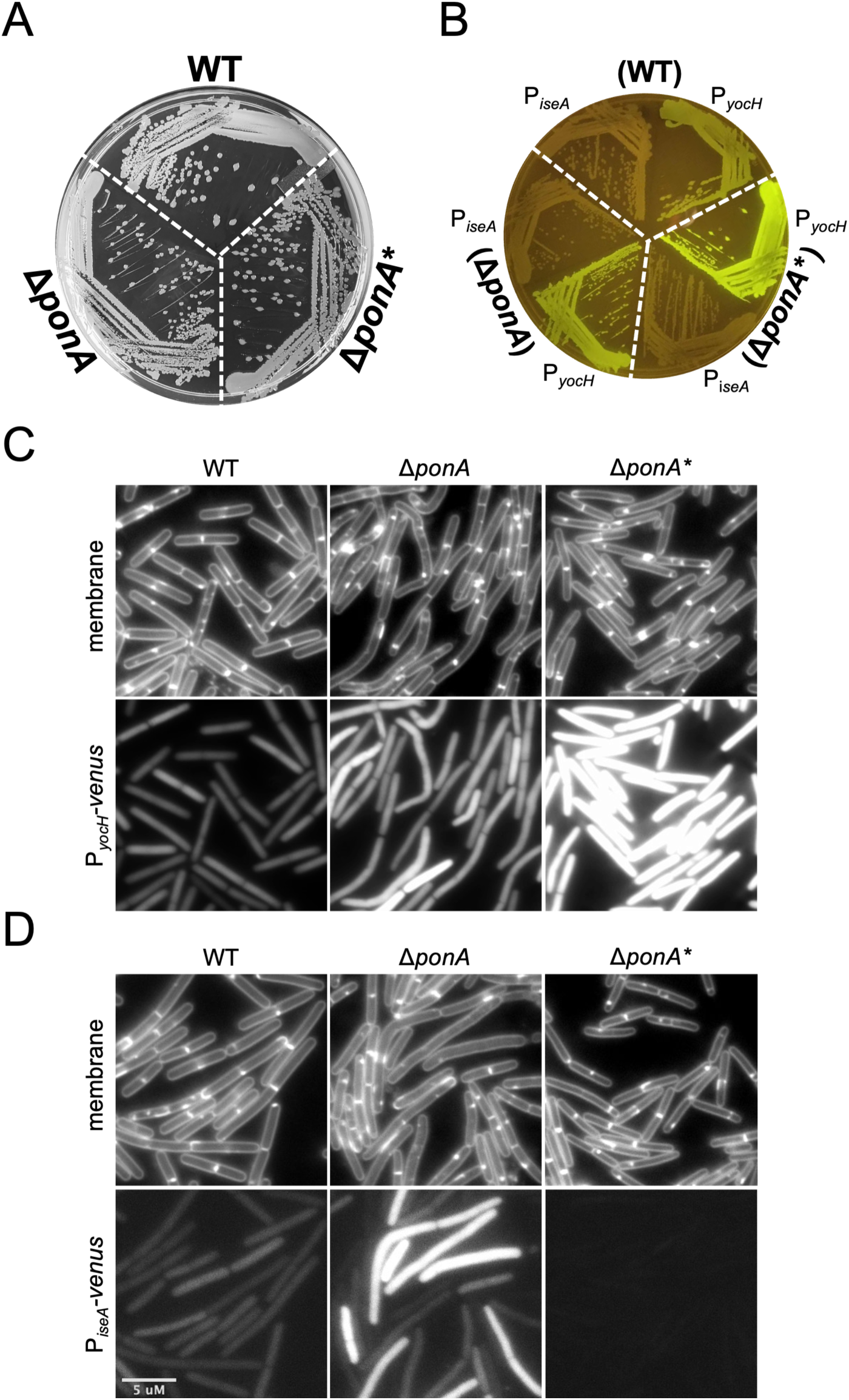
(A) Growth of WT 168 (BJH004), Δ*ponA* (BTG042), and Δ*ponA\** (*walH* Q242stop) on an LB-Mg plate. Strains harboring P_yocH_-*venus* or P_iseA_*-venus* were grown at at 37°C (B) on an LB-Mg plate and imaged under blue light or (C and D) imaged during exponential stage growth in liquid LB-Mg. Membranes were stained with TMA. All images are shown at the same magnification. The images for each reporter set are scaled identically (1000-16,500 for P_yocH_-*venus* and 200-1000 for P_iseA_-*venus*).

**Table S1.** Compounds tested in thermal assay. Assays contained either 10.0 µM YisK, 5.0 mM MnCl_2_, and 1.0 mM of each the indicated compounds or, when indicated by (*), 1.0 µM YisK, 1.0 mM MnCl_2_, 1.0 mM MgCl_2,_ 1.0 mM CaCl_2_ and 1.0 mM of each the indicated compounds. Compounds in bold stabilized YisK ≥ 10℃. Compounds that stabilized YisK ≤ 2.5°C were not considered significant (ns). (**) compounds may be unstable in assay.

| 1.0 mM each compound | $\Delta T_m$ °C | 1.0 mM each compound | $\Delta T_m$ °C |
| --- | --- | --- | --- |
| acetate* | ns | NAD+ | ns |
| ammonium chloride | ns | NADH | ns |
| formate | ns | NADP+ | ns |
| urea* | ns | ATP | ns |
| D-glyceric acid* | ns | ADP | ns |
| D-3-phosphoglycerate | ns | AMPPNP | ns |
| phosphoenolpyruvate | ns | D-malate | ns |
| pyruvate* | ns | D-malate + fumarate | ns |
| pyruvate + urea* | ns | fumarate + pyruvate | ns |
| citrate* | ns | L-malate* | ns |
| isocitrate* | ns | L-lactate* | ns |
| 2-oxoglutarate* | ns | glycolic acid | ns |
| succinic semialdehyde* | ns | dihydroxyfumarate | 9.0 |
| succinate* | ns | **oxalomalate\*\*** | **16.0** |
| fumarate* | ns | **D-4-hydroxy-2-oxoglutarate\*** | **16.5** |
| oxaloacetate** | ns | glyoxylate* | ns |
| oxaloacetate** + acetate | ns | glyoxylate + fumarate * | 3.5 |
| oxaloacetate** + pyruvate | ns | glyoxylate + malate* | 3.5 |
| oxaloacetate** + urea | ns | glyoxylate + pyruvate* | 4.0 |
| glyoxylate + succinate* | ns | **glyoxylate + oxaloacetate\*\*** | **17.0** |
| glyoxylate + urea* | ns | L-tartrate* | 9.0 |
| glyoxylate + 2-oxoglutarate* | ns | L-tartrate + acetate* | 7.5 |
| D-alanine | ns | L-tartrate + urea* | 7.5 |
| L-asparagine | ns | **L-tartrate + glyoxylate\*** | **10.0** |
| L-aspartate | ns | **oxalate\*** | **17.0** |
| L-glutamate | ns | **oxalate + fumarate\*** | **16.5** |
| D-aspartate | ns | **oxalate + succinate\*** | **17.0** |
| D-glutamate | ns | **oxalate + pyruvate\*** | **17.5** |
| D-alanine-D-alanine | ns | **oxalate + malate\*** | **17.5** |
| cyclic D-alanine L-alanine | ns | **oxalate + L-tartrate\*** | **18.0** |
| meso-2,6-diaminopimelic acid (mDAP) | ns | **oxalate + oxaloacetate\*\*** | **18.5** |
| ampicillin | ns | **oxalate + 2-oxoglutarate\*** | **19.0** |
| carbenicillin | ns |  |  |
| 2-oxoadipic acid | ns |  |  |

**Table S2.** Strains.

| Strain | Wildtype (WT) Strains | Reference |
| --- | --- | --- |
| BJH004 | *Bacillus subtilis* 168 | Bacillus Genetic Stock Center 1A1 |
|  | ***Bacillus subtilis* 168 derivatives** |  |
| BKE22320 | ΔponA*::erm | Bacillus Genetic Stock Center |
| BAS041 | amyE::P<sub>hy</sub>-yisK (spec) | (1) |
| BAS319 | ΔftsEX::tet | This study |
| BGD110 | amyE::P<sub>iseA</sub>-optRBS-venus (cat) | (2) |
| BGD300 | amyE::P<sub>yocH</sub>-optRBS-venus (cat) | (2) |
| BTG010 | amyE::P<sub>hy</sub>-yisK E148A E150A (spec)<br>yhdG::P<sub>hy</sub>-yisK E148A E150A (phleo) | Fig 7 |
| BTG011 | amyE::P<sub>hy</sub>-yisK E30A (spec)<br>yhdG::P<sub>hy</sub>-yisK E30A (phleo) | Fig 7 |
| BTG042 | ΔponA::erm (clean background) | Fig S5 |
| BTG068 | amyE::P<sub>hy</sub>-yisK<sub>SW</sub>-GFP (spec) | Fig 3 |
| BTG078 | yisK::yisK<sub>SW</sub>-GFP | Fig 2, Fig S2 |
| BTG079 | yisK::yisK<sub>SW</sub>-GFP<br>ΔftsEX::tet | Fig 2 |
| BTG081 | yisK::yisK<sub>SW</sub>-GFP, ponA::erm | Fig 3 |
| BTG082 | yisK::yisK<sub>SW</sub>-GFP, ponA::erm, Δmbl (kan-linked) | Fig 3 |
| BTG088 | yisK::yisK<sub>SW</sub>-GFP E30A | Fig 7F |
| BTG160 | amyE::P<sub>hy</sub>-yisK<sub>SW</sub>-GFP E148A E150A (spec) | Fig 7 |
| BTG169 | trpC+ | (3) |
| BTG278 | amyE ::P<sub>yocH</sub>-optRBS-venus (cat) | Fig S5 |
| BTG279 | amyE::P<sub>iseA</sub>-optRBS-venus (cat) | Fig S5 |
| BTG468 | ΔponA*::erm (walH Q242stop)<br>amyE ::P<sub>yocH</sub>-optRBS-venus (cat) | Fig S5 |
| BTG469 | ΔponA*::erm (walH Q242stop)<br>amyE::P<sub>iseA</sub>-optRBS-venus (cat) | Fig S5 |
| BTG470 | ΔponA::erm (clean background)<br>amyE ::P<sub>yocH</sub>-optRBS-venus (cat) | Fig S5 |
| BTG471 | ΔponA::erm (clean background)<br>amyE::P<sub>iseA</sub>-optRBS-venus (cat) | Fig S5 |
| BTG607 | amyE::P<sub>hy</sub>-yisK E30A (spec), trpC+ | Fig 7D |
| BTG608 | amyE::P<sub>hy</sub>-yisK (spec), trpC+ | Fig 7D |
| BTG895 | amyE::P<sub>hy</sub>-yisK (spec)<br>ΔftsEX::tet | Fig 2 |
| BYD074 | amyE::P<sub>hy</sub>-yisK (spec)<br>yhdG::P<sub>hy</sub>-yisK (phleo) | (1) |
| BYD172 | ponA*::erm (walH Q242stop) | BKE22320 (4) |
| BYD278 | ΔyisK | (1) |
| BYD587 | amyE::P<sub>hy</sub>-yisK E148A E150A (spec) | Fig 7 |
| BYD588 | amyE::P<sub>hy</sub>-yisK E30A (spec) | Fig 7 |
| BTH101 | F-, *cya-99*, *araD139*, *galE15*, *galK16*, *rpsL1* (*Str r*), *hsdR2*, *mcrA1*, *mcrB1* | (5) |
| cYD372 | *empty-T18 (amp)*, *T25-Mbl (kan)* | Fig 2A |
| cYD420 | *Mbl-T25 (kan)*, *empty-T18 (amp)* | Fig 2A |
| cYD565 | *YisK-T18 (amp)*, *Mbl-T25 (kan)* | Fig 2A |
| cYD566 | *YisK-T18 (amp)*, *T25-Mbl (kan)* | Fig 2A |
| cYD784 | *yisK-pET24b (kan)* | This study |
| cYD962 | *YisK-T18 (amp)*, *FtsE-T25 (kan)* | Fig 2B |
| cYD970 | *YisK-T18 (amp)*, *FtsX-T25 (kan)* | Fig 2B |
| cYD978 | *YisK-T18 (amp)*, *FtsEX-T25 (kan)* | Fig 2B |
| cYD982 | *YisK-T25 (kan)*, *empty-T18(amp)* | Fig 2B |
| cYD984 | *YisK-T18 (amp)*, *empty-T25 (kan)* | Fig 2B |
| cYD985 | *empty-T18 (amp)*, *FtsE-T25 (kan)* | Fig 2B |
| cYD987 | *empty-T18 (amp)*, *FtsEX-T25 (kan)* | Fig 2B |
| cYD988 | *YisK-T18 (amp)*, *T25-empty (kan)* | Fig 2B |
| cYD1017 | *empty-T18 (amp)*, *FtsX-T25 (kan)* | Fig 2B |
| cYD1077 | *yisK-pET24b E148A E150A (kan)* | This study |
| cYD1080 | *YisK-T18 (amp)*, *YisK-T25(kan)* | Fig 7 |
| cYD1154 | *yisK-pET24b E30A (kan)* | This study |
| cYD1157 | *YisK-T18 E30A (amp)*, *YisK-T25 E30A (kan)* | Fig 7 |
| cYD1158 | *empty-T18 (amp)*, *YisK-T25 E30A (kan)* | Fig 7 |
| cYD1159 | *empty-T25 (kan)*, *YisK-T18 E30A (amp)* | Fig 7 |

**Table S3.** Plasmids.

| Plasmid | Description | Reference |
| --- | --- | --- |
| pDR111 | *amyE::P<sub>hy</sub> (spec)(amp)* | David Rudner |
| pMiniMAD2 | *ori<sup>BsTs</sup> (amp)(erm)* | (6) |
| pTG004 | *pMiniMAD2 yisK<sub>SW</sub>-GFP (amp)(erm)* | This study |
| pTG006 | *pMiniMAD2 yisK<sub>SW</sub>-GFP E30A (amp)(erm)* | This study |
| pKT25 | *T25-empty (kan)* | Tom Bernhardt |
| pKNT25 | *empty-T25 (kan)* | Tom Bernhardt |
| pCH364 | *empty-T18 (amp)* | Tom Bernhardt |
| pYD206b | *Mbl-T25 (kan)* | This study |
| pYD207b | *T25-Mbl (kan)* | This study |
| pYD216 | *YisK-T18 (amp)* | This study |
| pYD218 | *YisK-T25 (kan)* | This study |
| pET24b | *C-terminal 6X-His tag vector (kan)* | Novagen |
| pYD104 | *yisK-pET24b (kan)* | This study |
| pYD220 | *FtsE-T25 (kan)* | This study |
| pYD221 | *FtsX-T25 (kan)* | This study |
| pYD227 | *FtsEX-T25 (kan)* | This study |
| pYD228 | *yisK-pET24b E148A E150A (kan)* | This study |
| pYD229 | *YisK-T25 E30A (kan)* | This study |
| pYD230 | *YisK-T18 E30A (amp)* | This study |
| pYD231 | *yisK-pET24b E30A (kan)* | This study |

Plasmid inserts and fusions were confirmed by sequencing.

**Table S4.**
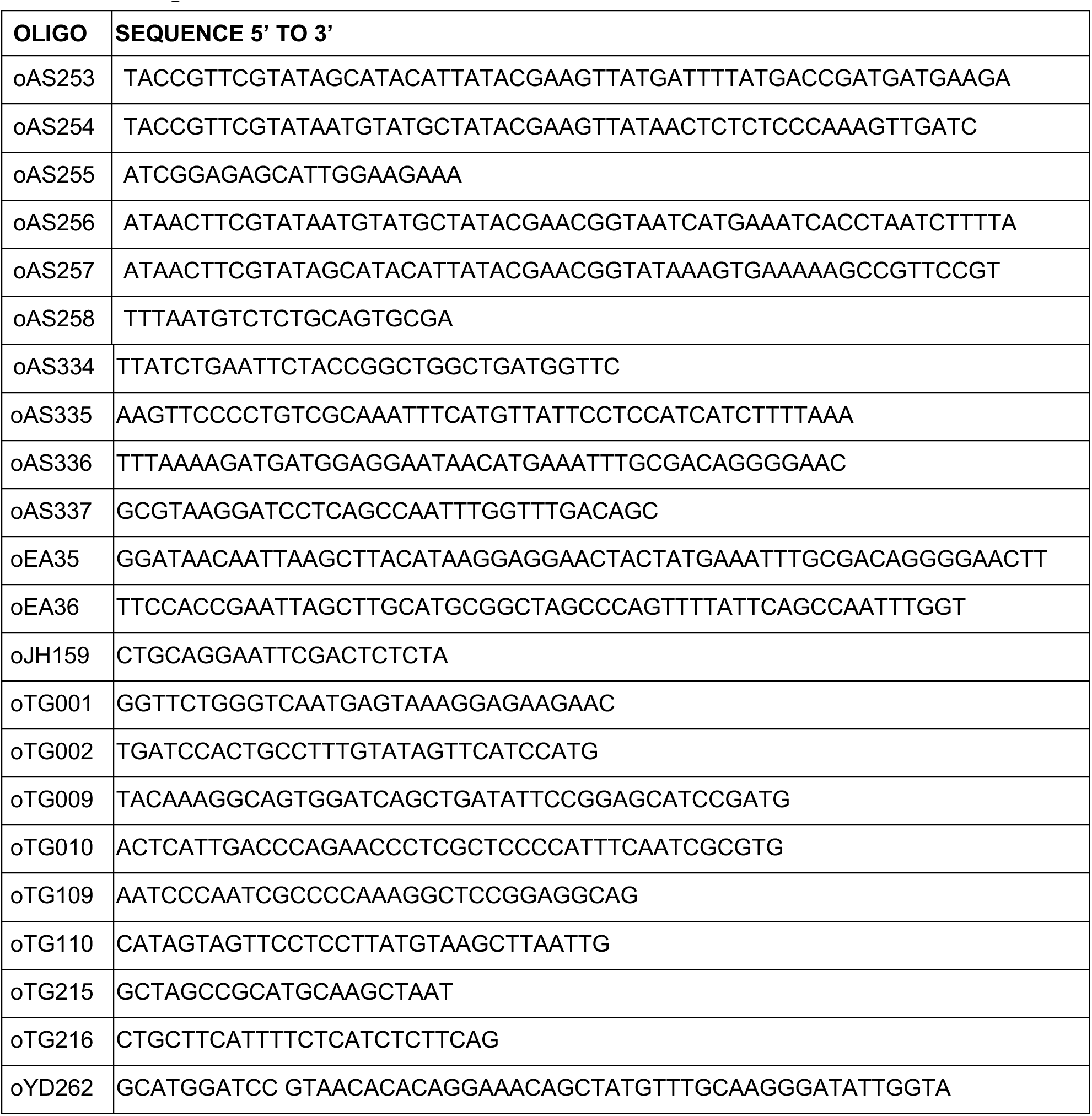
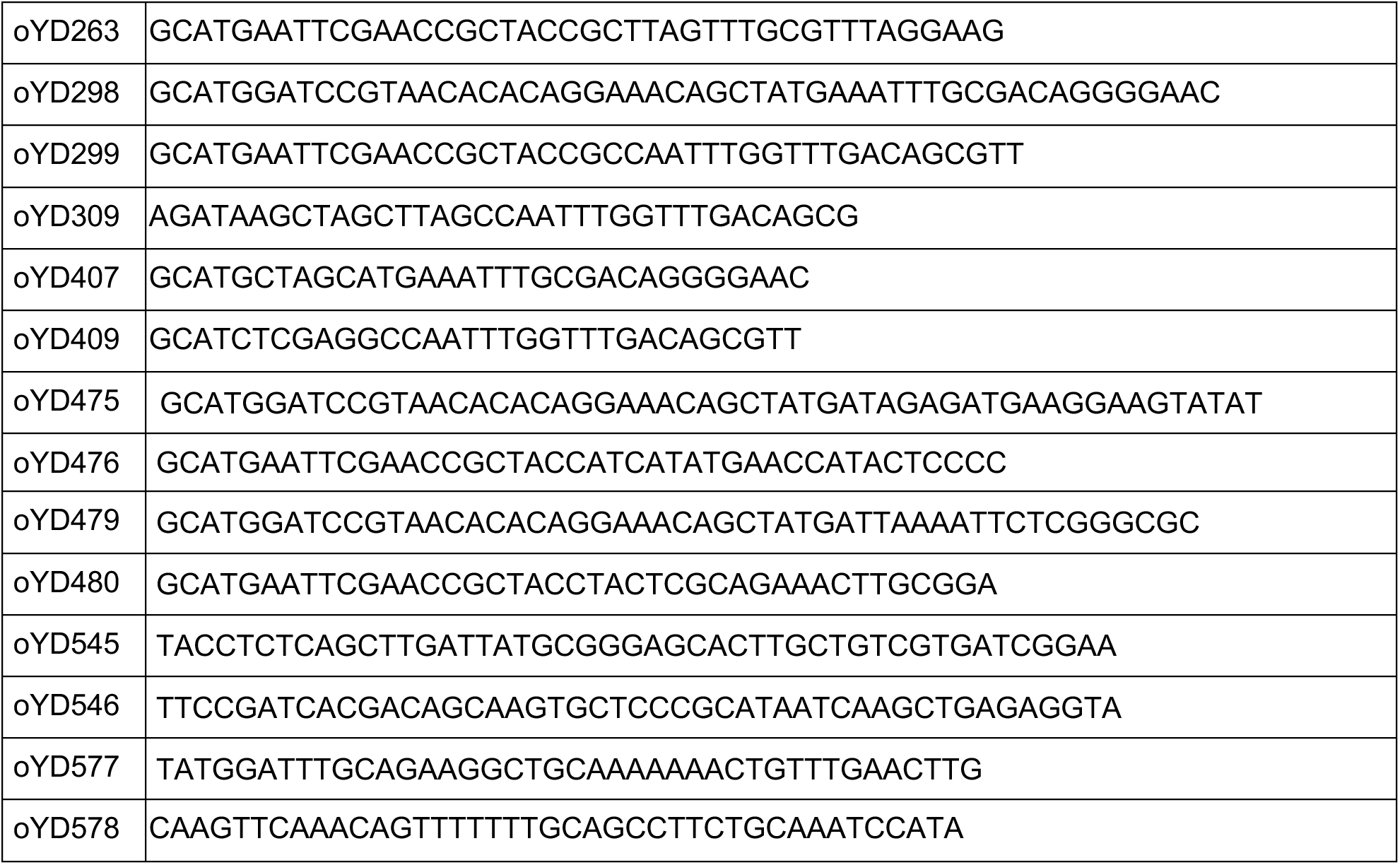
Oligonucleotides.

## Text S1. Strain construction

Deletion/knockout strains were confirmed by PCR. Point mutations were confirmed by PCR and sequencing. **BAS319** [△*ftsEX::tet*] was created by by 3-part Gibson assembly with a region upstream of *ftsEX* (amplified with primer pair oAS255 and oAS256), a tetracyline cassette flanked by Lox recombination sites (amplified with primer pair oAS255 and oAS256) and a region downstream of *ftsEX* (amplified with primer pair oAS257 and oAS258). The assembly reaction was transformed into *B. subtilis* 168, selecting for tetracycline resistance.

**BTG010** [*amyE::P_hy_ -yisK E148A E150A (spec), yhdG::P_hy_-yisK E148A E150A (phleo)*] was created by transforming *B. subtilis 168* with a linear Gibson assembly product encoding three fragments including a region upstream of *amyE* with a spectinomycin cassette using oTG109 and oTG110, the *P_hy_-yisK E148A E150A* using oEA35 and oEA36 amplified from pYD228, a region downstream of *amyE* into *B. subtilis* 168 using oTG215 and oTG216, and selecting for growth on LB plates containing 100 µg ml-1 spectinomycin. The confirmed clone was then transformed ScaI-linearized plasmid *yhdG::P_hy_-yisK E148A E150A (phleo)*, selecting for for growth on LB plates containing 400 µg ml-1 phleomycin.

**BTG011** [*amyE::P_hy_ -yisK E30A (spec), yhdG::P_hy_-yisK E30A (phleo)*] *amyE::P_hy_ -yisK E30A (spec)* was created by transforming *B. subtilis* 168 with a linear Gibson assembly product encoding three fragments including a region upstream of *amyE* with a spectinomycin cassette using oTG109 and oTG110, the *P_hy_-yisK E30A* using oEA35 and oEA36 amplified from pYD231, a region downstream of *amyE* into *B. subtilis* 168 using oTG215 and oTG216, and selecting for for growth on LB plates containing 100 µg ml-1 spectinomycin. The confirmed clone was then transformed with ScaI-linearized plasmid *yhdG::P_hy_-yisK E30A (phleo)*, selecting for for growth on LB plates containing 400 µg ml-1 phleomycin.

**BTG042** [△*ponA::erm*] was created by transforming BJH004 with genomic DNA from BYD172 [△*ponA::erm*] selecting for growth on LB plates containing 1 µg/ml erythromycin (erm) plus 25 µg/ml lincomycin (MLS) and 10.0 mM MgCl_2_.

In the process of generating the strains for the experiments in Fig 3, we noted that the Δ*ponA*::*erm* strain we made (designated as Δ*ponA*) grew slightly slower than the strain obtained from the *Bacillus* Genetic Stock Center (designated as Δ*ponA\**)(Fig S5A). In addition, we noted that even when grown in the presence of excess MgCl_2_, the slower growing Δ*ponA* cells were more irregular in shape than the Δ*ponA\** cells (Fig S5C). Whole genome sequencing revealed Δ*ponA\** possessed a mutation in *walH* (CAGèTAG; Q242stop), a gene encoding a negative regulator of the WalRK envelope stress response (7, 8). Consistent with loss of *walH* function, Δ*ponA\** cells exhibited strong activation of a WalR-activated promoter (P*_yocH_*) and enhanced repression of a WalR-repressed promoter (P*_iseA_*) compared to a wildtype control (Fig S5B and S5C). In contrast, the slower growing Δ*ponA* strain used in the YisK experiments exhibited only modest and heterogeneous activation of P*_yocH_* (Fig S5B and S5C) and repression of P*_iseA_* (Fig S5D) relative to wildtype. These results suggest that in the absence of *ponA*, a constitutively active WalRK response is favorable to growth.

**BTG068** [*amyE::P_hy_-opt rbs-yisK_SW_-GFP (spec) (E111-GFP-A112)*] was created by transforming *B. subtilis* 168 with a linear Gibson assembly product encoding three fragments including a region upstream of *amyE* with a spectinomycin cassette using oTG109 and oTG110, the *P_hy_-opt rbs-yisK_SW_-GFP (spec) E111-GFP-A112* using oEA35 and oTG10, oEA36 and oTG009, oTG001 and oTG002, a region downstream of *amyE* into *B. subtilis* 168 using oTG215 and oTG216, and selecting for for growth on LB plates containing 100 µg ml-1 spectinomycin. The detailed protocol is mentioned in the lab previous work (1).

### Allelic replacement of wild-type *yisK* with *yisK-GFP*

The native *yisK_SW_-GFP* were generated through allelic replacement. Briefly, *yisK_SW_-GFP* gene was generated using overlap extension PCR and cloned into the vector pMiniMad2. The plasmid was then purified from TG-1 E. coli and used to transform *B. subtilis* 168. Single-crossover integration was selected for by plating cells at 37°C in the presence of erythromycin (1 µg/ml) and lincomycin (25 µg/ml). Six independent colonies were inoculated into six independent 3 ml LB cultures and grown overnight at room temperature in a rotary drum set at 60 rpm. The cultures were back-diluted 150X in fresh LB, and grown 8 hr at room temperature. 100 μl of a 10^−5^ dilution of each culture was plated on 6 independent LB plates and incubated overnight at 37°C. Ten single colonies from each plate were patched on an LB plate and an LB plate supplemented with erythromycin (1 µg/ml) and lincomycin (25 µg/ml). After streaking for isolated colonies, genomic DNA was collected from several antibiotic sensitive colonies obtained from independent cultures. The YisK region was then PCR amplified and strains carrying the desired mutation were identified by sequencing with primers oAS334 and oAS337.

**BTG078** [*yisK::yisK_SW_-GFP*] was created by transforming *B. subtilis* 168 with a plamid pTG004 following alleic replacement process.

**BTG079** [*yisK::yisK_SW_-GFP*, △*ftsEX::tet*] was created by transforming BTG078 with genomic DNA from BAS319 (△*ftsEX::tet)* and selecting for growth on LB plates containing 10 µg/ml tetracycline.

**BTG081** [*yisK::yisK_SW_-GFP*, △*pona::erm*] was created by transforming BTG78 with genomic DNA from BYD172 [△*ponA::erm*] selecting for growth on LB plates containing 1 µg/ml erythromycin (erm) plus 25 µg/ml lincomycin (MLS) and 10.0 mM MgCl_2_.

**BTG082** [*yisK::yisK_SW_-GFP*, △*ponA::erm, △mbl (kan linked)*] was created by transforming BTG81 with genomic DNA from BYD258 [△*mbl (Kan linked)*](9) and selecting for growth on LB plates containing 10 µg/ml kanamycin and 10mM MgCl_2_.

**BTG088** [*yisK::yisK_SW_-GFP E30A*] was created by transforming *B. subtilis* 168 with a plamid pTG006 following alleic replacement process.

**BTG160** [*amyE::P_hy_-yisK_SW_-GFP E148A E150A (spec)*] was created by transforming *B. subtilis* 168 with a linear gibson assembly product encoding three fragments including a region upstream of *amyE* with a spectinomycin cassette using oTG109 and oTG110, the *P_hy_-opt rbs-yisK (spec) E111-GFP-A112 E148A E150A* using oEA35 and oTG10, oEA36 and oTG009, oTG001 and oTG002, a region downstream of *amyE* into *B. subtilis* 168 using oTG215 and oTG216, and selecting for for growth on LB plates containing 100 µg ml-1 spectinomycin.

**BTG278** [*amyE ::P_yocH_-optRBS-venus (cat)*] was created by transforming BJH004 with genomic DNA from bGD300. *amyE ::P_yocH_-optRBS-venus (cat)* was selected for by growth on LB plates containing 5 µg/ml chloramphenicol (cat).

**BTG279** [*amyE ::P_iseA_-optRBS-venus (cat)*] was created by transforming BJH004 with genomic DNA from BGD110 [*amyE ::P_iseA_-optRBS-venus (cat)*] and selecting for growth on LB plates containing 5 µg/ml chloramphenicol (cat).

**BTG468** [△*ponA::erm (suppressor)*, *amyE ::P_yocH_-optRBS-venus (cat)*] was created by transforming BYD172 with genomic DNA from bGD300. *amyE ::P_yocH_-optRBS-venus (cat)*] was selected for by growth on LB plates containing 5 µg/ml chloramphenicol (cat).

**BTG469** [△*ponA*::erm (suppressor), amyE ::P_iseA_-optRBS-venus (cat)*] was created by transforming BYD172 with genomic DNA from bGD110. *amyE ::P_iseA_-optRBS-venus (cat)*] was selected for by growth on LB plates containing 5 µg/ml chloramphenicol (cat).

**BTG470** [△*ponA*::erm*, *amyE ::P_yocH_-optRBS-venus (cat)*] was created by transforming BTG42 with genomic DNA from bGD300. *amyE ::P_yocH_-optRBS-venus (cat)* was selected for by growth on LB plates containing 5 µg/ml chloramphenicol (cat).

**BTG471** [△*ponA::erm, amyE ::P_iseA_-optRBS-venus (cat)*] was created by transforming BTG042 with genomic DNA from bGD110. *amyE ::P_iseA_-optRBS-venus (cat)* was selected for by growth on LB plates containing 5 µg/ml chloramphenicol (cat).

**BTG607** [*amyE::P_hy_ -yisK E30A (spec), tryptophan protrophic*] was created by transforming BTG169 tryptophan protrophic strian with genomic DNA from *amyE::P_hy_ -yisK E30A (spec)* [*amyE::P_hy_ -yisK E30A (spec)*] selecting for growth on LB plates containing 100 µg ml-1 spectinomycin.

**BTG608** [*amyE::P_hy_ -yisK (spec), tryptophan protrophic*] was created by transforming BTG169 tryptophan protrophic strian with genomic DNA from BAS41 [*amyE::P_hy_ -yisK (spec)*] selecting for growth on LB plates containing 100 µg ml-1 spectinomycin.

**BTG895** [*amyE::P_hy_ -yisK (spec)*, △*ftsEX::tet*] was created by transforming BAS41 with genomic DNA from BAS319 [△*ftsEX::tet*] selecting for growth on LB plates containing 10 µg/ml tetracycline.

**BYD587** [*amyE::P_hy_ -yisK (spec)*] was created by transforming *B. subtilis* 168 with a linear Gibson assembly product encoding three fragments including a region upstream of *amyE* with a spectinomycin cassette using oTG109 and oTG110, the *P_hy_-yisK WT* using oEA35 and oEA36 amplified from qYD104, a region downstream of *amyE* into *B. subtilis* 168 using oTG215 and oTG216 and selecting for for growth on LB plates containing 100 µg ml-1 spectinomycin. The detailed protocol is mentioned in the lab previous work (1).

**BYD588** [*amyE::P_hy_ -yisK E30A (spec)*] was created by transforming *B. subtilis* 168 with a linear Gibson assembly product encoding three fragments including a region upstream of *amyE* with a spectinomycin cassette using oTG109 and oTG110, the *P_hy_-yisK E30A* using oEA35 and oEA36 amplified from qYD497, a region downstream of *amyE* into *B. subtilis* 168 using oTG215 and oTG216 and selecting for for growth on LB plates containing 100 µg ml-1 spectinomycin. The detailed protocol is mentioned in the lab previous work (1).

### Plasmid construction

**pTG004** was generated with overlap extension PCR. The “UP” product was amplified from *B. subtilis 168* genomic DNA with primer pair oAS334/oAS335. The “DOWN” product was amplified from BTG68 with primer pair oAS336/ oAS337. The two PCR products were used as template for overlap extension PCR with primer pair oAS334/oAS337. The amplified fragment was cut with EcoRI and BamHI and cloned into pMiniMAD2 cut with the same enzymes.

**pTG006** was generated with overlap extension PCR. The “UP” product was amplified from *B. subtilis 168* genomic DNA with primer pair oAS334/oAS335. The “DOWN” product was amplified from BTG87 with primer pair oAS336/oAS337. The two PCR products were used as template for overlap extension PCR with primer pair oAS334/oAS337. The amplified fragment was cut with EcoRI and BamHI and cloned into pMiniMAD2 cut with the same enzymes.

**pYD206b** was generated by cloning PCR product from oYD262 and oYD263 amplification of *B. subtilis 168* genomic DNA into pKNT25 (EcoRI/BamHI).

**pYD207b** was generated by cloning PCR product from oYD262 and oYD263 amplification of *B. subtilis 168* genomic DNA into pKT25 (EcoRI/BamHI).

**pYD216** was generated by cloning PCR product from oYD298 and oYD299 amplification of *B. subtilis 168* genomic DNA into pCH364 (EcoRI/BamHI).

**pYD218** was generated by cloning PCR product from oYD298 and oYD299 amplification of *B. subtilis 168* genomic DNA into pKNT25 (EcoRI/BamHI).

**pYD104** was generated by cloning PCR product from oYD407 and oYD409 amplification of *B. subtilis 168* genomic DNA into pET24b (NheI/XhoI).

**pYD220** was generated by cloning PCR product from oYD475 and oYD476 amplification of *B. subtilis 168* genomic DNA into pKNT25 (EcoRI/BamHI).

**pYD221** was generated by cloning PCR product from oYD479 and oYD480 amplification of *B. subtilis 168* genomic DNA into pKNT25 (EcoRI/BamHI).

**pYD227** was generated by cloning PCR product from oYD475 and oYD480 amplification of *B. subtilis 168* genomic DNA into pKNT25 (EcoRI/BamHI).

**pYD228** was generated with overlap extension PCR. The “UP” product was amplified from *B. subtilis 168* genomic DNA with primer pair oYD407/oYD546. The “DOWN” product was amplified from pJW006 with primer pair oYD409/ oYD545. The two PCR products were used as template for overlap extension PCR with primer pair oYD407/oYD409. The amplified fragment was cut with NheI and XhoI and cloned into pET24b cut with the same enzymes.

**pYD229** was generated with overlap extension PCR. The “UP” product was amplified from *B. subtilis 168* genomic DNA with primer pair oYD298/oYD578. The “DOWN” product was amplified from pJW006 with primer pair oYD299/ oYD577. The two PCR products were used as template for overlap extension PCR with primer pair oYD298/oYD299. The amplified fragment was cut with EcoRI and BamHI and cloned into pKNT25 cut with the same enzymes.

**pYD230** was generated with overlap extension PCR. The “UP” product was amplified from *B. subtilis 168* genomic DNA with primer pair oYD298/oYD578. The “DOWN” product was amplified from pJW006 with primer pair oYD299/ oYD577. The two PCR products were used as template for overlap extension PCR with primer pair oYD298/oYD299. The amplified fragment was cut with EcoRI and BamHI and cloned into pCH364 cut with the same enzymes.

**pYD231** was generated by cloning PCR product from oYD407 and oYD409 amplification of pYD229 into pET24b (NheI/XhoI).

## 1 Overall quality at a glance

The following experimental techniques were used to determine the structure:

*X-RAY DIFFRACTION*

The reported resolution of this entry is 2.26 Å.

Percentile scores (ranging between 0-100) for global validation metrics of the entry are shown in the following graphic. The table shows the number of entries on which the scores are based.

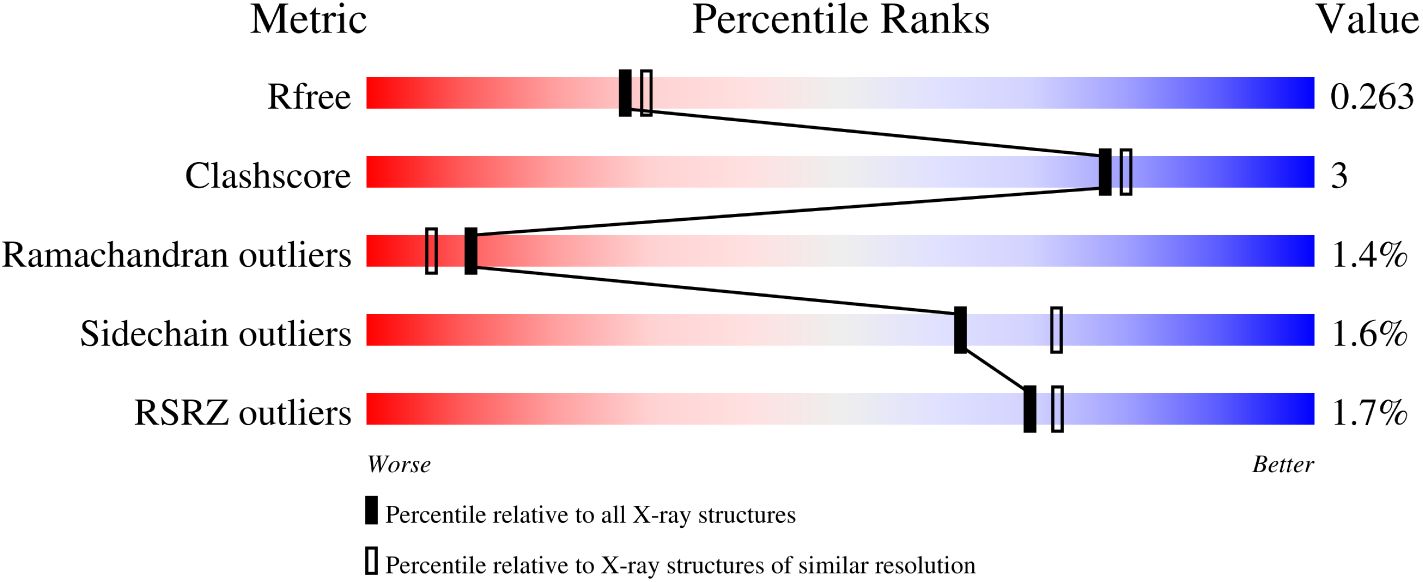

| Metric | Whole archive<br>(#Entries) | Similar resolution<br>(#Entries, resolution range(Å)) |
| --- | --- | --- |
| $R_{free}$ | 130704 | 1377 (2.26-2.26) |
| Clashscore | 141614 | 1487 (2.26-2.26) |
| Ramachandran outliers | 138981 | 1449 (2.26-2.26) |
| Sidechain outliers | 138945 | 1450 (2.26-2.26) |
| RSRZ outliers | 127900 | 1356 (2.26-2.26) |

The table below summarises the geometric issues observed across the polymeric chains and their fit to the electron density. The red, orange, yellow and green segments of the lower bar indicate the fraction of residues that contain outliers for *>*=3, 2, 1 and 0 types of geometric quality criteria respectively. A grey segment represents the fraction of residues that are not modelled. The numeric value for each fraction is indicated below the corresponding segment, with a dot representing fractions *<*=5% The upper red bar (where present) indicates the fraction of residues that have poor fit to the electron density. The numeric value is given above the bar.

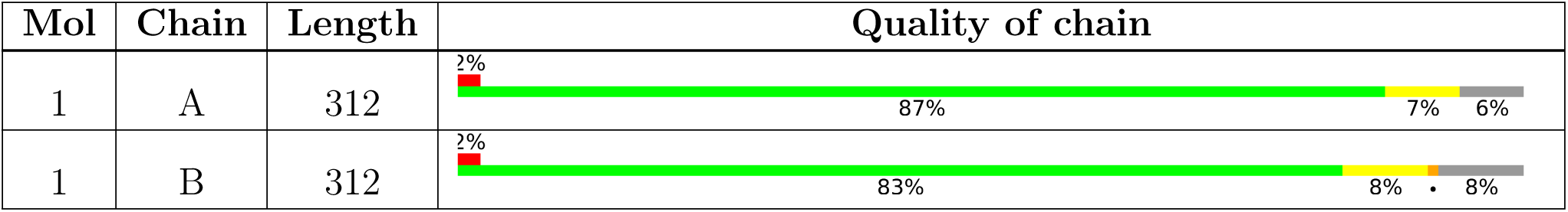

## 2 Entry composition

There are 3 unique types of molecules in this entry. The entry contains 4552 atoms, of which 0 are hydrogens and 0 are deuteriums.

In the tables below, the ZeroOcc column contains the number of atoms modelled with zero occupancy, the AltConf column contains the number of residues with at least one atom in alternate conformation and the Trace column contains the number of residues modelled with at most 2 atoms.

- Molecule 1 is a protein called Fumarylacetoacetate hydrolase family protein.

| Mol | Chain | Residues | Atoms |  |  |  |  | ZeroOcc | AltConf | Trace |
| --- | --- | --- | --- | --- | --- | --- | --- | --- | --- | --- |
| 1 | A | 293 | Total | C | N | O | S | 0 | 0 | 0 |
|  |  |  | 2269 | 1440 | 382 | 436 | 11 |  |  |  |
| 1 | B | 288 | Total | C | N | O | S | 0 | 0 | 0 |
|  |  |  | 2231 | 1420 | 374 | 426 | 11 |  |  |  |

There are 22 discrepancies between the modelled and reference sequences:

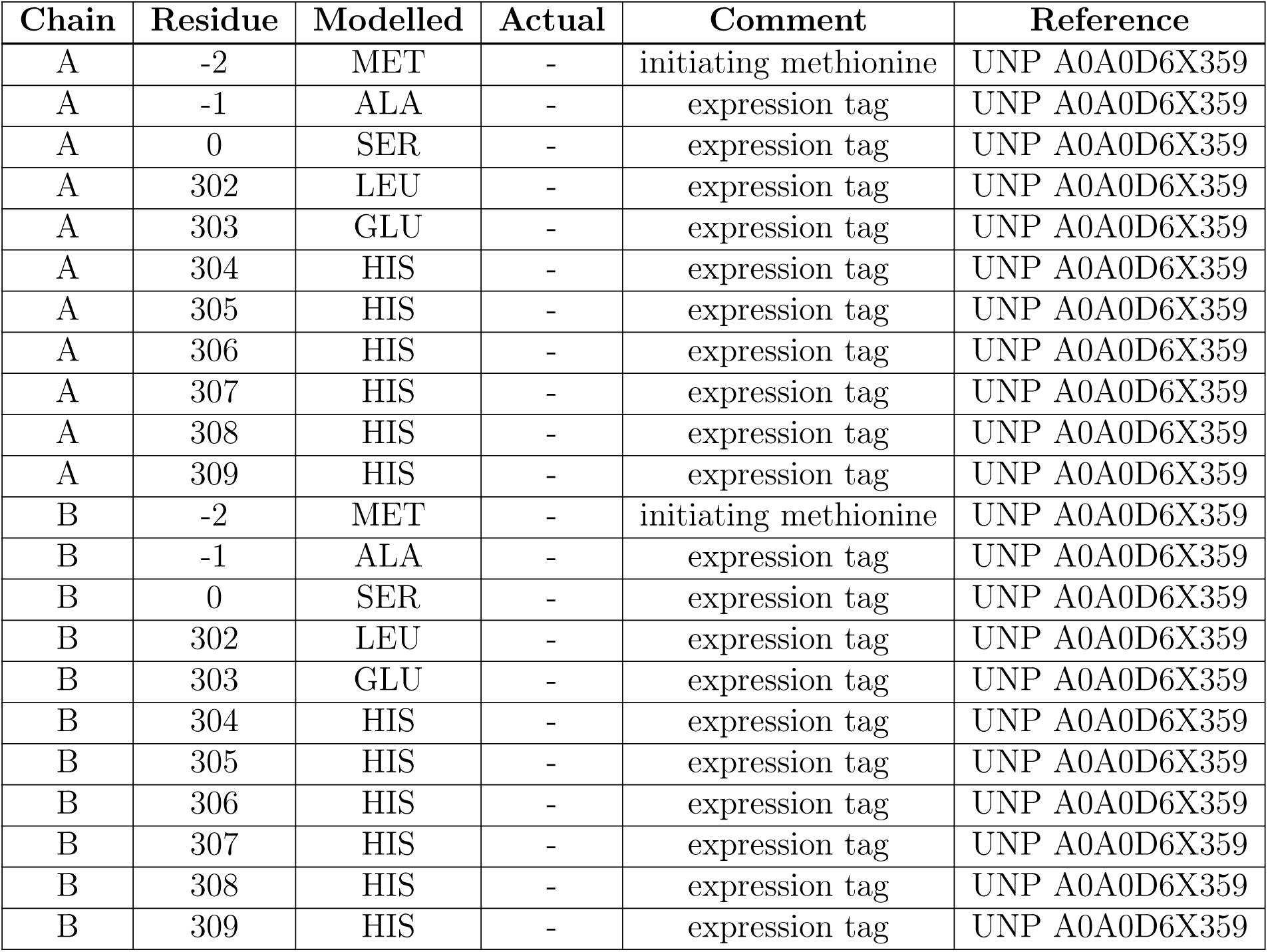

- Molecule 2 is MANGANESE (II) ION (three-letter code: MN) (formula: Mn).

| Mol | Chain | Residues | Atoms |  | ZeroOcc | AltConf |
| --- | --- | --- | --- | --- | --- | --- |
| 2 | A | 1 | Total<br>1 | Mn<br>1 | 0 | 0 |
| 2 | B | 1 | Total<br>1 | Mn<br>1 | 0 | 0 |

- Molecule 3 is water.

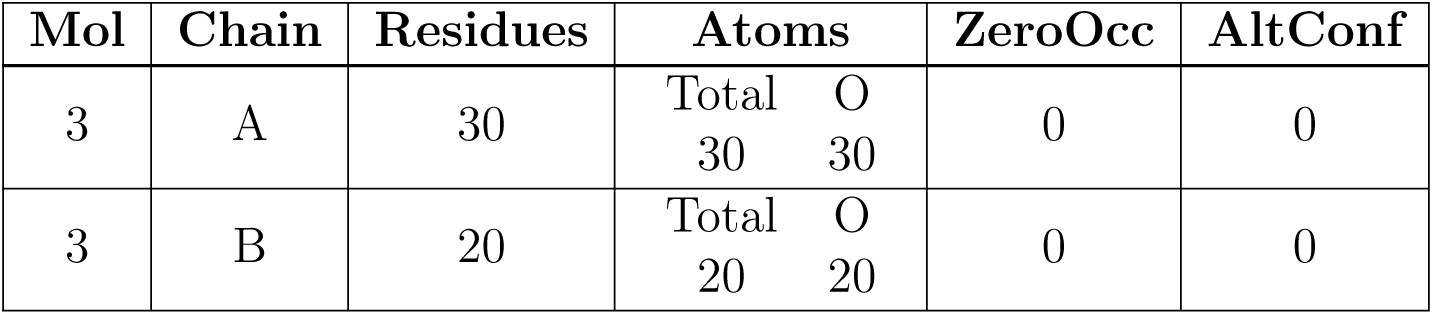

| Mol | Chain | Residues | Atoms |  | ZeroOcc | AltConf |
| --- | --- | --- | --- | --- | --- | --- |
| 3 | A | 30 | Total<br>30 | O<br>30 | 0 | 0 |
| 3 | B | 20 | Total<br>20 | O<br>20 | 0 | 0 |

## 3 Residue-property plots

These plots are drawn for all protein, RNA, DNA and oligosaccharide chains in the entry. The first graphic for a chain summarises the proportions of the various outlier classes displayed in the second graphic. The second graphic shows the sequence view annotated by issues in geometry and electron density. Residues are color-coded according to the number of geometric quality criteria for which they contain at least one outlier: green = 0, yellow = 1, orange = 2 and red = 3 or more. A red dot above a residue indicates a poor fit to the electron density (RSRZ *>* 2). Stretches of 2 or more consecutive residues without any outlier are shown as a green connector. Residues present in the sample, but not in the model, are shown in grey.

- Molecule 1: Fumarylacetoacetate hydrolase family protein

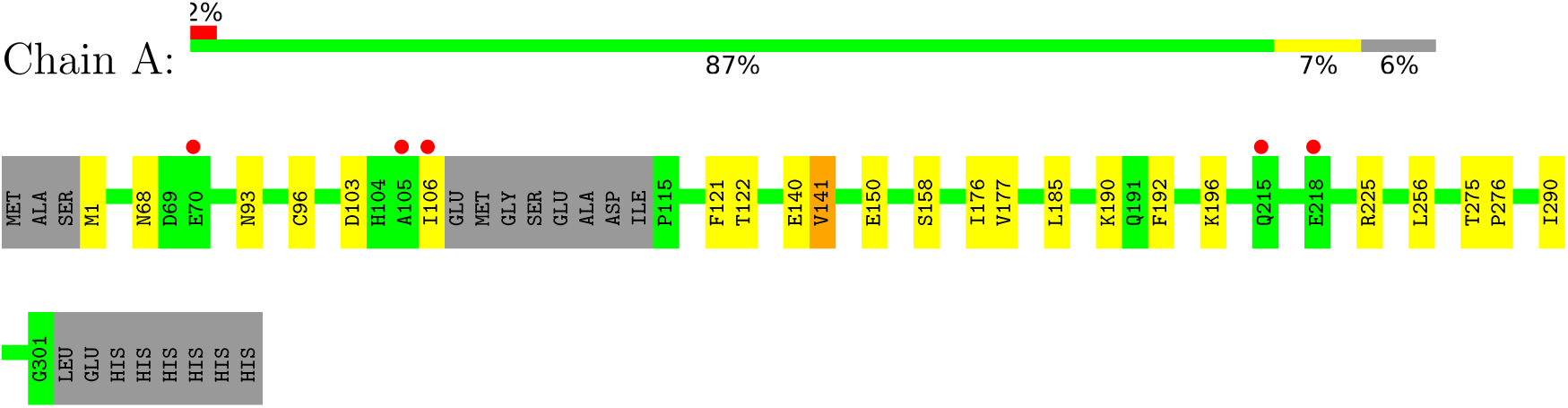

- Molecule 1: Fumarylacetoacetate hydrolase family protein

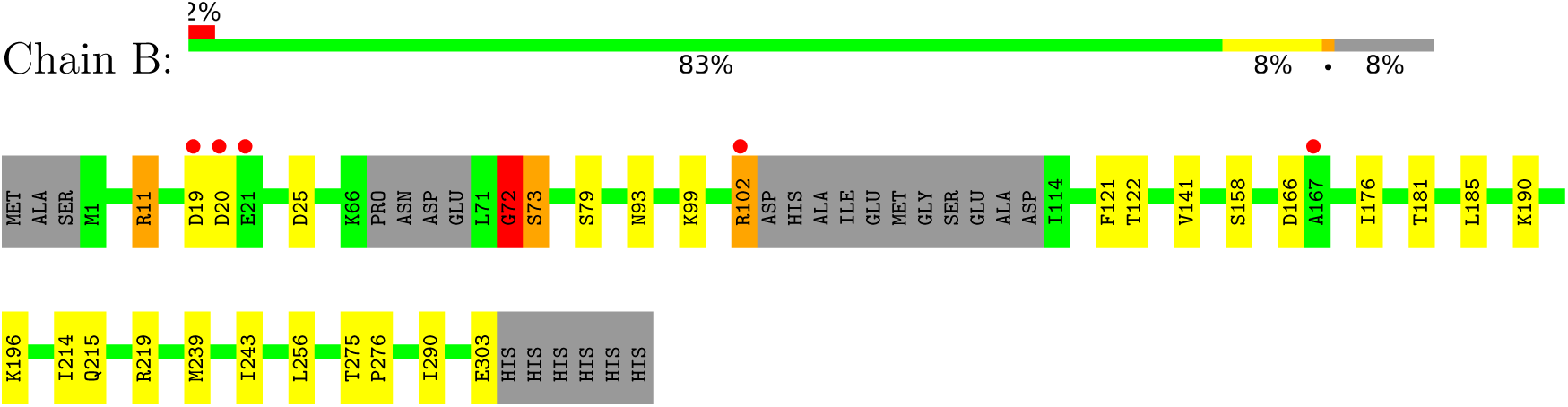

## 4 Data and refinement statistics

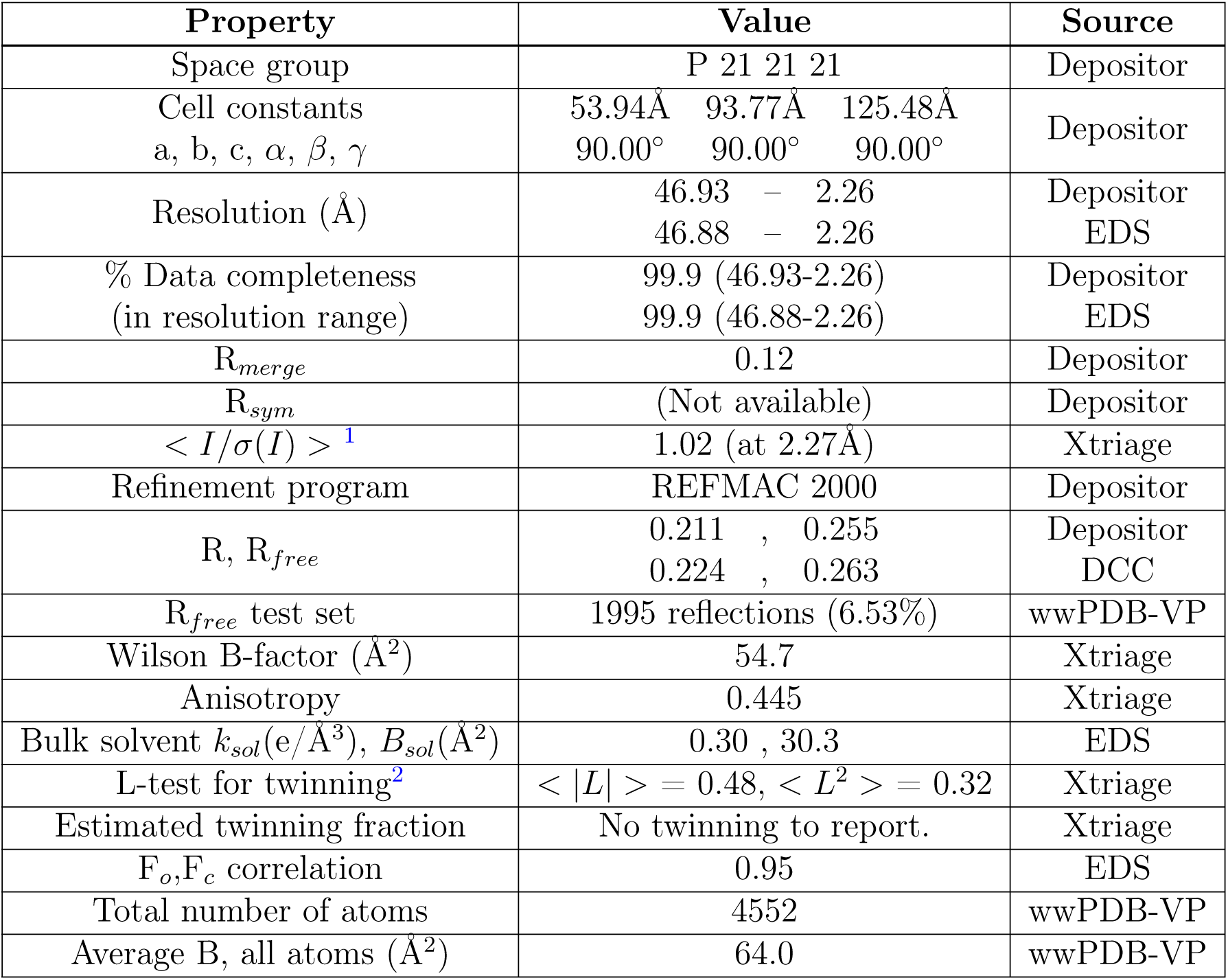

Xtriage’s analysis on translational NCS is as follows: *The largest off-origin peak in the Patterson function is 5.08% of the height of the origin peak. No significant pseudotranslation is detected*.

## 5 Model quality

### 5.1 Standard geometry

Bond lengths and bond angles in the following residue types are not validated in this section: MN

The Z score for a bond length (or angle) is the number of standard deviations the observed value is removed from the expected value. A bond length (or angle) with *|Z| >* 5 is considered an outlier worth inspection. RMSZ is the root-mean-square of all Z scores of the bond lengths (or angles).

| Mol | Chain | Bond lengths |  | Bond angles |  |
| --- | --- | --- | --- | --- | --- |
|  |  | RMSZ | #\|Z\| >5 | RMSZ | #\|Z\| >5 |
| 1 | A | 0.33 | 0/2312 | 0.57 | 0/3117 |
| 1 | B | 0.33 | 0/2271 | 0.58 | 0/3059 |
| All | All | 0.33 | 0/4583 | 0.57 | 0/6176 |

Chiral center outliers are detected by calculating the chiral volume of a chiral center and verifying if the center is modelled as a planar moiety or with the opposite hand.A planarity outlier is detected by checking planarity of atoms in a peptide group, atoms in a mainchain group or atoms of a sidechain that are expected to be planar.

| Mol | Chain | #Chirality outliers | #Planarity outliers |
| --- | --- | --- | --- |
| 1 | A | 0 | 2 |
| 1 | B | 0 | 4 |
| All | All | 0 | 6 |

There are no bond length outliers.

There are no bond angle outliers.

There are no chirality outliers.

All (6) planarity outliers are listed below:

| Mol | Chain | Res | Type | Group |
| --- | --- | --- | --- | --- |
| 1 | A | 225 | ARG | Sidechain |
| 1 | A | 275 | THR | Peptide |
| 1 | B | 102 | ARG | Sidechain |
| 1 | B | 11 | ARG | Sidechain |
| 1 | B | 275 | THR | Peptide |
| 1 | B | 72 | GLY | Peptide |

### 5.2 Too-close contacts

In the following table, the Non-H and H(model) columns list the number of non-hydrogen atoms and hydrogen atoms in the chain respectively. The H(added) column lists the number of hydrogen atoms added and optimized by MolProbity. The Clashes column lists the number of clashes within the asymmetric unit, whereas Symm-Clashes lists symmetry-related clashes.

| Mol | Chain | Non-H | H(model) | H(added) | Clashes | Symm-Clashes |
| --- | --- | --- | --- | --- | --- | --- |
| 1 | A | 2269 | 0 | 2298 | 12 | 0 |
| 1 | B | 2231 | 0 | 2274 | 14 | 0 |
| 2 | A | 1 | 0 | 0 | 0 | 0 |
| 2 | B | 1 | 0 | 0 | 0 | 0 |
| 3 | A | 30 | 0 | 0 | 0 | 0 |
| 3 | B | 20 | 0 | 0 | 0 | 0 |
| All | All | 4552 | 0 | 4572 | 26 | 0 |

The all-atom clashscore is defined as the number of clashes found per 1000 atoms (including hydrogen atoms). The all-atom clashscore for this structure is 3.

All (26) close contacts within the same asymmetric unit are listed below, sorted by their clash magnitude.

| Atom-1 | Atom-2 | Interatomic distance (Å) | Clash overlap (Å) |
| --- | --- | --- | --- |
| 1:B:72:GLY:HA3 | 1:B:73:SER:HB2 | 1.67 | 0.76 |
| 1:B:72:GLY:HA3 | 1:B:73:SER:CB | 2.18 | 0.74 |
| 1:B:11:ARG:NH2 | 1:B:25:ASP:OD2 | 2.27 | 0.65 |
| 1:B:181:THR:HG23 | 1:B:196:LYS:HD3 | 1.86 | 0.58 |
| 1:B:176:ILE:HG12 | 1:B:290:ILE:HD12 | 1.88 | 0.55 |
| 1:B:72:GLY:CA | 1:B:73:SER:CB | 2.86 | 0.54 |
| 1:A:140:GLU:O | 1:A:141:VAL:HB | 2.10 | 0.52 |
| 1:A:140:GLU:O | 1:A:141:VAL:CB | 2.59 | 0.51 |
| 1:B:141:VAL:CG1 | 1:B:185:LEU:HD11 | 2.40 | 0.51 |
| 1:A:150:GLU:HB2 | 1:A:177:VAL:HG23 | 1.94 | 0.49 |
| 1:A:141:VAL:CG1 | 1:A:185:LEU:HD11 | 2.42 | 0.49 |
| 1:B:121:PHE:CE1 | 1:B:196:LYS:HE3 | 2.48 | 0.48 |
| 1:A:176:ILE:HG12 | 1:A:290:ILE:HD12 | 1.97 | 0.47 |
| 1:B:239:MET:HE1 | 1:B:243:ILE:CD1 | 2.46 | 0.46 |
| 1:B:93:ASN:HB3 | 1:B:122:THR:HB | 1.97 | 0.46 |
| 1:A:140:GLU:O | 1:A:141:VAL:HG23 | 2.16 | 0.45 |
| 1:A:93:ASN:HB3 | 1:A:122:THR:HB | 1.98 | 0.45 |
| 1:A:158:SER:HA | 1:A:256:LEU:O | 2.17 | 0.45 |
| 1:A:103:ASP:OD1 | 1:A:106:ILE:HD12 | 2.17 | 0.44 |
| 1:A:141:VAL:HG12 | 1:A:185:LEU:HD11 | 2.00 | 0.44 |
| 1:B:214:ILE:O | 1:B:215:GLN:CG | 2.66 | 0.43 |

| Atom-1 | Atom-2 | Interatomic distance (Å) | Clash overlap (Å) |
| --- | --- | --- | --- |
| 1:A:192:PHE:O | 1:A:196:LYS:HB2 | 2.19 | 0.42 |
| 1:A:96:CYS:HB2 | 1:A:121:PHE:CE1 | 2.54 | 0.42 |
| 1:B:141:VAL:HG12 | 1:B:185:LEU:HD11 | 2.02 | 0.41 |
| 1:B:303:GLU:OE1 | 1:B:303:GLU:N | 2.52 | 0.41 |
| 1:B:158:SER:HA | 1:B:256:LEU:O | 2.21 | 0.40 |

There are no symmetry-related clashes.

### 5.3 Torsion angles

#### 5.3.1 Protein backbone

In the following table, the Percentiles column shows the percent Ramachandran outliers of the chain as a percentile score with respect to all X-ray entries followed by that with respect to entries of similar resolution.

The Analysed column shows the number of residues for which the backbone conformation was analysed, and the total number of residues.

| Mol | Chain | Analysed | Favoured | Allowed | Outliers | Percentiles |  |
| --- | --- | --- | --- | --- | --- | --- | --- |
| 1 | A | 289/312 (93%) | 281 (97%) | 5 (2%) | 3 (1%) | 15 | 13 |
| 1 | B | 282/312 (90%) | 267 (95%) | 10 (4%) | 5 (2%) | 8 | 4 |
| All | All | 571/624 (92%) | 548 (96%) | 15 (3%) | 8 (1%) | 11 | 7 |

All (8) Ramachandran outliers are listed below:

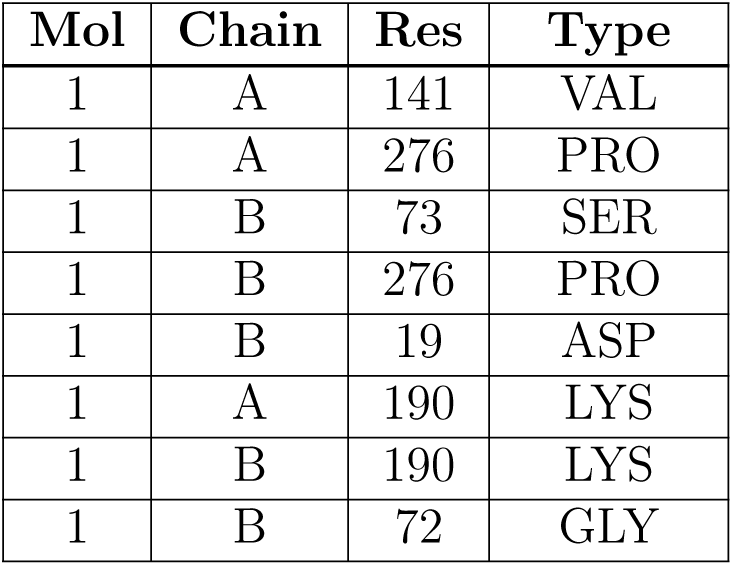

| Mol | Chain | Res | Type |
| --- | --- | --- | --- |
| 1 | A | 141 | VAL |
| 1 | A | 276 | PRO |
| 1 | B | 73 | SER |
| 1 | B | 276 | PRO |
| 1 | B | 19 | ASP |
| 1 | A | 190 | LYS |
| 1 | B | 190 | LYS |
| 1 | B | 72 | GLY |

#### 5.3.2 Protein sidechains

In the following table, the Percentiles column shows the percent sidechain outliers of the chain as a percentile score with respect to all X-ray entries followed by that with respect to entries of similar resolution.

The Analysed column shows the number of residues for which the sidechain conformation was analysed, and the total number of residues.

| Mol | Chain | Analysed | Rotameric | Outliers | Percentiles |  |
| --- | --- | --- | --- | --- | --- | --- |
| 1 | A | 253/269 (94%) | 251 (99%) | 2 (1%) | 81 | 88 |
| 1 | B | 249/269 (93%) | 243 (98%) | 6 (2%) | 49 | 58 |
| All | All | 502/538 (93%) | 494 (98%) | 8 (2%) | 62 | 73 |

All (8) residues with a non-rotameric sidechain are listed below:

| Mol | Chain | Res | Type |
| --- | --- | --- | --- |
| 1 | A | 1 | MET |
| 1 | A | 68 | ASN |
| 1 | B | 20 | ASP |
| 1 | B | 79 | SER |
| 1 | B | 99 | LYS |
| 1 | B | 102 | ARG |
| 1 | B | 166 | ASP |
| 1 | B | 219 | ARG |

Sometimes sidechains can be flipped to improve hydrogen bonding and reduce clashes. All (3) such sidechains are listed below:

| Mol | Chain | Res | Type |
| --- | --- | --- | --- |
| 1 | A | 68 | ASN |
| 1 | A | 84 | HIS |
| 1 | A | 117 | HIS |

#### 5.3.3 RNA

There are no RNA molecules in this entry.

### 5.4 Non-standard residues in protein, DNA, RNA chains

There are no non-standard protein/DNA/RNA residues in this entry.

### 5.5 Carbohydrates

There are no monosaccharides in this entry.

### 5.6 Ligand geometry

Of 2 ligands modelled in this entry, 2 are monoatomic – leaving 0 for Mogul analysis.

There are no bond length outliers.

There are no bond angle outliers.

There are no chirality outliers.

There are no torsion outliers.

There are no ring outliers.

No monomer is involved in short contacts.

### 5.7 Other polymers

There are no such residues in this entry.

### 5.8 Polymer linkage issues

There are no chain breaks in this entry.

## 6 Fit of model and data

### 6.1 Protein, DNA and RNA chains

In the following table, the column labelled ‘#RSRZ*>* 2’ contains the number (and percentage) of RSRZ outliers, followed by percent RSRZ outliers for the chain as percentile scores relative to all X-ray entries and entries of similar resolution. The OWAB column contains the minimum, median, 95*th* percentile and maximum values of the occupancy-weighted average B-factor per residue. The column labelled ‘Q*<* 0.9’ lists the number of (and percentage) of residues with an average occupancy less than 0.9.

| Mol | Chain | Analysed | <RSRZ> | #RSRZ > 2 | OWAB(Å<sup>2</sup>) | Q < 0.9 |
| --- | --- | --- | --- | --- | --- | --- |
| 1 | A | 293/312 (93%) | 0.10 | 5 (1%) 70 73 | 43, 58, 84, 112 | 0 |
| 1 | B | 288/312 (92%) | 0.16 | 5 (1%) 70 73 | 46, 64, 92, 110 | 0 |
| All | All | 581/624 (93%) | 0.13 | 10 (1%) 70 73 | 43, 61, 88, 112 | 0 |

All (10) RSRZ outliers are listed below:

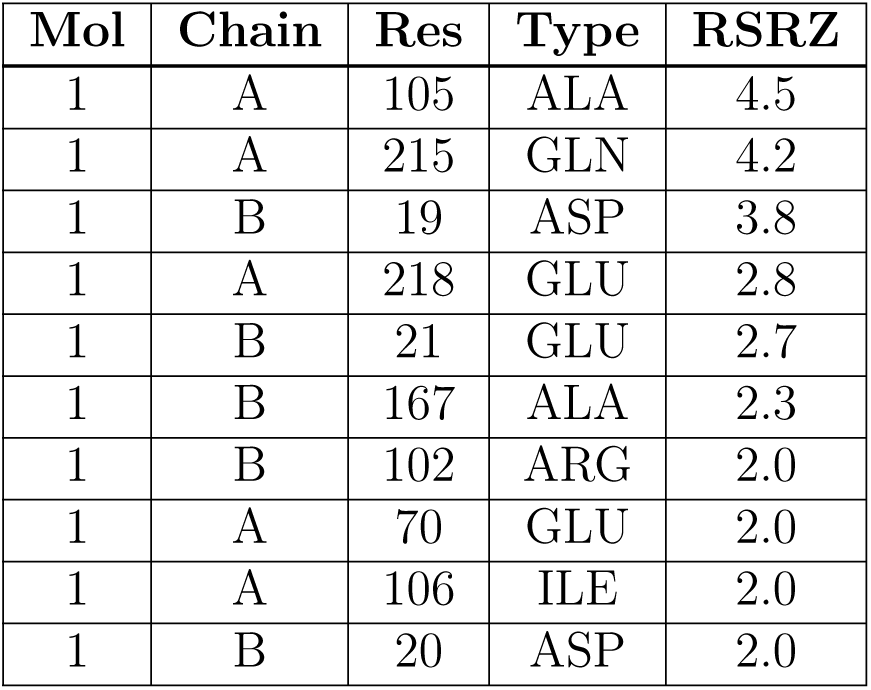

| Mol | Chain | Res | Type | RSRZ |
| --- | --- | --- | --- | --- |
| 1 | A | 105 | ALA | 4.5 |
| 1 | A | 215 | GLN | 4.2 |
| 1 | B | 19 | ASP | 3.8 |
| 1 | A | 218 | GLU | 2.8 |
| 1 | B | 21 | GLU | 2.7 |
| 1 | B | 167 | ALA | 2.3 |
| 1 | B | 102 | ARG | 2.0 |
| 1 | A | 70 | GLU | 2.0 |
| 1 | A | 106 | ILE | 2.0 |
| 1 | B | 20 | ASP | 2.0 |

### 6.2 Non-standard residues in protein, DNA, RNA chains

There are no non-standard protein/DNA/RNA residues in this entry.

### 6.3 Carbohydrates

There are no monosaccharides in this entry.

### 6.4 Ligands

In the following table, the Atoms column lists the number of modelled atoms in the group and the number defined in the chemical component dictionary. The B-factors column lists the minimum, median, 95*th* percentile and maximum values of B factors of atoms in the group. The column labelled ‘Q*<* 0.9’ lists the number of atoms with occupancy less than 0.9.

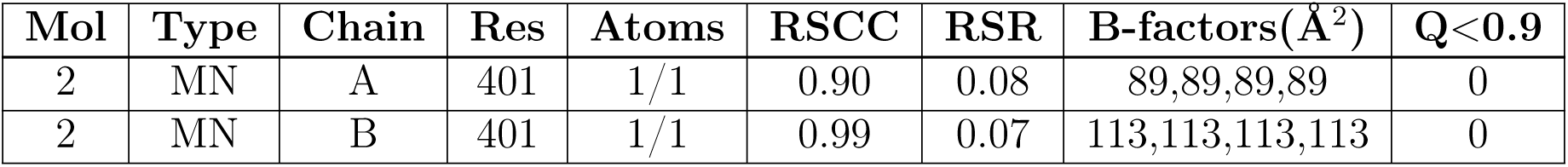

### 6.5 Other polymers

There are no such residues in this entry.

## 1 Overall quality at a glance

The following experimental techniques were used to determine the structure:

### X-RAY DIFFRACTION

The reported resolution of this entry is 2.40 Å.

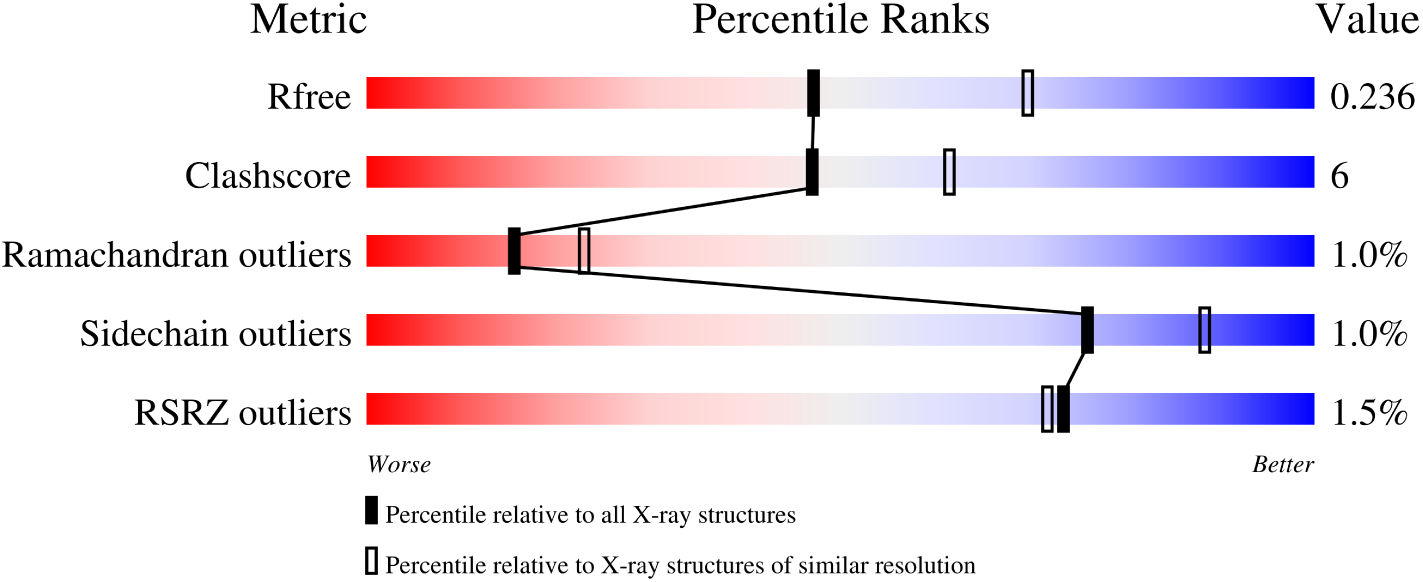

| Metric | Whole archive<br>(#Entries) | Similar resolution<br>(#Entries, resolution range(Å)) |
| --- | --- | --- |
| $R_{free}$ | 130704 | 3907 (2.40-2.40) |
| Clashscore | 141614 | 4398 (2.40-2.40) |
| Ramachandran outliers | 138981 | 4318 (2.40-2.40) |
| Sidechain outliers | 138945 | 4319 (2.40-2.40) |
| RSRZ outliers | 127900 | 3811 (2.40-2.40) |

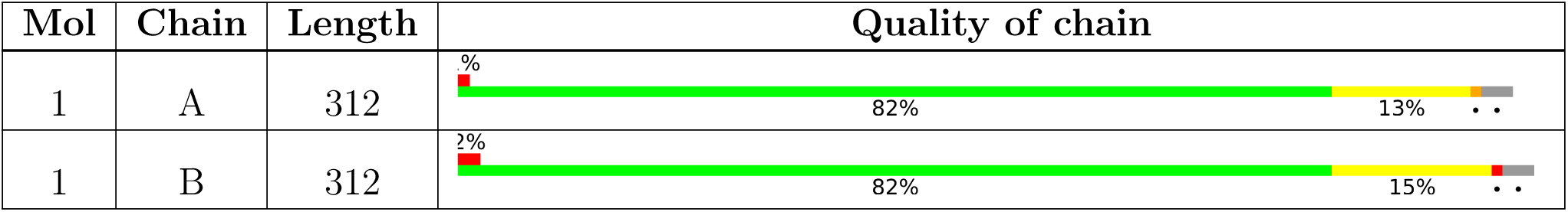

The following table lists non-polymeric compounds, carbohydrate monomers and non-standard residues in protein, DNA, RNA chains that are outliers for geometric or electron-density-fit criteria:

| Mol | Type | Chain | Res | Chirality | Geometry | Clashes | Electron density |
| --- | --- | --- | --- | --- | --- | --- | --- |
| 2 | OXL | B | 401 | - | X | - | - |

## 2 Entry composition

There are 4 unique types of molecules in this entry. The entry contains 4805 atoms, of which 0 are hydrogens and 0 are deuteriums.

- Molecule 1 is a protein called Fumarylacetoacetate hydrolase family protein.

| Mol | Chain | Residues | Atoms |  |  |  |  | ZeroOcc | AltConf | Trace |
| --- | --- | --- | --- | --- | --- | --- | --- | --- | --- | --- |
| 1 | A | 302 | Total | C | N | O | S | 4 | 1 | 0 |
|  |  |  | 2338 | 1482 | 391 | 453 | 12 |  |  |  |
| 1 | B | 303 | Total | C | N | O | S | 0 | 0 | 0 |
|  |  |  | 2336 | 1481 | 392 | 451 | 12 |  |  |  |

There are 22 discrepancies between the modelled and reference sequences:

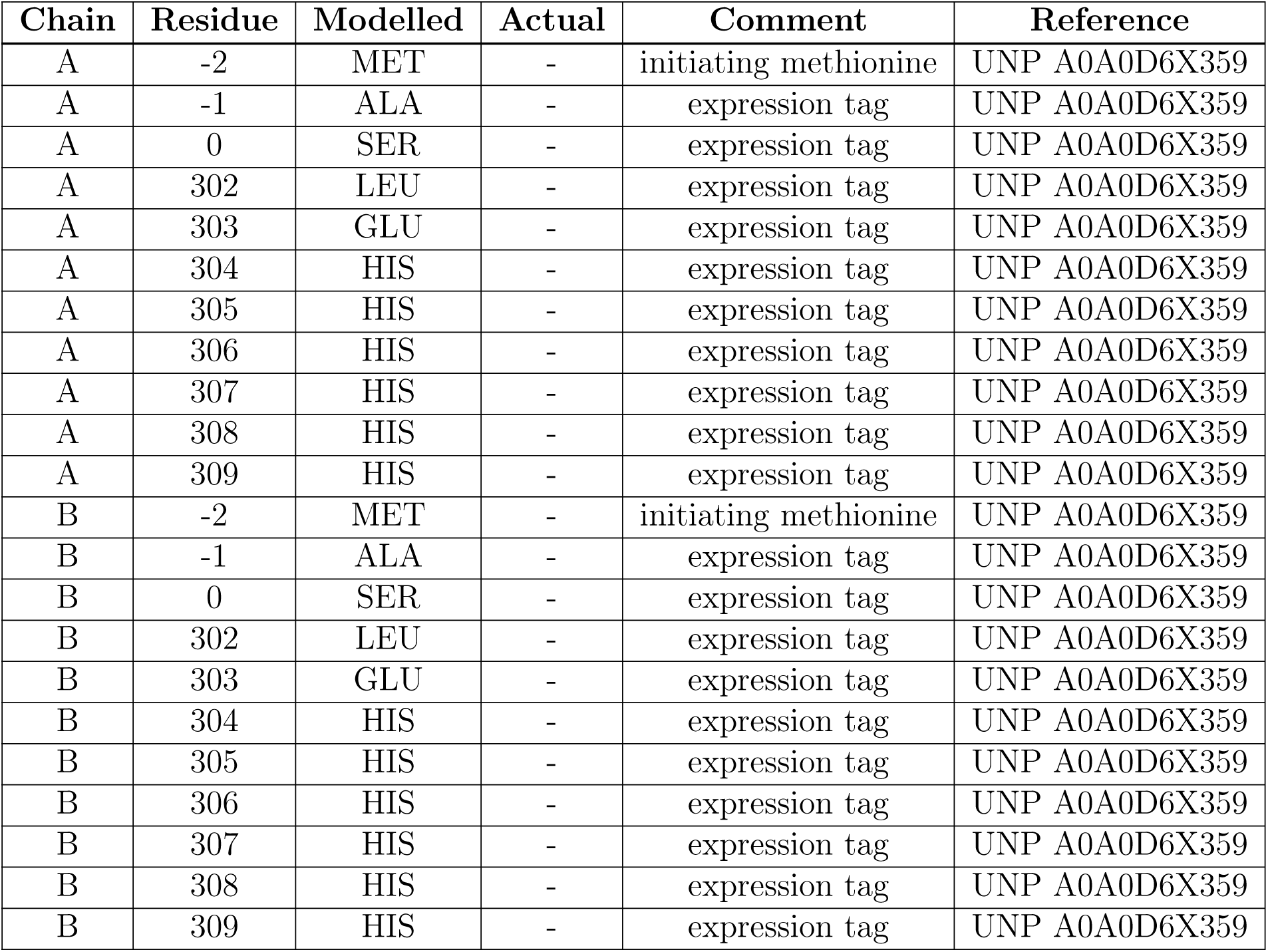

- Molecule 2 is OXALATE ION (three-letter code: OXL) (formula: C_2_O_4_) (labeled as “Ligand of Interest„ by depositor).

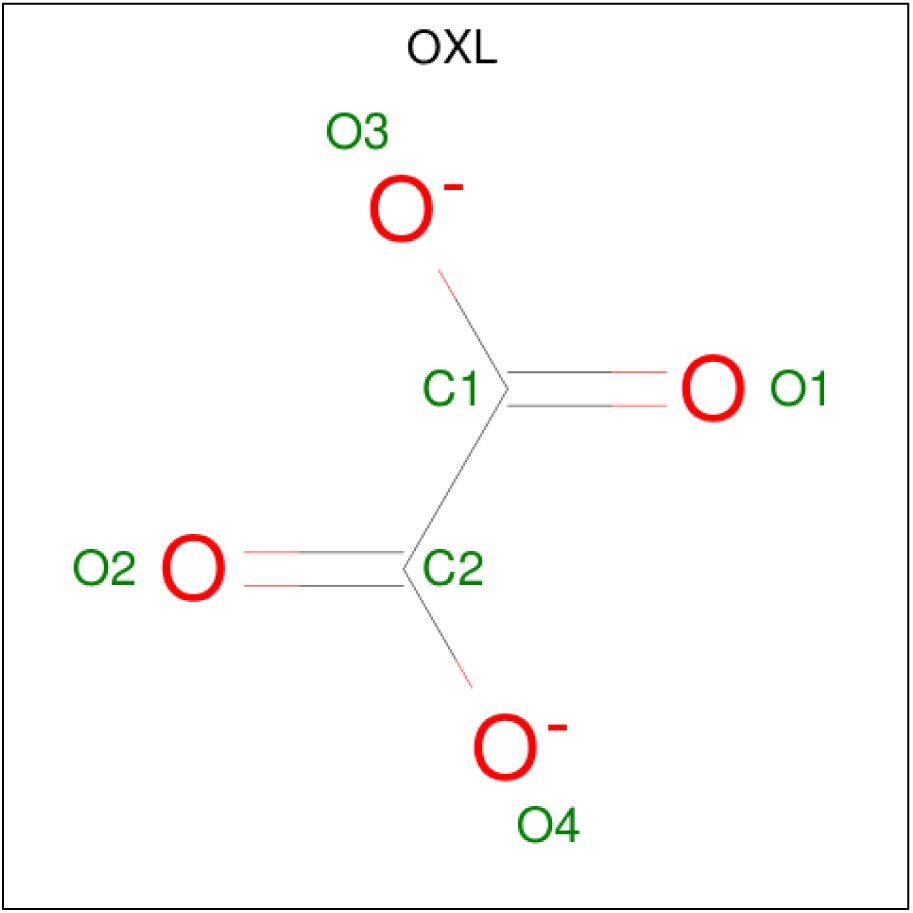

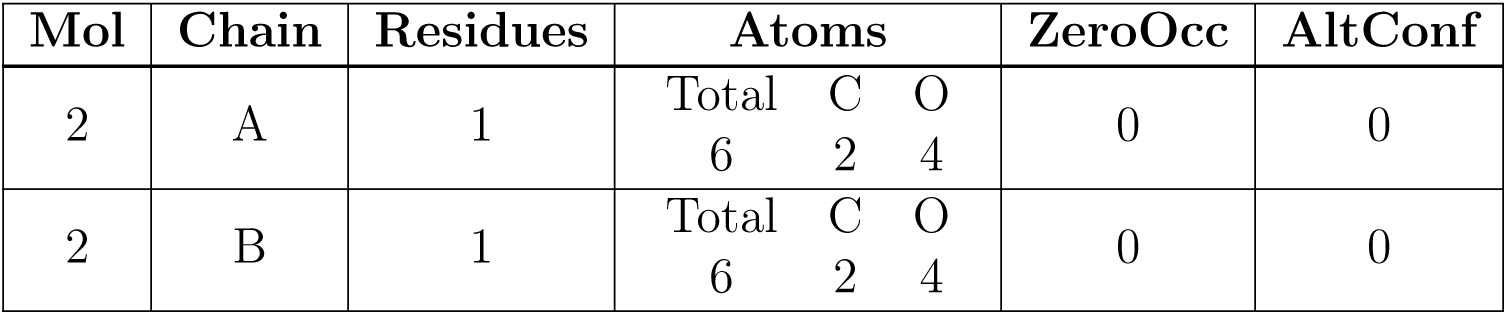

- Molecule 3 is MANGANESE (II) ION (three-letter code: MN) (formula: Mn).

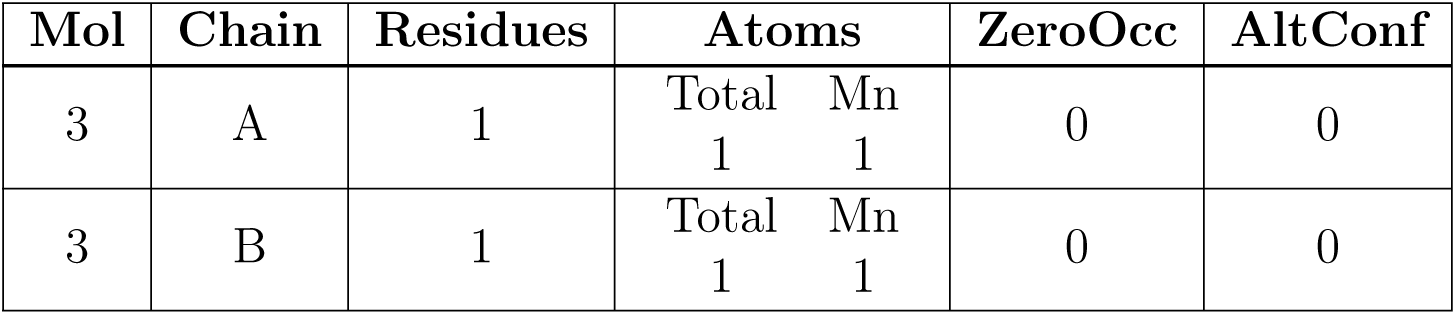

- Molecule 4 is water.

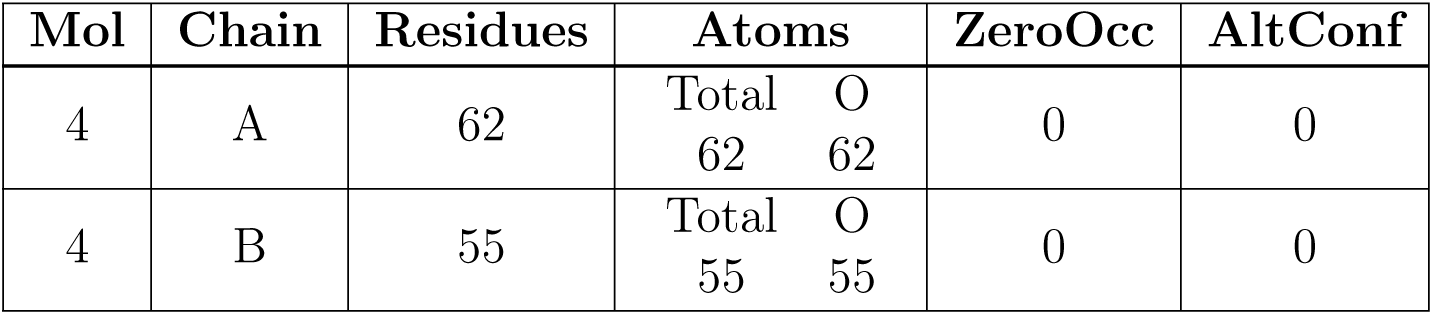

## 3 Residue-property plots

*•* Molecule 1: Fumarylacetoacetate hydrolase family protein

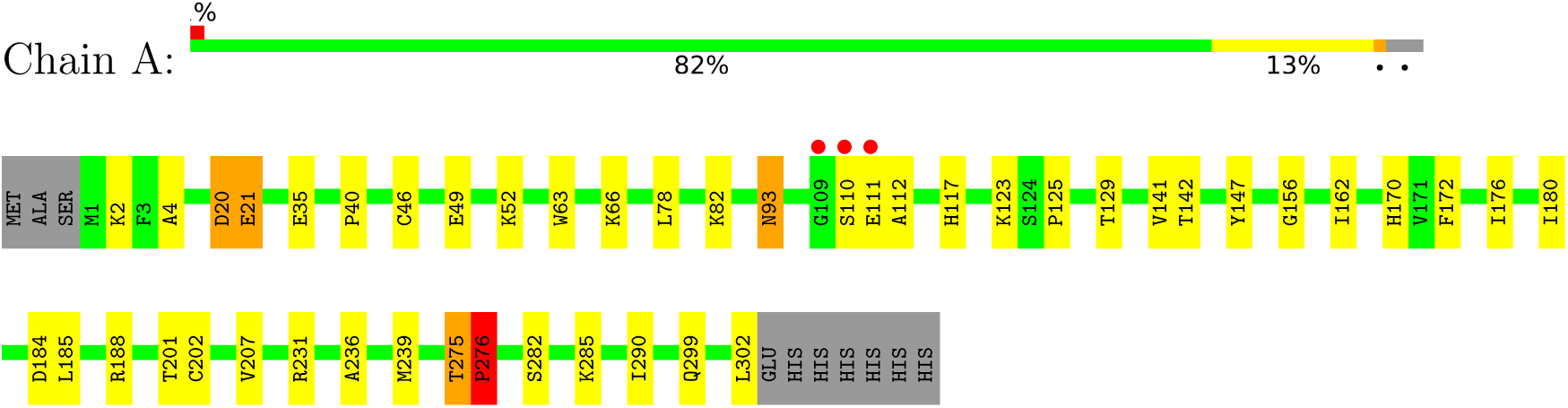

*•* Molecule 1: Fumarylacetoacetate hydrolase family protein

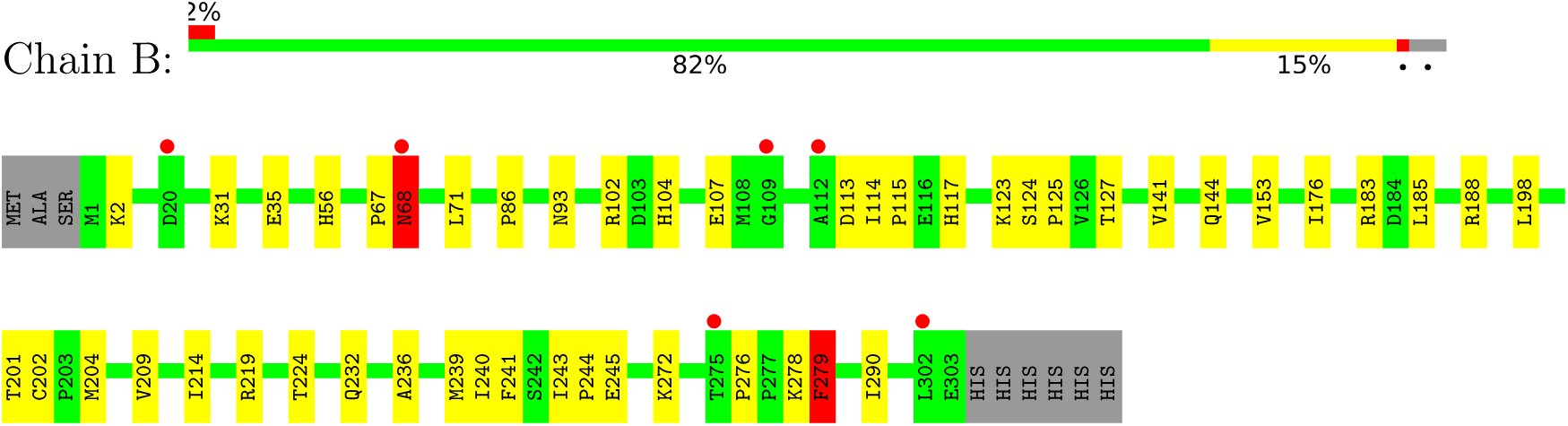

## 4 Data and refinement statistics

| Property | Value | Source |
| --- | --- | --- |
| Space group | P 21 21 21 | Depositor |
| Cell constants<br>a, b, c, $\alpha$, $\beta$, $\gamma$ | 52.34Å 94.84Å 123.94Å<br>90.00° 90.00° 90.00° | Depositor |
| Resolution (Å) | 48.21 – 2.40<br>48.21 – 2.40 | Depositor<br>EDS |
| % Data completeness<br>(in resolution range) | 98.9 (48.21-2.40)<br>98.9 (48.21-2.40) | Depositor<br>EDS |
| $R_{merge}$ | 0.14 | Depositor |
| $R_{sym}$ | (Not available) | Depositor |
| $\langle I/\sigma(I) \rangle$ <sup>1</sup> | 1.46 (at 2.39Å) | Xtriage |
| Refinement program | PHENIX 1.18.2_3874 | Depositor |
| R, $R_{free}$ | 0.189 , 0.236<br>0.189 , 0.236 | Depositor<br>DCC |
| $R_{free}$ test set | 1215 reflections (4.95%) | wwPDB-VP |
| Wilson B-factor (Å<sup>2</sup>) | 45.5 | Xtriage |
| Anisotropy | 0.113 | Xtriage |
| Bulk solvent $k_{sol}$(e/Å<sup>3</sup>), $B_{sol}$(Å<sup>2</sup>) | 0.36 , 45.0 | EDS |
| L-test for twinning<sup>2</sup> | $\langle \vert L\vert \rangle = 0.50$, $\langle L^2 \rangle = 0.33$ | Xtriage |
| Estimated twinning fraction | No twinning to report. | Xtriage |
| $F_o, F_c$ correlation | 0.95 | EDS |
| Total number of atoms | 4805 | wwPDB-VP |
| Average B, all atoms (Å<sup>2</sup>) | 48.0 | wwPDB-VP |

Xtriage’s analysis on translational NCS is as follows: *The largest off-origin peak in the Patterson function is 5.21% of the height of the origin peak. No significant pseudotranslation is detected*.

## 5 Model quality

### 5.1 Standard geometry

Bond lengths and bond angles in the following residue types are not validated in this section: OXL, MN

| Mol | Chain | Bond lengths |  | Bond angles |  |
| --- | --- | --- | --- | --- | --- |
|  |  | RMSZ | #\|Z\| >5 | RMSZ | #\|Z\| >5 |
| 1 | A | 0.40 | 0/2385 | 0.60 | 2/3218 (0.1%) |
| 1 | B | 0.43 | 0/2380 | 0.64 | 4/3211 (0.1%) |
| All | All | 0.41 | 0/4765 | 0.62 | 6/6429 (0.1%) |

There are no bond length outliers.

All (6) bond angle outliers are listed below:

There are no bond length outliers. All (6) bond angle outliers are listed below:
| Mol | Chain | Res | Type | Atoms | Z | Observed(°) | Ideal(°) |
| --- | --- | --- | --- | --- | --- | --- | --- |
| 1 | B | 276 | PRO | N-CA-C | -7.35 | 92.99 | 112.10 |
| 1 | A | 275 | THR | N-CA-C | -7.33 | 91.19 | 111.00 |
| 1 | A | 275 | THR | C-N-CD | -5.79 | 107.87 | 120.60 |
| 1 | B | 279 | PHE | N-CA-C | 5.66 | 126.29 | 111.00 |
| 1 | B | 278 | LYS | N-CA-C | 5.42 | 125.64 | 111.00 |
| 1 | B | 68 | ASN | N-CA-CB | -5.07 | 101.48 | 110.60 |

There are no chirality outliers.

There are no planarity outliers.

### 5.2 Too-close contacts

| Mol | Chain | Non-H | H(model) | H(added) | Clashes | Symm-Clashes |
| --- | --- | --- | --- | --- | --- | --- |
| 1 | A | 2338 | 0 | 2365 | 33 | 0 |
| 1 | B | 2336 | 0 | 2358 | 27 | 0 |
| 2 | A | 6 | 0 | 0 | 0 | 0 |

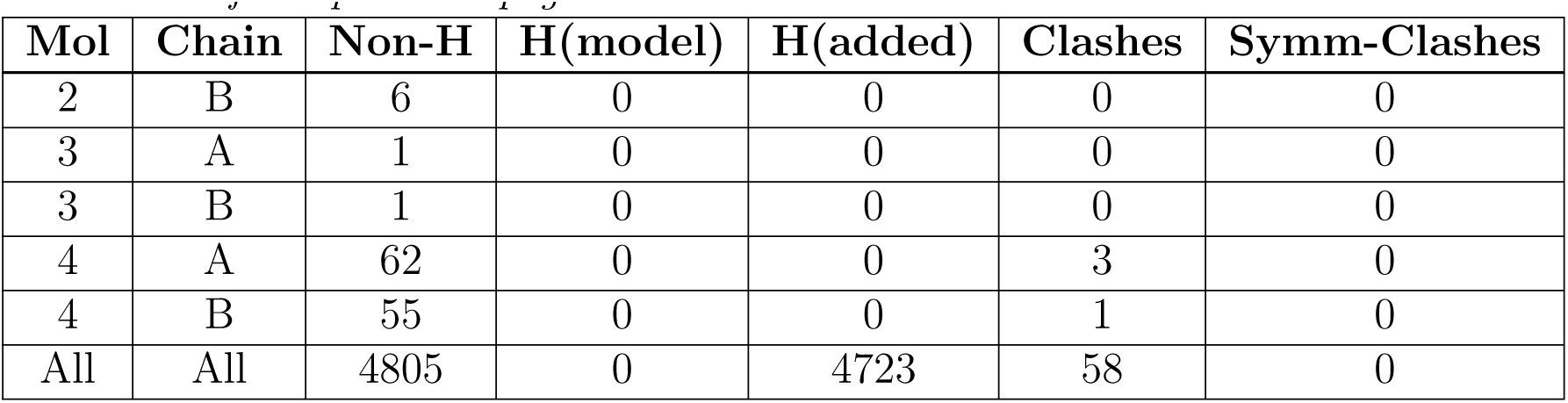

| Mol | Chain | Non-H | H(model) | H(added) | Clashes | Symm-Clashes |
| --- | --- | --- | --- | --- | --- | --- |
| 2 | B | 6 | 0 | 0 | 0 | 0 |
| 3 | A | 1 | 0 | 0 | 0 | 0 |
| 3 | B | 1 | 0 | 0 | 0 | 0 |
| 4 | A | 62 | 0 | 0 | 3 | 0 |
| 4 | B | 55 | 0 | 0 | 1 | 0 |
| All | All | 4805 | 0 | 4723 | 58 | 0 |

The all-atom clashscore is defined as the number of clashes found per 1000 atoms (including hydrogen atoms). The all-atom clashscore for this structure is 6.

All (58) close contacts within the same asymmetric unit are listed below, sorted by their clash magnitude.

| Atom-1 | Atom-2 | Interatomic distance (Å) | Clash overlap (Å) |
| --- | --- | --- | --- |
| 1:A:63:TRP:HA | 1:A:66:LYS:HD3 | 1.63 | 0.81 |
| 1:B:127:THR:HG23 | 1:B:202:CYS:HB3 | 1.75 | 0.68 |
| 1:B:224:THR:HB | 1:B:232:GLN:HB2 | 1.76 | 0.66 |
| 1:B:104:HIS:HA | 1:B:107:GLU:HG2 | 1.80 | 0.62 |
| 1:A:282:SER:HB3 | 1:A:302:LEU:HA | 1.82 | 0.61 |
| 1:B:124:SER:O | 1:B:127:THR:HG22 | 2.02 | 0.60 |
| 1:A:156:GLY:HA3 | 1:A:172:PHE:HB2 | 1.84 | 0.59 |
| 1:A:93:ASN:ND2 | 1:A:125:PRO:HD3 | 2.18 | 0.59 |
| 1:A:2:LYS:NZ | 4:A:501:HOH:O | 2.25 | 0.58 |
| 1:B:123:LYS:HE2 | 1:B:202:CYS:HB2 | 1.84 | 0.58 |
| 1:A:275:THR:HG22 | 1:A:276:PRO:HD2 | 1.85 | 0.58 |
| 1:A:188:ARG:HH11 | 1:A:188:ARG:HG3 | 1.69 | 0.57 |
| 1:A:110:SER:C | 1:A:112:ALA:H | 2.07 | 0.57 |
| 1:B:236:ALA:O | 1:B:239:MET:HG3 | 2.05 | 0.56 |
| 1:A:20:ASP:O | 1:A:21:GLU:HB3 | 2.06 | 0.56 |
| 1:B:115:PRO:HB2 | 1:B:117:HIS:O | 2.08 | 0.53 |
| 1:B:176:ILE:HG12 | 1:B:290:ILE:HD12 | 1.89 | 0.53 |
| 1:A:123:LYS:HE2 | 1:A:202:CYS:HB2 | 1.89 | 0.53 |
| 1:A:82:LYS:HD2 | 4:A:539:HOH:O | 2.10 | 0.52 |
| 1:A:285:LYS:HD3 | 1:A:299:GLN:HG3 | 1.91 | 0.52 |
| 1:B:102:ARG:HG3 | 1:B:240:ILE:HD11 | 1.92 | 0.51 |
| 1:A:156:GLY:CA | 1:A:172:PHE:HB2 | 2.42 | 0.49 |
| 1:B:144:GLN:HB3 | 1:B:183:ARG:HB2 | 1.94 | 0.49 |
| 1:B:56:HIS:CE1 | 4:B:526:HOH:O | 2.65 | 0.48 |
| 1:A:93:ASN:HD22 | 1:A:93:ASN:N | 2.10 | 0.48 |
| 1:A:21:GLU:H | 1:A:78:LEU:HD12 | 1.79 | 0.47 |
| 1:B:68:ASN:HB2 | 1:B:71:LEU:HB2 | 1.98 | 0.46 |
| 1:B:198:LEU:HB2 | 1:B:201:THR:HG21 | 1.97 | 0.46 |

Continued on next page...

| Atom-1 | Atom-2 | Interatomic distance (Å) | Clash overlap (Å) |
| --- | --- | --- | --- |
| 1:B:2:LYS:HG2 | 1:B:209:VAL:HG22 | 1.96 | 0.46 |
| 1:A:147:TYR:OH | 1:A:231:ARG:HD2 | 2.16 | 0.46 |
| 1:A:117:HIS:CG | 1:B:188:ARG:HD2 | 2.51 | 0.45 |
| 1:B:86:PRO:HD2 | 1:B:204:MET:HG2 | 1.98 | 0.45 |
| 1:A:21:GLU:HA | 1:A:78:LEU:HB2 | 1.98 | 0.45 |
| 1:A:236:ALA:O | 1:A:239:MET:HG2 | 2.17 | 0.45 |
| 1:B:141:VAL:HG12 | 1:B:185:LEU:HD11 | 1.99 | 0.45 |
| 1:A:188:ARG:HG3 | 1:A:188:ARG:NH1 | 2.31 | 0.44 |
| 1:A:141:VAL:O | 1:A:185:LEU:HD11 | 2.17 | 0.44 |
| 1:A:184:ASP:N | 1:A:184:ASP:OD1 | 2.51 | 0.44 |
| 1:B:93:ASN:OD1 | 1:B:125:PRO:HD3 | 2.18 | 0.44 |
| 1:B:243:ILE:HB | 1:B:244:PRO:HD3 | 2.00 | 0.44 |
| 1:A:49:GLU:O | 1:A:52:LYS:HE3 | 2.18 | 0.43 |
| 1:A:4:ALA:HB2 | 1:A:207:VAL:HG12 | 2.01 | 0.43 |
| 1:A:35:GLU:HB3 | 4:A:557:HOH:O | 2.18 | 0.43 |
| 1:B:31:LYS:O | 1:B:35:GLU:HA | 2.18 | 0.43 |
| 1:B:153:VAL:HG21 | 1:B:243:ILE:HD12 | 2.01 | 0.42 |
| 1:A:110:SER:C | 1:A:112:ALA:N | 2.72 | 0.42 |
| 1:B:183:ARG:HD2 | 1:B:272:LYS:HD3 | 2.01 | 0.42 |
| 1:B:241:PHE:HA | 1:B:245:GLU:OE2 | 2.19 | 0.42 |
| 1:A:142:THR:HB | 1:A:185:LEU:HD13 | 2.03 | 0.41 |
| 1:A:162:ILE:HD13 | 1:A:170:HIS:CG | 2.56 | 0.41 |
| 1:A:176:ILE:HG12 | 1:A:290:ILE:HD12 | 2.02 | 0.41 |
| 1:B:144:GLN:HG2 | 1:B:279:PHE:CE2 | 2.56 | 0.41 |
| 1:A:93:ASN:ND2 | 1:B:93:ASN:H | 2.19 | 0.41 |
| 1:A:40:PRO:HG2 | 1:A:46:CYS:SG | 2.60 | 0.41 |
| 1:B:113:ASP:O | 1:B:114:ILE:C | 2.58 | 0.40 |
| 1:A:20:ASP:O | 1:A:21:GLU:CB | 2.70 | 0.40 |
| 1:B:214:ILE:HD13 | 1:B:214:ILE:HG21 | 1.85 | 0.40 |
| 1:A:180:ILE:HB | 1:A:201:THR:HB | 2.04 | 0.40 |

There are no symmetry-related clashes.

### 5.3 Torsion angles

#### 5.3.1 Protein backbone

| Mol | Chain | Analysed | Favoured | Allowed | Outliers | Percentiles |  |
| --- | --- | --- | --- | --- | --- | --- | --- |
| 1 | A | 301/312 (96%) | 289 (96%) | 8 (3%) | 4 (1%) | 12 | 17 |
| 1 | B | 301/312 (96%) | 288 (96%) | 11 (4%) | 2 (1%) | 22 | 32 |
| All | All | 602/624 (96%) | 577 (96%) | 19 (3%) | 6 (1%) | 15 | 23 |

All (6) Ramachandran outliers are listed below:

| Mol | Chain | Res | Type |
| --- | --- | --- | --- |
| 1 | A | 111 | GLU |
| 1 | A | 276 | PRO |
| 1 | B | 279 | PHE |
| 1 | A | 21 | GLU |
| 1 | A | 20 | ASP |
| 1 | B | 67 | PRO |

#### 5.3.2 Protein sidechains

| Mol | Chain | Analysed | Rotameric | Outliers | Percentiles |  |
| --- | --- | --- | --- | --- | --- | --- |
| 1 | A | 261/269 (97%) | 257 (98%) | 4 (2%) | 65 | 80 |
| 1 | B | 259/269 (96%) | 257 (99%) | 2 (1%) | 81 | 91 |
| All | All | 520/538 (97%) | 514 (99%) | 6 (1%) | 76 | 85 |

All (6) residues with a non-rotameric sidechain are listed below:

| Mol | Chain | Res | Type |
| --- | --- | --- | --- |
| 1 | A | 93 | ASN |
| 1 | A | 129[A] | THR |
| 1 | A | 129[B] | THR |
| 1 | A | 276 | PRO |
| 1 | B | 68 | ASN |
| 1 | B | 219 | ARG |

Sometimes sidechains can be flipped to improve hydrogen bonding and reduce clashes. All (1) such sidechains are listed below:

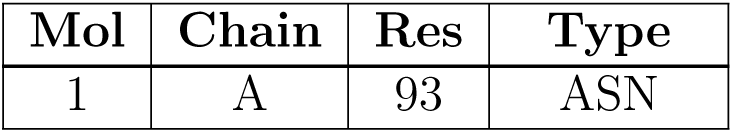

#### 5.3.3 RNA

There are no RNA molecules in this entry.

### 5.4 Non-standard residues in protein, DNA, RNA chains

There are no non-standard protein/DNA/RNA residues in this entry.

### 5.5 Carbohydrates

There are no monosaccharides in this entry.

### 5.6 Ligand geometry

Of 4 ligands modelled in this entry, 2 are monoatomic – leaving 2 for Mogul analysis.

In the following table, the Counts columns list the number of bonds (or angles) for which Mogul statistics could be retrieved, the number of bonds (or angles) that are observed in the model and the number of bonds (or angles) that are defined in the Chemical Component Dictionary. The Link column lists molecule types, if any, to which the group is linked. The Z score for a bond length (or angle) is the number of standard deviations the observed value is removed from the expected value. A bond length (or angle) with *|Z| >* 2 is considered an outlier worth inspection. RMSZ is the root-mean-square of all Z scores of the bond lengths (or angles).

| Mol | Type | Chain | Res | Link | Bond lengths |  |  | Bond angles |  |  |
| --- | --- | --- | --- | --- | --- | --- | --- | --- | --- | --- |
| | | | | | Counts | RMSZ | $\#\vert Z\vert > 2$ | Counts | RMSZ | $\#\vert Z\vert > 2$ |
| 2 | OXL | B | 401 | 3 | 5,5,5 | 1.27 | 0 | 6,6,6 | 1.59 | 2 (33%) |
| 2 | OXL | A | 401 | 3 | 5,5,5 | 1.30 | 0 | 6,6,6 | 1.65 | 2 (33%) |

In the following table, the Chirals column lists the number of chiral outliers, the number of chiral centers analysed, the number of these observed in the model and the number defined in the Chemical Component Dictionary. Similar counts are reported in the Torsion and Rings columns. ’-’ means no outliers of that kind were identified.

| Mol | Type | Chain | Res | Link | Chirals | Torsions | Rings |
| --- | --- | --- | --- | --- | --- | --- | --- |
| 2 | OXL | B | 401 | 3 | - | 4/4/4/4 | - |

| Mol | Type | Chain | Res | Link | Chirals | Torsions | Rings |
| --- | --- | --- | --- | --- | --- | --- | --- |
| 2 | OXL | A | 401 | 3 | - | 0/4/4/4 | - |

There are no bond length outliers.

All (4) bond angle outliers are listed below:

There are no bond length outliers. All (4) bond angle outliers are listed below:
| Mol | Chain | Res | Type | Atoms | Z | Observed(°) | Ideal(°) |
| --- | --- | --- | --- | --- | --- | --- | --- |
| 2 | A | 401 | OXL | O4-C2-C1 | 2.34 | 120.11 | 113.16 |
| 2 | A | 401 | OXL | O3-C1-C2 | 2.30 | 120.00 | 113.16 |
| 2 | B | 401 | OXL | O4-C2-C1 | 2.28 | 119.93 | 113.16 |
| 2 | B | 401 | OXL | O3-C1-C2 | 2.26 | 119.89 | 113.16 |

There are no chirality outliers.

All (4) torsion outliers are listed below:

There are no chirality outliers. All (4) torsion outliers are listed below:
| Mol | Chain | Res | Type | Atoms |
| --- | --- | --- | --- | --- |
| 2 | B | 401 | OXL | O1-C1-C2-O2 |
| 2 | B | 401 | OXL | O3-C1-C2-O4 |
| 2 | B | 401 | OXL | O3-C1-C2-O2 |
| 2 | B | 401 | OXL | O1-C1-C2-O4 |

There are no ring outliers.

No monomer is involved in short contacts.

The following is a two-dimensional graphical depiction of Mogul quality analysis of bond lengths, bond angles, torsion angles, and ring geometry for all instances of the Ligand of Interest. In addition, ligands with molecular weight *>* 250 and outliers as shown on the validation Tables will also be included. For torsion angles, if less then 5% of the Mogul distribution of torsion angles is within 10 degrees of the torsion angle in question, then that torsion angle is considered an outlier. Any bond that is central to one or more torsion angles identified as an outlier by Mogul will be highlighted in the graph. For rings, the root-mean-square deviation (RMSD) between the ring in question and similar rings identified by Mogul is calculated over all ring torsion angles. If the average RMSD is greater than 60 degrees and the minimal RMSD between the ring in question and any Mogul-identified rings is also greater than 60 degrees, then that ring is considered an outlier. The outliers are highlighted in purple. The color gray indicates Mogul did not find sufficient equivalents in the CSD to analyse the geometry.

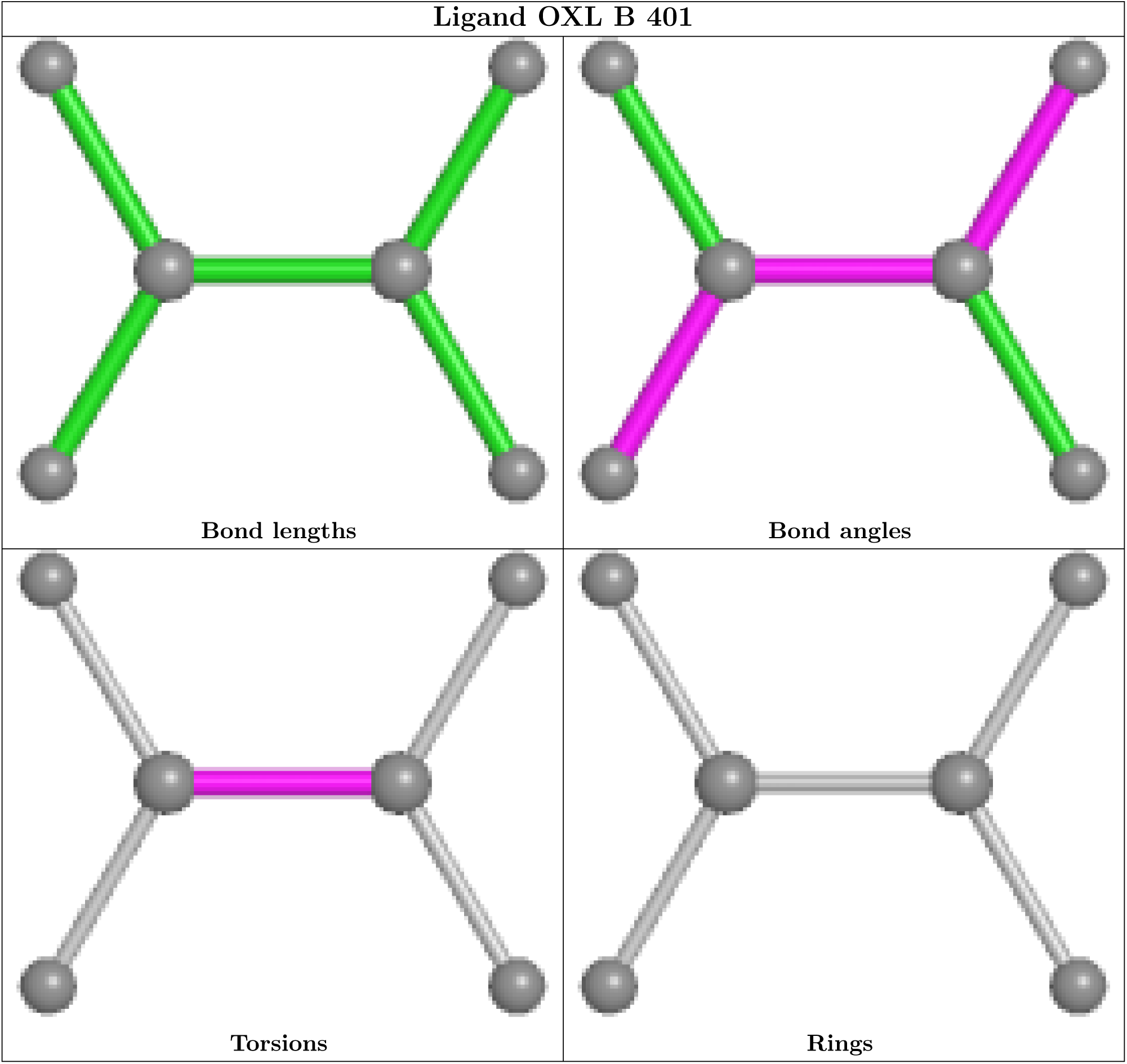

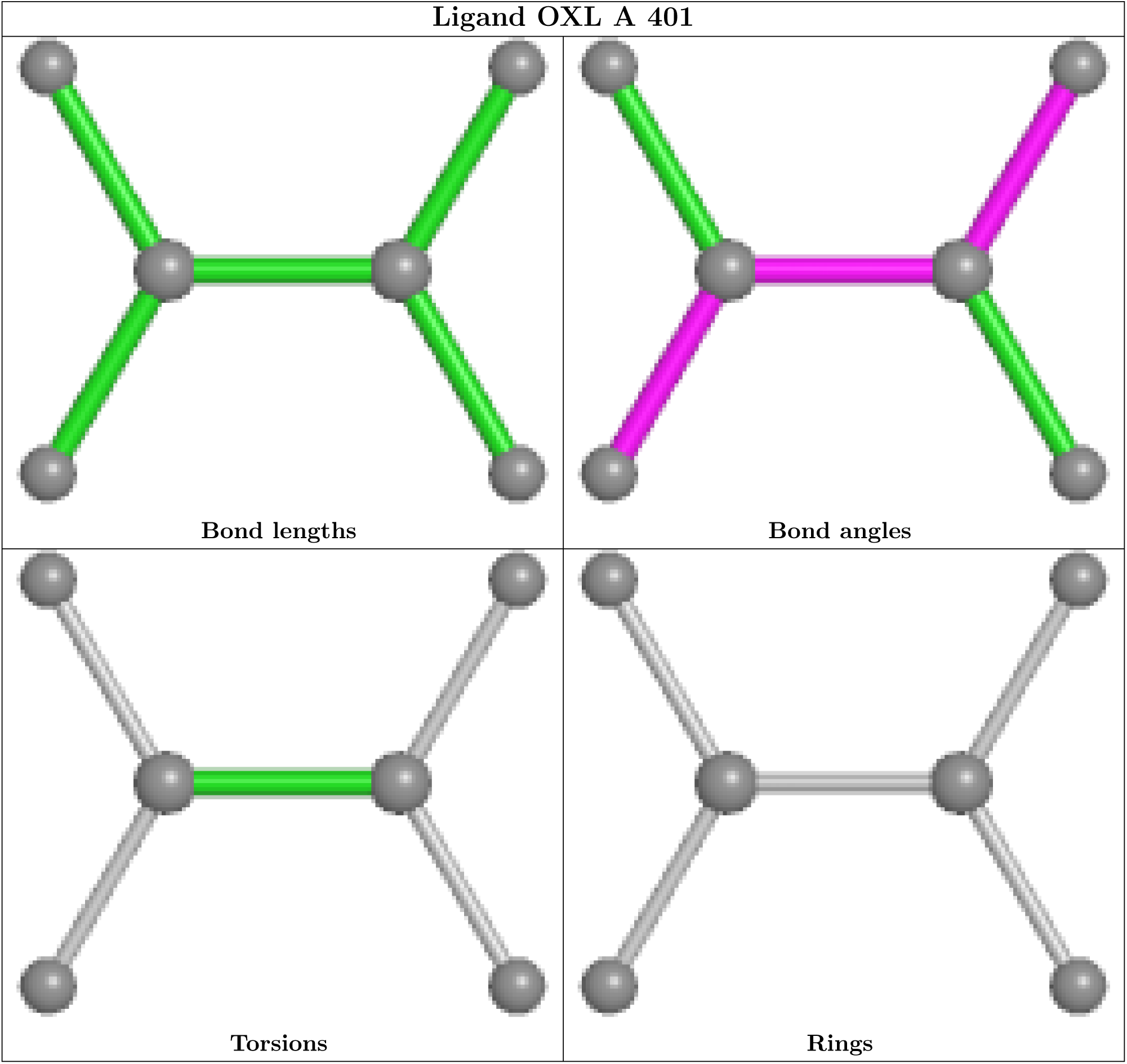

### 5.7 Other polymers

There are no such residues in this entry.

### 5.8 Polymer linkage issues

There are no chain breaks in this entry.

## 6 Fit of model and data

### 6.1 Protein, DNA and RNA chains

| Mol | Chain | Analysed | <RSRZ> | #RSRZ > 2 | OWAB(Å<sup>2</sup>) | Q < 0.9 |
| --- | --- | --- | --- | --- | --- | --- |
| 1 | A | 302/312 (96%) | -0.30 | 3 (0%) 82 80 | 31, 42, 67, 98 | 0 |
| 1 | B | 303/312 (97%) | -0.21 | 6 (1%) 65 63 | 33, 48, 78, 110 | 0 |
| All | All | 605/624 (96%) | -0.25 | 9 (1%) 73 72 | 31, 45, 74, 110 | 0 |

All (9) RSRZ outliers are listed below:

| Mol | Chain | Res | Type | RSRZ |
| --- | --- | --- | --- | --- |
| 1 | B | 68 | ASN | 5.2 |
| 1 | B | 275 | THR | 4.4 |
| 1 | B | 109 | GLY | 3.5 |
| 1 | B | 112 | ALA | 2.9 |
| 1 | B | 302 | LEU | 2.9 |
| 1 | B | 20 | ASP | 2.5 |
| 1 | A | 111 | GLU | 2.4 |
| 1 | A | 109 | GLY | 2.4 |
| 1 | A | 110 | SER | 2.3 |

### 6.2 Non-standard residues in protein, DNA, RNA chains

There are no non-standard protein/DNA/RNA residues in this entry.

### 6.3 Carbohydrates

There are no monosaccharides in this entry.

### 6.4 Ligands

| Mol | Type | Chain | Res | Atoms | RSCC | RSR | B-factors(Å<sup>2</sup>) | Q < 0.9 |
| --- | --- | --- | --- | --- | --- | --- | --- | --- |
| 2 | OXL | A | 401 | 6/6 | 0.91 | 0.17 | 43,44,52,53 | 0 |
| 2 | OXL | B | 401 | 6/6 | 0.94 | 0.14 | 50,54,55,57 | 0 |
| 3 | MN | A | 402 | 1/1 | 0.98 | 0.06 | 68,68,68,68 | 0 |
| 3 | MN | B | 402 | 1/1 | 0.98 | 0.07 | 76,76,76,76 | 0 |

The following is a graphical depiction of the model fit to experimental electron density of all instances of the Ligand of Interest. In addition, ligands with molecular weight *>* 250 and outliers as shown on the geometry validation Tables will also be included. Each fit is shown from different orientation to approximate a three-dimensional view.

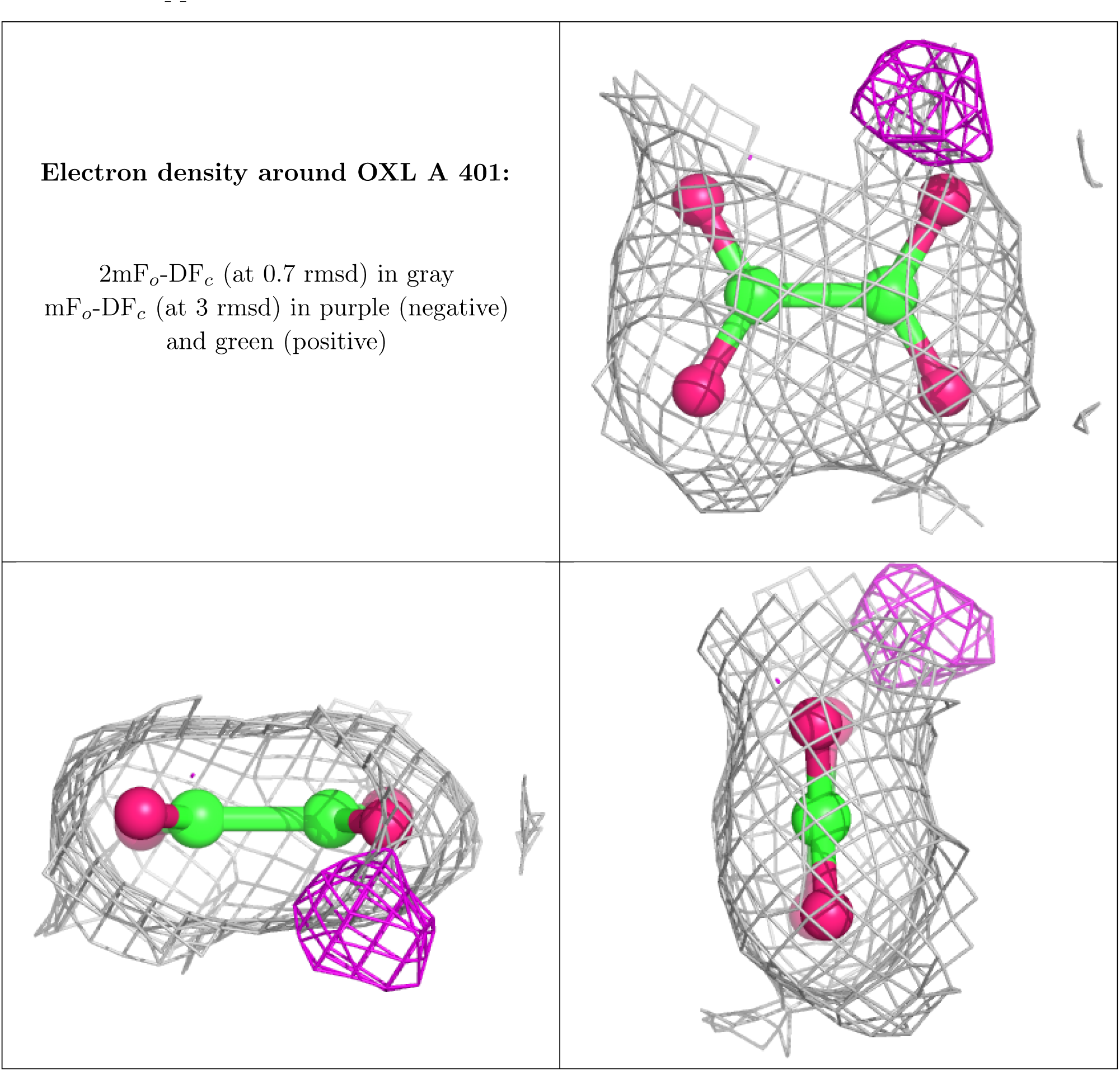

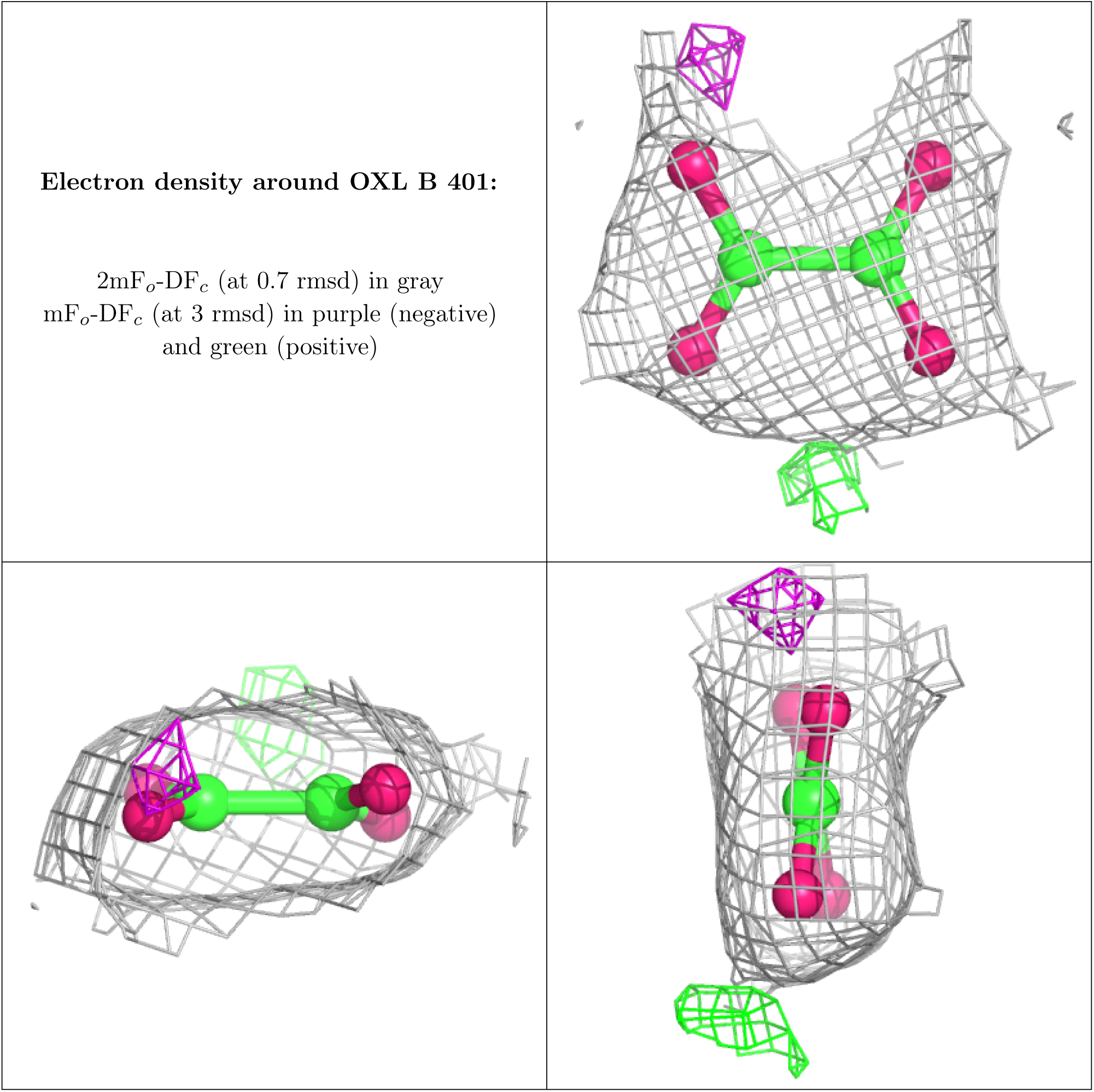

### 6.5 Other polymers

There are no such residues in this entry.

## 1 Overall quality at a glance

The following experimental techniques were used to determine the structure:

### X-RAY DIFFRACTION

The reported resolution of this entry is 1.93 Å.

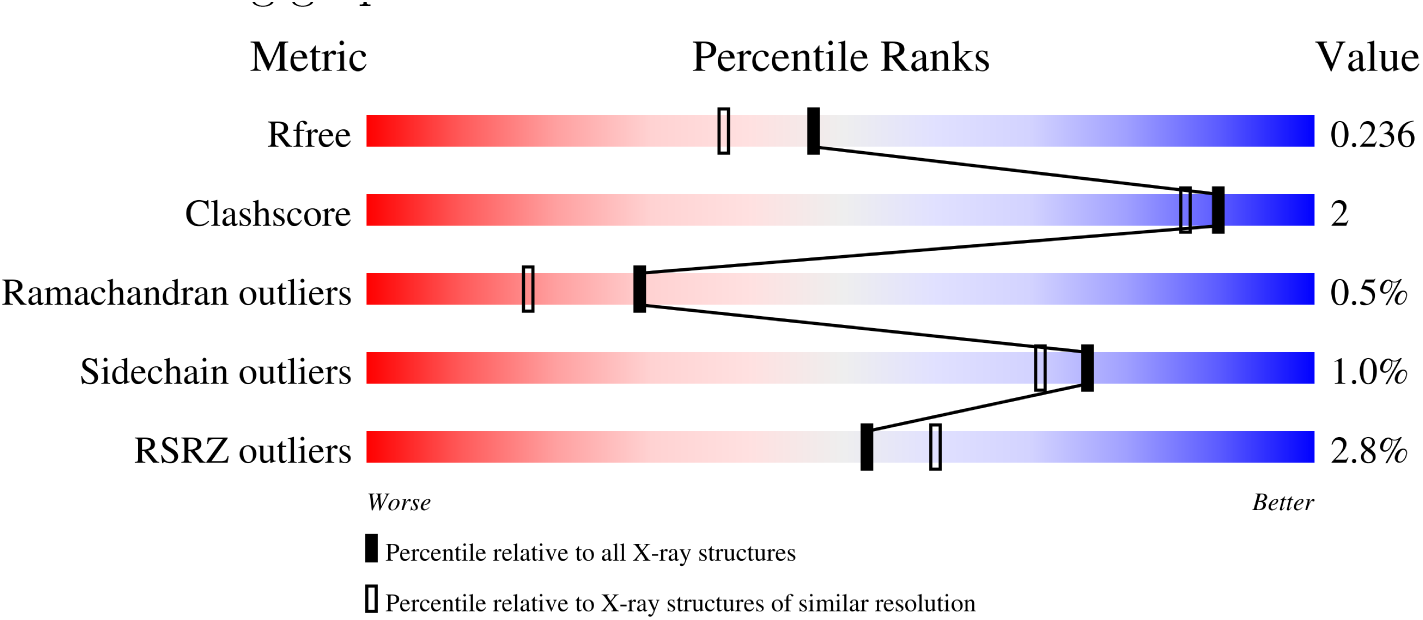

| Metric | Whole archive<br>(#Entries) | Similar resolution<br>(#Entries, resolution range(Å)) |
| --- | --- | --- |
| $R_{free}$ | 130704 | 4310 (1.96-1.92) |
| Clashscore | 141614 | 1023 (1.94-1.94) |
| Ramachandran outliers | 138981 | 1007 (1.94-1.94) |
| Sidechain outliers | 138945 | 1007 (1.94-1.94) |
| RSRZ outliers | 127900 | 4250 (1.96-1.92) |

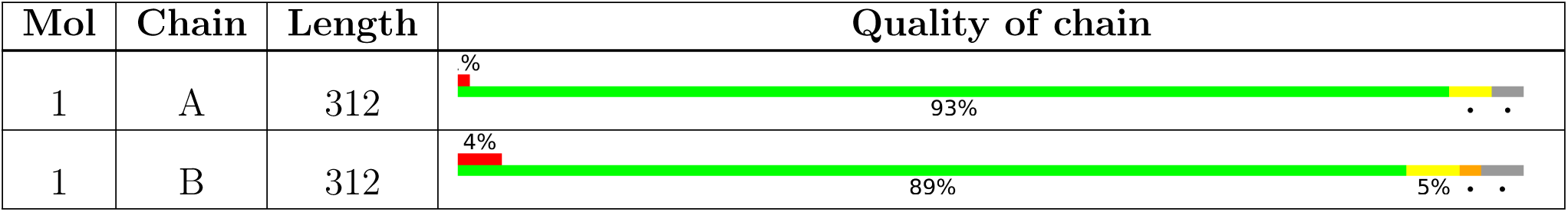

## 2 Entry composition

There are 4 unique types of molecules in this entry. The entry contains 4840 atoms, of which 0 are hydrogens and 0 are deuteriums.

- Molecule 1 is a protein called Fumarylacetoacetate hydrolase family protein.

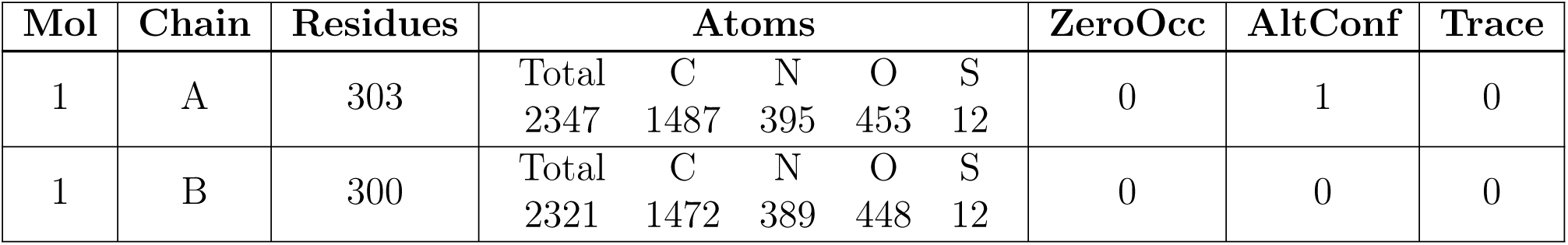

There are 22 discrepancies between the modelled and reference sequences:

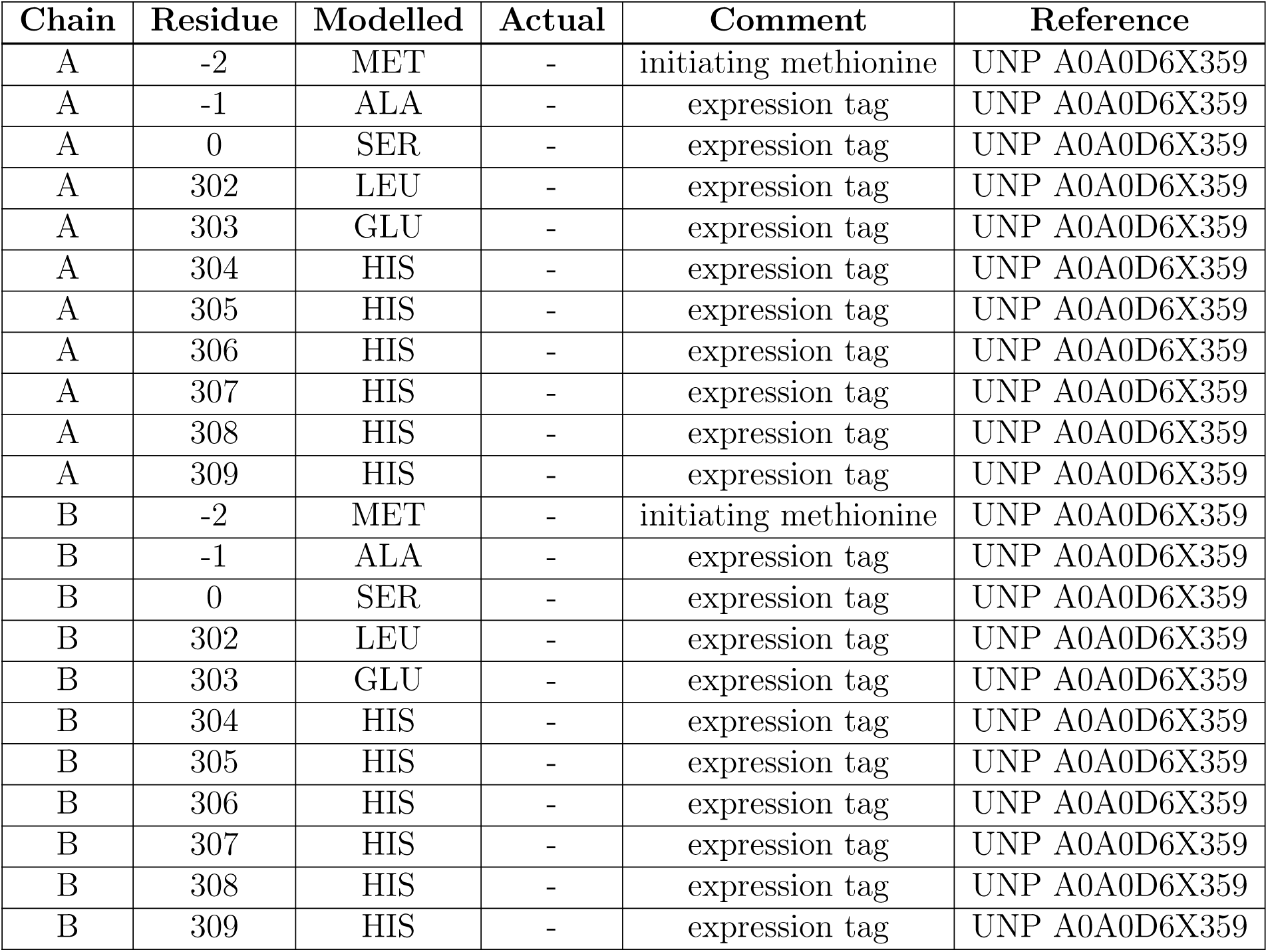

- Molecule 2 is (2 {R})-2-oxidanyl-4-oxidanylidene-pentanedioic acid (three-letter code: IO9) (formula: C_5_H_6_O_6_) (labeled as “Ligand of Interest„ by depositor).

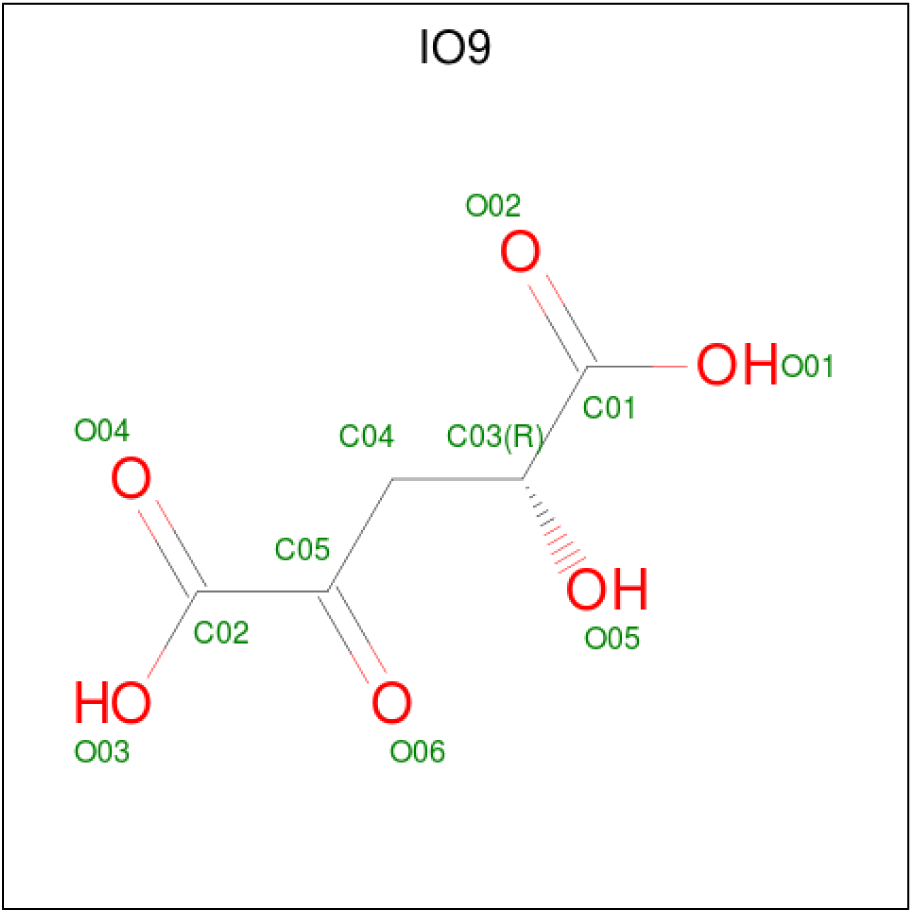

| Mol | Chain | Residues | Atoms |  |  | ZeroOcc | AltConf |
| --- | --- | --- | --- | --- | --- | --- | --- |
| 2 | A | 1 | Total | C | O | 0 | 0 |
|  |  |  | 11 | 5 | 6 |  |  |
| 2 | B | 1 | Total | C | O | 0 | 0 |
|  |  |  | 11 | 5 | 6 |  |  |

- Molecule 3 is MANGANESE (II) ION (three-letter code: MN) (formula: Mn).

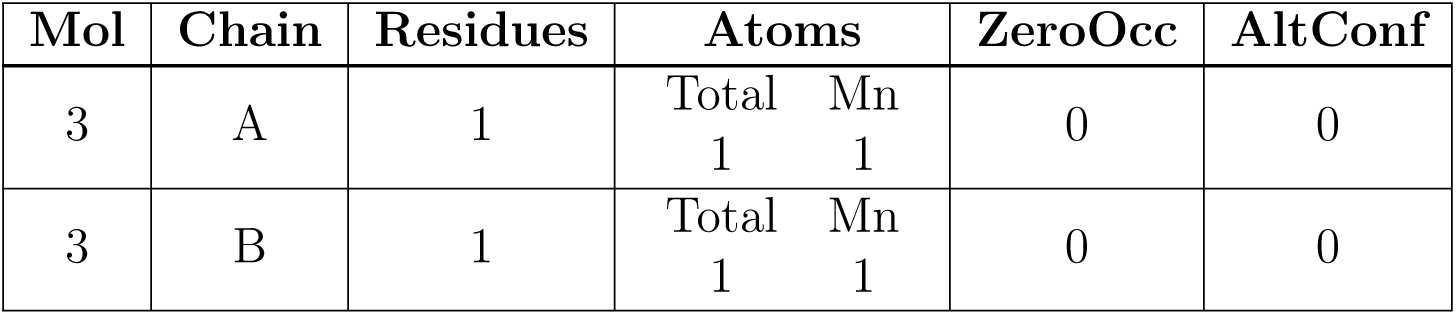

- Molecule 4 is water.

| Mol | Chain | Residues | Atoms |  | ZeroOcc | AltConf |
| --- | --- | --- | --- | --- | --- | --- |
| 4 | A | 78 | Total | O | 0 | 0 |
|  |  |  | 78 | 78 |  |  |
| 4 | B | 70 | Total | O | 0 | 0 |
|  |  |  | 70 | 70 |  |  |

## 3 Residue-property plots

*•* Molecule 1: Fumarylacetoacetate hydrolase family protein

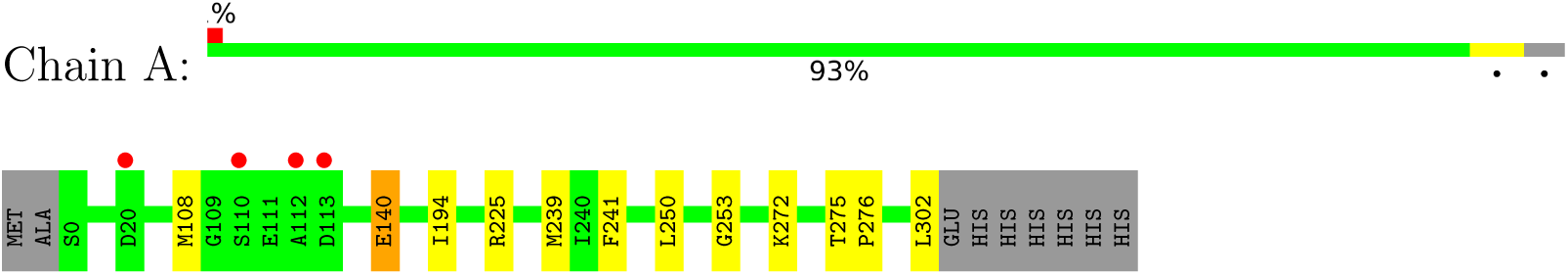

*•* Molecule 1: Fumarylacetoacetate hydrolase family protein

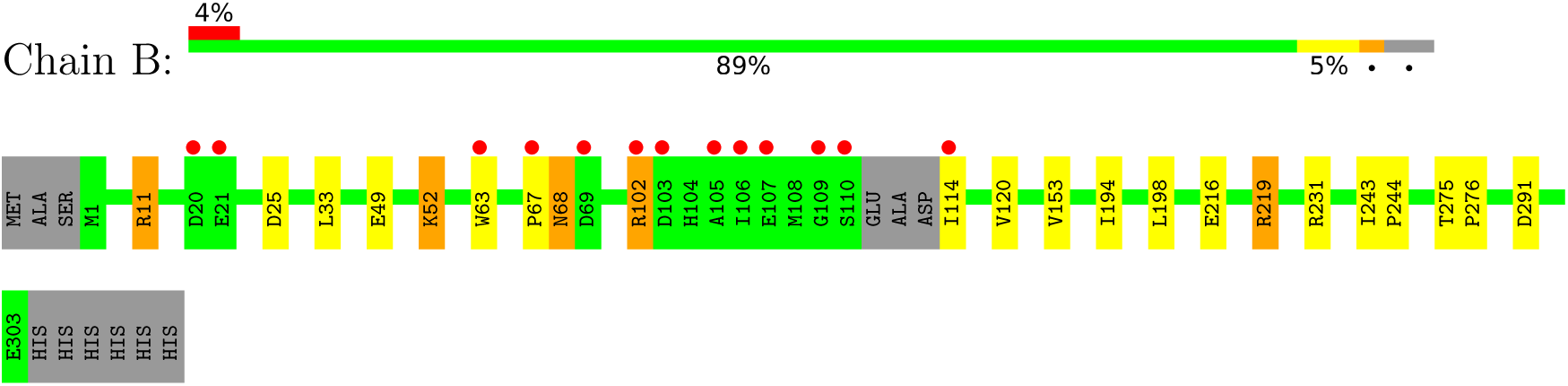

## 4 Data and refinement statistics

| Property | Value | Source |
| --- | --- | --- |
| Space group | P 21 21 21 | Depositor |
| Cell constants<br>a, b, c, $\alpha$, $\beta$, $\gamma$ | 53.69Å 93.98Å 124.98Å<br>90.00° 90.00° 90.00° | Depositor |
| Resolution (Å) | 47.04 – 1.93<br>46.99 – 1.93 | Depositor<br>EDS |
| % Data completeness<br>(in resolution range) | 97.7 (47.04-1.93)<br>97.7 (46.99-1.93) | Depositor<br>EDS |
| $R_{merge}$ | 0.07 | Depositor |
| $R_{sym}$ | (Not available) | Depositor |
| $\langle I/\sigma(I) \rangle$ <sup>1</sup> | 0.99 (at 1.92Å) | Xtriage |
| Refinement program | REFMAC 2000 | Depositor |
| R, $R_{free}$ | 0.183 , 0.230<br>0.193 , 0.236 | Depositor<br>DCC |
| $R_{free}$ test set | 1998 reflections (4.23%) | wwPDB-VP |
| Wilson B-factor (Å<sup>2</sup>) | 40.9 | Xtriage |
| Anisotropy | 0.067 | Xtriage |
| Bulk solvent $k_{sol}$(e/Å<sup>3</sup>), $B_{sol}$(Å<sup>2</sup>) | 0.31 , 33.7 | EDS |
| L-test for twinning<sup>2</sup> | $\langle \vert L\vert \rangle = 0.48$, $\langle L^2 \rangle = 0.31$ | Xtriage |
| Estimated twinning fraction | No twinning to report. | Xtriage |
| $F_o, F_c$ correlation | 0.97 | EDS |
| Total number of atoms | 4840 | wwPDB-VP |
| Average B, all atoms (Å<sup>2</sup>) | 49.0 | wwPDB-VP |

Xtriage’s analysis on translational NCS is as follows: *The largest off-origin peak in the Patterson function is 5.33% of the height of the origin peak. No significant pseudotranslation is detected*.

## 5 Model quality

### 5.1 Standard geometry

Bond lengths and bond angles in the following residue types are not validated in this section: MN, IO9

| Mol | Chain | Bond lengths |  | Bond angles |  |
| --- | --- | --- | --- | --- | --- |
|  |  | RMSZ | #\|Z\| >5 | RMSZ | #\|Z\| >5 |
| 1 | A | 0.46 | 0/2394 | 0.66 | 0/3229 |
| 1 | B | 0.44 | 0/2364 | 0.66 | 0/3187 |
| All | All | 0.45 | 0/4758 | 0.66 | 0/6416 |

| Mol | Chain | #Chirality outliers | #Planarity outliers |
| --- | --- | --- | --- |
| 1 | A | 0 | 3 |
| 1 | B | 0 | 5 |
| All | All | 0 | 8 |

There are no bond length outliers.

There are no bond angle outliers.

There are no chirality outliers.

All (8) planarity outliers are listed below:

| Mol | Chain | Res | Type | Group |
| --- | --- | --- | --- | --- |
| 1 | A | 225[A] | ARG | Sidechain |
| 1 | A | 225[B] | ARG | Sidechain |
| 1 | A | 275 | THR | Peptide |
| 1 | B | 102 | ARG | Sidechain |
| 1 | B | 11 | ARG | Sidechain |
| 1 | B | 219 | ARG | Sidechain |
| 1 | B | 231 | ARG | Sidechain |
| 1 | B | 275 | THR | Peptide |

### 5.2 Too-close contacts

| Mol | Chain | Non-H | H(model) | H(added) | Clashes | Symm-Clashes |
| --- | --- | --- | --- | --- | --- | --- |
| 1 | A | 2347 | 0 | 2373 | 7 | 0 |
| 1 | B | 2321 | 0 | 2348 | 13 | 0 |
| 2 | A | 11 | 0 | 0 | 0 | 0 |
| 2 | B | 11 | 0 | 0 | 0 | 0 |
| 3 | A | 1 | 0 | 0 | 0 | 0 |
| 3 | B | 1 | 0 | 0 | 0 | 0 |
| 4 | A | 78 | 0 | 0 | 0 | 0 |
| 4 | B | 70 | 0 | 0 | 0 | 0 |
| All | All | 4840 | 0 | 4721 | 17 | 0 |

The all-atom clashscore is defined as the number of clashes found per 1000 atoms (including hydrogen atoms). The all-atom clashscore for this structure is 2.

All (17) close contacts within the same asymmetric unit are listed below, sorted by their clash magnitude.

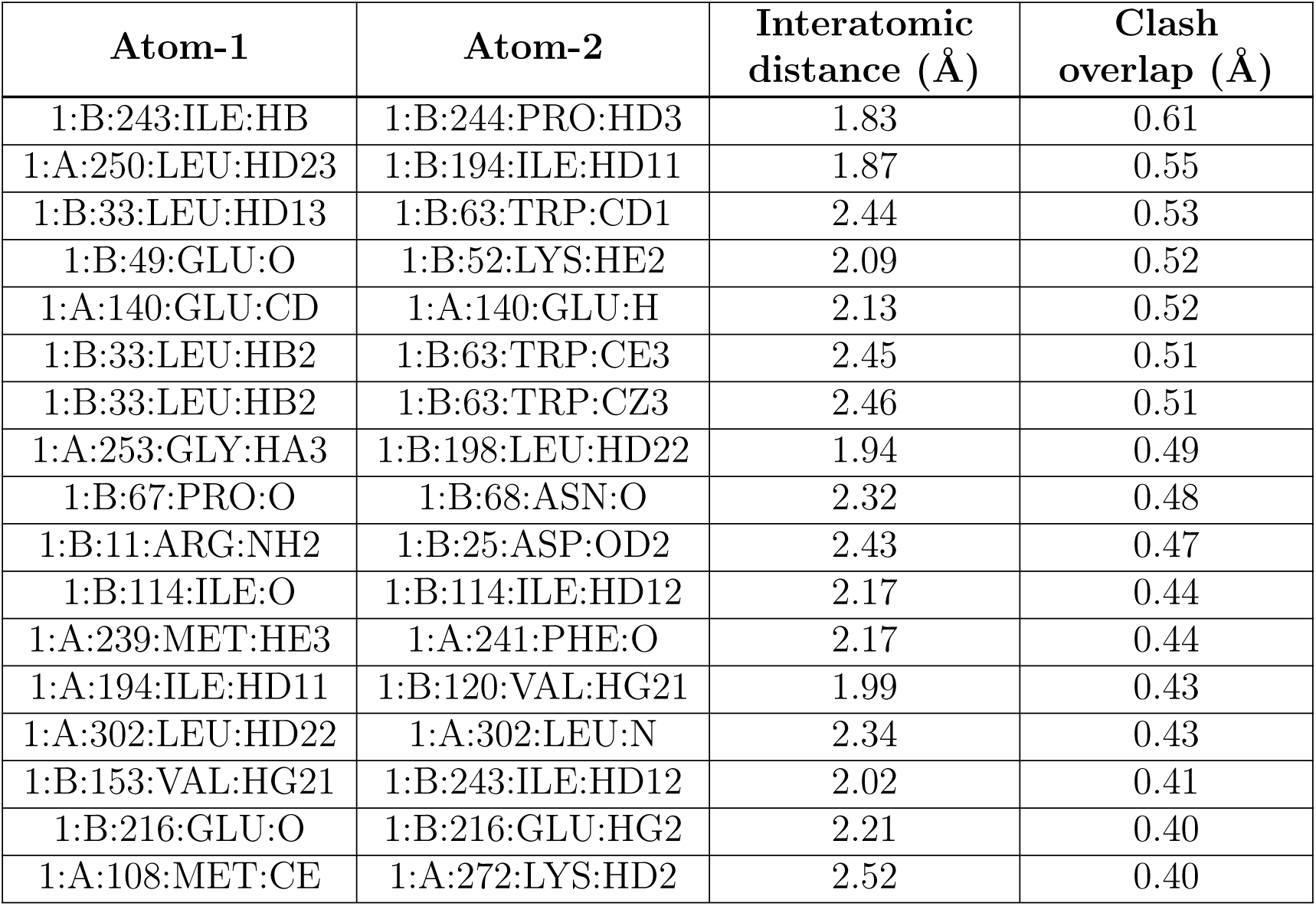

| Atom-1 | Atom-2 | Interatomic distance (Å) | Clash overlap (Å) |
| --- | --- | --- | --- |
| 1:B:243:ILE:HB | 1:B:244:PRO:HD3 | 1.83 | 0.61 |
| 1:A:250:LEU:HD23 | 1:B:194:ILE:HD11 | 1.87 | 0.55 |
| 1:B:33:LEU:HD13 | 1:B:63:TRP:CD1 | 2.44 | 0.53 |
| 1:B:49:GLU:O | 1:B:52:LYS:HE2 | 2.09 | 0.52 |
| 1:A:140:GLU:CD | 1:A:140:GLU:H | 2.13 | 0.52 |
| 1:B:33:LEU:HB2 | 1:B:63:TRP:CE3 | 2.45 | 0.51 |
| 1:B:33:LEU:HB2 | 1:B:63:TRP:CZ3 | 2.46 | 0.51 |
| 1:A:253:GLY:HA3 | 1:B:198:LEU:HD22 | 1.94 | 0.49 |
| 1:B:67:PRO:O | 1:B:68:ASN:O | 2.32 | 0.48 |
| 1:B:11:ARG:NH2 | 1:B:25:ASP:OD2 | 2.43 | 0.47 |
| 1:B:114:ILE:O | 1:B:114:ILE:HD12 | 2.17 | 0.44 |
| 1:A:239:MET:HE3 | 1:A:241:PHE:O | 2.17 | 0.44 |
| 1:A:194:ILE:HD11 | 1:B:120:VAL:HG21 | 1.99 | 0.43 |
| 1:A:302:LEU:HD22 | 1:A:302:LEU:N | 2.34 | 0.43 |
| 1:B:153:VAL:HG21 | 1:B:243:ILE:HD12 | 2.02 | 0.41 |
| 1:B:216:GLU:O | 1:B:216:GLU:HG2 | 2.21 | 0.40 |
| 1:A:108:MET:CE | 1:A:272:LYS:HD2 | 2.52 | 0.40 |

There are no symmetry-related clashes.

### 5.3 Torsion angles

#### 5.3.1 Protein backbone

| Mol | Chain | Analysed | Favoured | Allowed | Outliers | Percentiles |  |
| --- | --- | --- | --- | --- | --- | --- | --- |
| 1 | A | 302/312 (97%) | 291 (96%) | 10 (3%) | 1 (0%) | 41 | 32 |
| 1 | B | 296/312 (95%) | 285 (96%) | 9 (3%) | 2 (1%) | 22 | 11 |
| All | All | 598/624 (96%) | 576 (96%) | 19 (3%) | 3 (0%) | 29 | 17 |

All (3) Ramachandran outliers are listed below:

| Mol | Chain | Res | Type |
| --- | --- | --- | --- |
| 1 | A | 276 | PRO |
| 1 | B | 68 | ASN |
| 1 | B | 276 | PRO |

#### 5.3.2 Protein sidechains

| Mol | Chain | Analysed | Rotameric | Outliers | Percentiles |  |
| --- | --- | --- | --- | --- | --- | --- |
| 1 | A | 261/269 (97%) | 260 (100%) | 1 (0%) | 91 | 91 |
| 1 | B | 259/269 (96%) | 255 (98%) | 4 (2%) | 65 | 56 |
| All | All | 520/538 (97%) | 515 (99%) | 5 (1%) | 76 | 71 |

All (5) residues with a non-rotameric sidechain are listed below:

| Mol | Chain | Res | Type |
| --- | --- | --- | --- |
| 1 | A | 140 | GLU |
| 1 | B | 52 | LYS |
| 1 | B | 102 | ARG |

| Mol | Chain | Res | Type |
| --- | --- | --- | --- |
| 1 | B | 219 | ARG |
| 1 | B | 291 | ASP |

Sometimes sidechains can be flipped to improve hydrogen bonding and reduce clashes. All (2) such sidechains are listed below:

| Mol | Chain | Res | Type |
| --- | --- | --- | --- |
| 1 | A | 84 | HIS |
| 1 | B | 56 | HIS |

#### 5.3.3 RNA

There are no RNA molecules in this entry.

### 5.4 Non-standard residues in protein, DNA, RNA chains

There are no non-standard protein/DNA/RNA residues in this entry.

### 5.5 Carbohydrates

There are no monosaccharides in this entry.

### 5.6 Ligand geometry

Of 4 ligands modelled in this entry, 2 are monoatomic – leaving 2 for Mogul analysis.

| Mol | Type | Chain | Res | Link | Bond lengths |  |  | Bond angles |  |  |
| --- | --- | --- | --- | --- | --- | --- | --- | --- | --- | --- |
| | | | | | Counts | RMSZ | $\#\vert Z\vert > 2$ | Counts | RMSZ | $\#\vert Z\vert > 2$ |
| 2 | IO9 | B | 402 | 3 | 10,10,10 | 1.13 | 0 | 11,13,13 | 2.50 | 4 (36%) |
| 2 | IO9 | A | 401 | 3 | 10,10,10 | 1.12 | 0 | 11,13,13 | 1.98 | 4 (36%) |

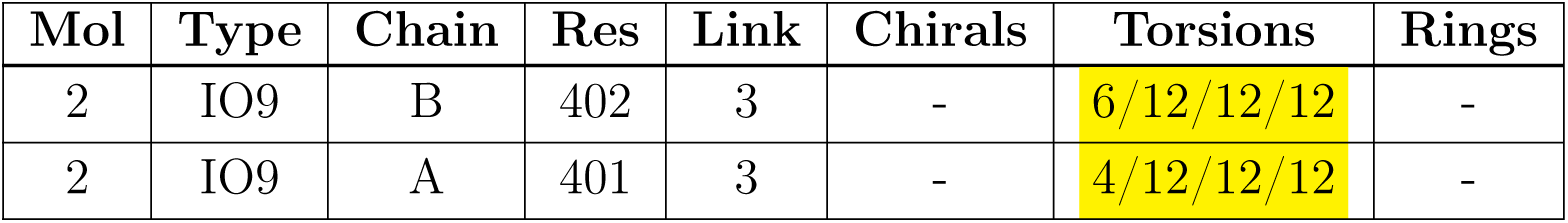

There are no bond length outliers.

All (8) bond angle outliers are listed below:

There are no bond length outliers. All (8) bond angle outliers are listed below:
| Mol | Chain | Res | Type | Atoms | Z | Observed(°) | Ideal(°) |
| --- | --- | --- | --- | --- | --- | --- | --- |
| 2 | B | 402 | IO9 | O04-C02-C05 | -4.97 | 115.08 | 121.72 |
| 2 | B | 402 | IO9 | O01-C01-C03 | 3.73 | 120.92 | 112.72 |
| 2 | A | 401 | IO9 | O03-C02-O04 | -3.27 | 116.13 | 123.61 |
| 2 | B | 402 | IO9 | O06-C05-C02 | -3.19 | 114.81 | 119.43 |
| 2 | B | 402 | IO9 | O03-C02-C05 | 3.10 | 122.46 | 113.97 |
| 2 | A | 401 | IO9 | O01-C01-C03 | 3.01 | 119.34 | 112.72 |
| 2 | A | 401 | IO9 | O06-C05-C02 | 2.81 | 123.51 | 119.43 |
| 2 | A | 401 | IO9 | O03-C02-C05 | 2.43 | 120.61 | 113.97 |

There are no chirality outliers.

All (10) torsion outliers are listed below:

There are no chirality outliers. All (10) torsion outliers are listed below:
| Mol | Chain | Res | Type | Atoms |
| --- | --- | --- | --- | --- |
| 2 | A | 401 | IO9 | C03-C04-C05-O06 |
| 2 | B | 402 | IO9 | C03-C04-C05-O06 |
| 2 | B | 402 | IO9 | O01-C01-C03-C04 |
| 2 | B | 402 | IO9 | O02-C01-C03-C04 |
| 2 | A | 401 | IO9 | C03-C04-C05-C02 |
| 2 | B | 402 | IO9 | C03-C04-C05-C02 |
| 2 | A | 401 | IO9 | O01-C01-C03-C04 |
| 2 | B | 402 | IO9 | O01-C01-C03-O05 |
| 2 | B | 402 | IO9 | O02-C01-C03-O05 |
| 2 | A | 401 | IO9 | O02-C01-C03-C04 |

There are no ring outliers.

No monomer is involved in short contacts.

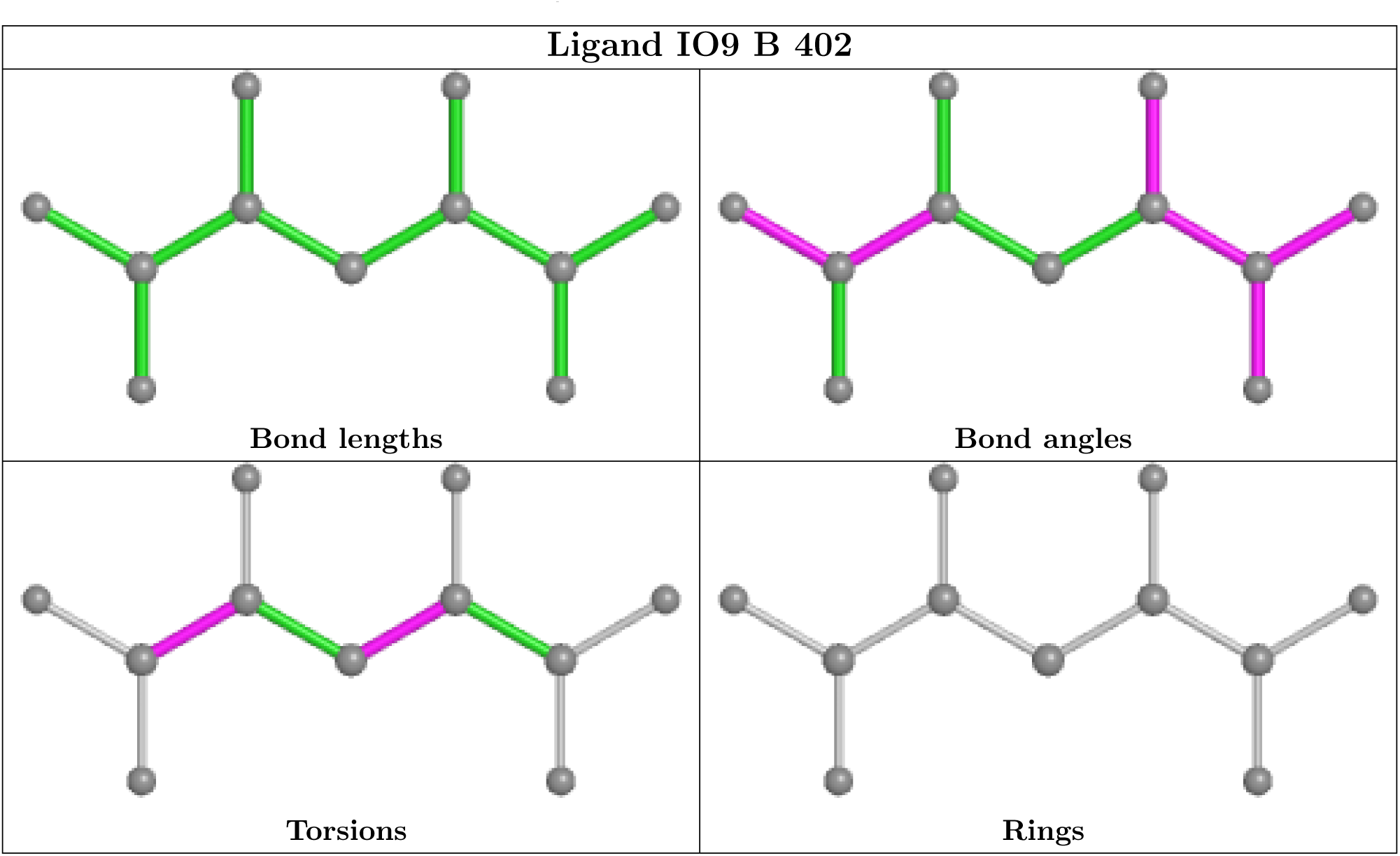

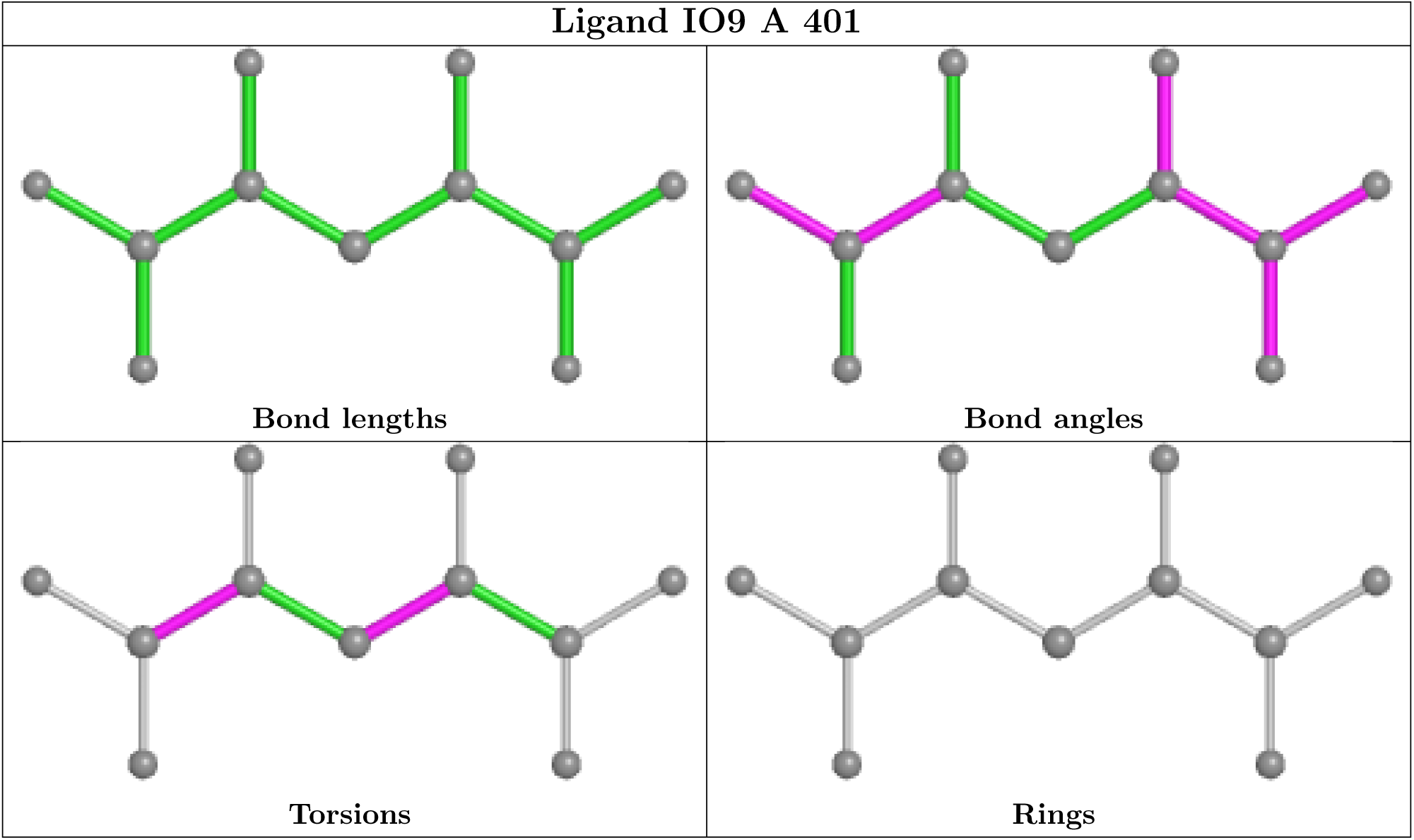

### 5.7 Other polymers

There are no such residues in this entry.

### 5.8 Polymer linkage issues

There are no chain breaks in this entry.

## 6 Fit of model and data

### 6.1 Protein, DNA and RNA chains

| Mol | Chain | Analysed | <RSRZ> | #RSRZ > 2 | OWAB(Å<sup>2</sup>) | Q < 0.9 |
| --- | --- | --- | --- | --- | --- | --- |
| 1 | A | 303/312 (97%) | -0.04 | 4 (1%) 77 81 | 30, 41, 73, 104 | 0 |
| 1 | B | 300/312 (96%) | 0.06 | 13 (4%) 35 42 | 31, 46, 87, 109 | 0 |
| All | All | 603/624 (96%) | 0.01 | 17 (2%) 53 60 | 30, 43, 81, 109 | 0 |

All (17) RSRZ outliers are listed below:

| Mol | Chain | Res | Type | RSRZ |
| --- | --- | --- | --- | --- |
| 1 | B | 69 | ASP | 4.5 |
| 1 | B | 110 | SER | 4.4 |
| 1 | A | 110 | SER | 3.3 |
| 1 | B | 67 | PRO | 2.9 |
| 1 | B | 106 | ILE | 2.9 |
| 1 | B | 102 | ARG | 2.8 |
| 1 | B | 63 | TRP | 2.4 |
| 1 | B | 109 | GLY | 2.4 |
| 1 | A | 20 | ASP | 2.4 |
| 1 | A | 113 | ASP | 2.4 |
| 1 | B | 103 | ASP | 2.4 |
| 1 | B | 20 | ASP | 2.3 |
| 1 | B | 107 | GLU | 2.3 |
| 1 | B | 21 | GLU | 2.2 |
| 1 | B | 105 | ALA | 2.2 |
| 1 | A | 112 | ALA | 2.1 |
| 1 | B | 114 | ILE | 2.0 |

### 6.2 Non-standard residues in protein, DNA, RNA chains

There are no non-standard protein/DNA/RNA residues in this entry.

### 6.3 Carbohydrates

There are no monosaccharides in this entry.

### 6.4 Ligands

| Mol | Type | Chain | Res | Atoms | RSCC | RSR | B-factors(Å<sup>2</sup>) | Q<0.9 |
| --- | --- | --- | --- | --- | --- | --- | --- | --- |
| 3 | MN | B | 401 | 1/1 | 0.78 | 0.09 | 78,78,78,78 | 0 |
| 2 | IO9 | B | 402 | 11/11 | 0.83 | 0.21 | 44,67,107,120 | 0 |
| 2 | IO9 | A | 401 | 11/11 | 0.94 | 0.15 | 36,50,80,105 | 0 |
| 3 | MN | A | 402 | 1/1 | 0.98 | 0.04 | 61,61,61,61 | 0 |

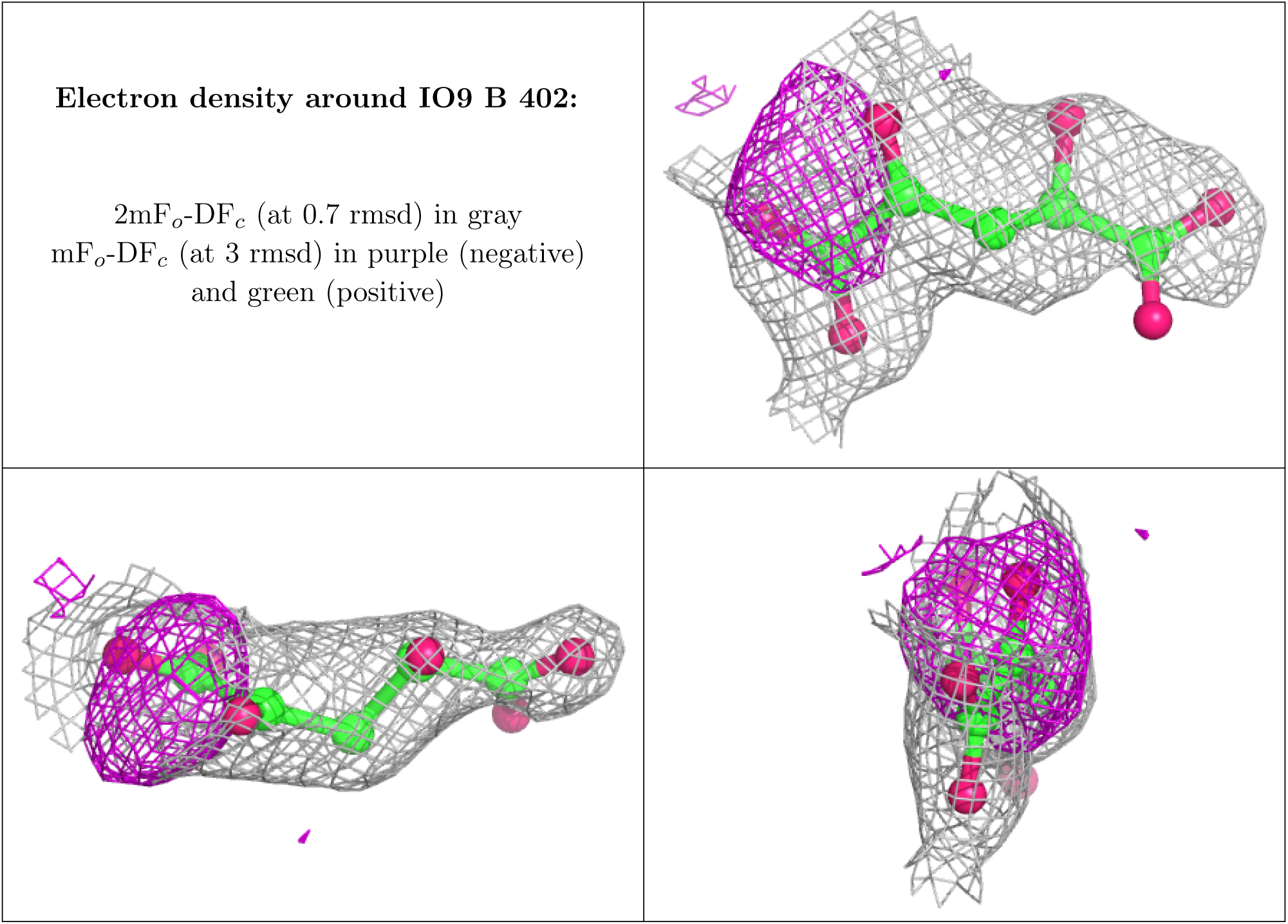

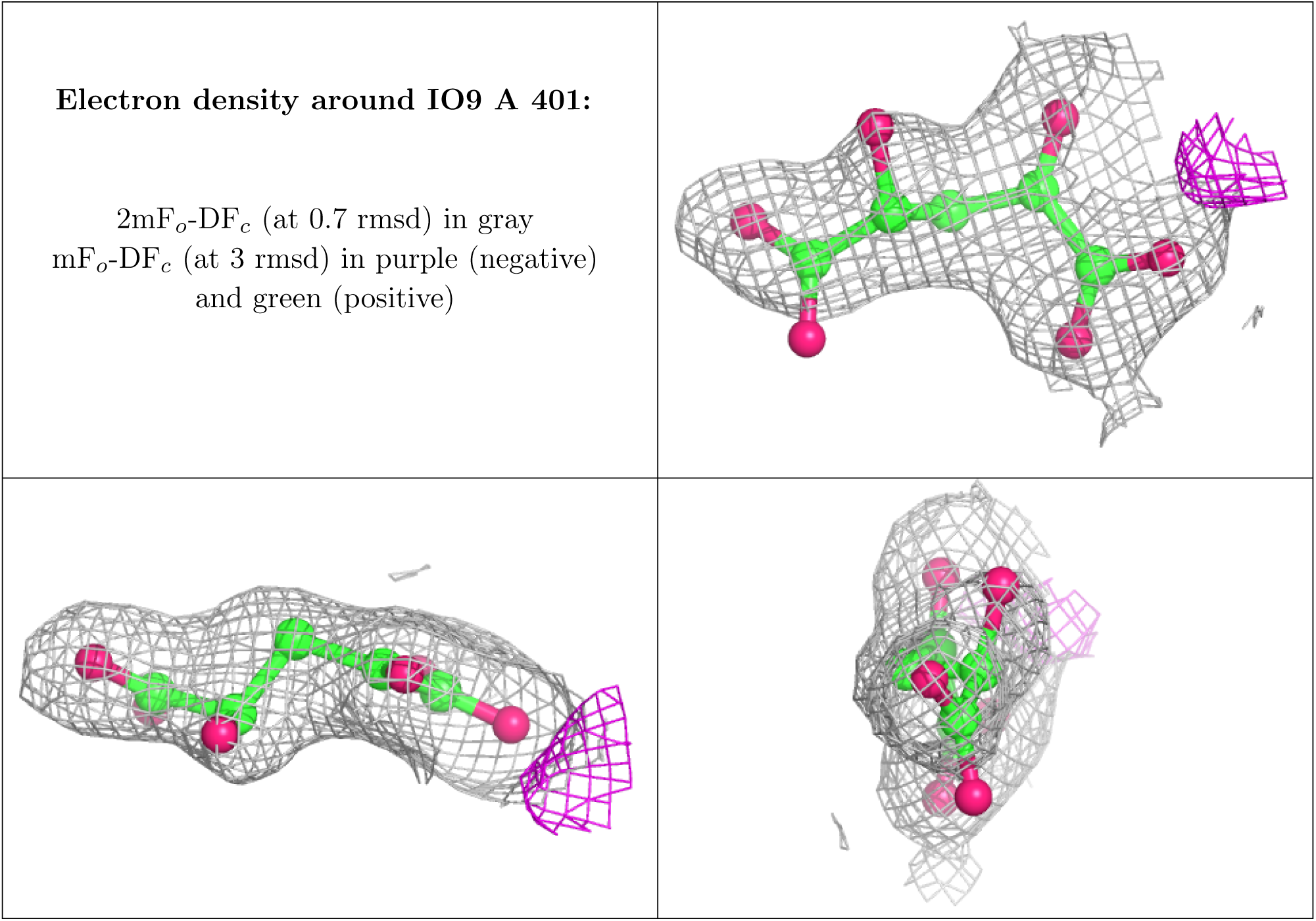

### 6.5 Other polymers

There are no such residues in this entry.

## Footnotes

1 Intensities estimated from amplitudes.

2 Theoretical values of *< |L| >*, *< L*^2^ *>* for acentric reflections are 0.5, 0.333 respectively for untwinned datasets, and 0.375, 0.2 for perfectly twinned datasets.

Validation Pipeline (wwPDB-VP) : 2.32.2

1 Intensities estimated from amplitudes.

2 Theoretical values of *< |L| >*, *< L*^2^ *>* for acentric reflections are 0.5, 0.333 respectively for untwinned datasets, and 0.375, 0.2 for perfectly twinned datasets.

Ideal geometry (DNA, RNA) : Parkinson et al. (1996)

Validation Pipeline (wwPDB-VP) : 2.32.2

1 Intensities estimated from amplitudes.

2 Theoretical values of *< |L| >*, *< L*^2^ *>* for acentric reflections are 0.5, 0.333 respectively for untwinned datasets, and 0.375, 0.2 for perfectly twinned datasets.

